# Seminal extracellular vesicles from boar AI doses contain fertility-predictive protein and miRNA cargo and improve sperm physiology

**DOI:** 10.64898/2026.03.16.712050

**Authors:** Adrián Martín-San Juan, Claudia Cerrato Martín-Hinojal, Maria José Martinez-Alborcia, Helena Nieto-Cristóbal, Eduardo de Mercado, Manuel Álvarez-Rodríguez

## Abstract

Boar semen contains spermatozoa and seminal plasma (SP) that carries extracellular vesicles (EVs) among other components. However, artificial insemination (AI) doses produced by AI companies are highly diluted based solely on sperm concentration. The aim of this study was to evaluate the integrity of EVs isolated from AI doses, characterize the protein and miRNA content from high-fertility (HF) and reduced-fertility (RF) boars, and evaluate their functional impact on spermatozoa after dilution by a coincubation up to 24 hours at 38 °C. Proteomics identified 108 differentially expressed proteins between HF and RF EVs (97 upregulated in HF, 11 in RF), and transcriptomics revealed 80 differentially expressed miRNAs (DEMs) in EVs, 52 in SP, and 3 in spermatozoa, showing inverse expression in various shared DEMs between fertility rates, suggesting compartment-specific regulation. Functional coincubation demonstrated that EVs remain biologically active after dilution. HF EVs improved sperm quality parameters and reduced oxidative stress, while RF EVs increased total and progressive motility. Overall, our findings show that EVs from AI doses retain structural integrity, carry fertility-associated protein and miRNA signatures, and functionally modulate sperm quality *in vitro*. These features highlight porcine EVs as promising biomarkers and potential tools to optimize reproductive performance in swine production.

## 1. Introduction

In recent years, reproductive procedures in the porcine industry have evolved significantly and the percentage of artificial insemination (AI) in farms have increased exponentially (M.L.W.J. Broekhuijse et al., 2012). The use of AI commercial semen doses derived from male ejaculates is well established in intensive farming systems. These ejaculates are typically diluted with maintenance extenders to preserve sperm viability during transportation to farms3/16/2026 12:39:00 PMHowever, these dilution protocols exclusively focus on adjusting sperm concentration, without considering the dilution of seminal plasma, including the extracellular vesicles (EVs) content.

This oversight is particularly relevant because seminal plasma, far from being a mere transport medium, plays a crucial role in sperm function and maturation. Upon ejaculation, immature sperm cells exit the epididymis via the vas deferens and mix with seminal plasma, a complex fluid secreted by the accessory sex glands (Maxwell et al., 2007). Seminal plasma contains a large amount of diverse organic and inorganic components including sugars, lipids, nucleic acids, proteins, hormones, salts, ions and EVs (Druart et al., 2019). Moreover, it is now well established that seminal plasma plays a pivotal role in sperm maturation by supplying key molecular factors that act as mature elements and enhance the fertilization potential of spermatozoa (Milardi et al., 2012). Among these factors, EVs which are nanoparticles, delimited by a lipid bilayer membrane that cannot replicate on their own and are released from the sexual glands cells to the seminal plasma (Alvarez-Rodriguez et al., 2019; Welsh et al., 2024), are of particular interest as they functions as novel mediators of intercellular communication (Machtinger et al., 2016).

Boar seminal plasma contains a large amount of EVs, which have been described as an heterogeneous populations (Machtinger et al., 2016). These vesicles carry a variety of bioactive molecules with significant biological implications, ranging from sperm maturation to embryo development (Rodriguez-Martinez and Roca, 2022). Their cargo is derived from the cytoplasm of the cells from which they are secreted or internalized in the course of their biogenesis (van Niel et al., 2018). Throughout the spermiogenesis, the cytoplasmic volume of spermatozoa is drastically reduced, and the replacement of histones by protamines leads to chromatin condensation, turning the genome almost inaccessible for new protein expression (Cooper, 2011; Rathke et al., 2014). At this stage, the contribution of seminal plasma components becomes critical for sperm function. Before ejaculation, as sperm transit through the epididymis, spermatozoa are already exposed to EVs secreted by the epididymal epithelial cells, known as epididymosomes (Barrachina et al., 2022). During ejaculation, additional EVs from the prostate (prostasomes) and from the seminal vesicles come into contact with the sperm and interact with them (Girouard et al., 2011; Nixon et al., 2019; Rana et al., 2024). This process is mediated by specific binding proteins such as integrins, tetraspanins, lectins or lipid mediators leading to an uptake or activation of signaling pathways (Alvarez-Rodriguez et al., 2019; van Niel et al., 2018).

The potential role of EVs in sperm maturation was first suggested in 1985, when Yanagimachi et al. observed microvesicles on the surface of sperm cells (Yanagimachi et al., 1985). Since then, numerous studies have investigated the molecular cargo of EVs across various species, aiming to understand their physiological effects on sperm during key processes such as maturation, capacitation or fertilization (Xie et al., 2022). Some reports have described the effect of these EVs to avoid premature capacitation and acrosome reaction by modulating membrane stability, particularly due to its high relation between cholesterol/phospholipid ratio (Pons-Rejraji et al., 2011). Additionally, the presence of specific proteins and micro RNAs (miRNAs) within EVs has been linked to their regulatory effects on sperm behavior. However, the direct implications of EVs in fertility status and their potential application as biomarkers in livestock production remain poorly understood and require further investigations.

The uptake of EVs by spermatozoa, leads to the internalization of their cargo, enabling the transfer of bioactive molecules like proteins, into the cells. The involvement of seminal EVs and their protein cargo in regulating sperm motility has been reported in various clinical studies, where the absence of specific proteins within these vesicles was associated with infertile diagnoses (Murdica et al., 2019; Parra et al., 2023). Notably, the influence of EVs extends beyond sperm functionality alone. Several studies have identified the presence of different antioxidant proteins inside these seminal EVs, suggesting a protective role against oxidative stress during sperm transit through the female reproductive tract (Sullivan and Saez, 2013) due to the high demand of energy.

Another key component of EV cargo is the miRNA content. These molecules are small, single-stranded non-coding RNAs, typically of 18-25 nucleotides long, that regulate gene expression by targeting messenger RNAs (mRNAs) (Ying et al., 2008). These miRNAs are encapsulated within EVs, which protect them from degradation by extracellular RNases present in the seminal plasma (Vojtech et al., 2014). The presence of various miRNAs in EVs with relevant fertility functions have been described and the transfer of these transcripts inside the cells acting as modulators is another important evidence of the EVs functional roles in modulating sperm cell activity. Certain EVs-derived miRNAs have been shown to inhibit sperm apoptosis, such as miRNA-222 (Ding et al., 2021), to modulate the female genital tract immune system to facilitate sperm tolerance and embryo implantation (Aalberts et al., 2014), and to regulate key processes in sperm development, including spermatogonial proliferation, differentiation and even traits like litter size (Dlamini et al., 2023; Xu et al., 2020).

Despite these advances, the application of seminal EVs cargo as biomarkers for fertility assessment in boar remains poorly described. In this study, we aimed to evaluate the functionality and integrity of seminal plasma EVs from highly diluted AI semen doses. Our objective was to characterize the doses EVs by confirming their integrity after the high-dilution process and establishing a reference library of proteins and miRNAs that could serve as potential fertility biomarkers for livestock industry. Additionally, we investigated the capacity of EVs derived from high-fertility (HF) and reduced-fertility (RF) samples to improve sperm quality in RF doses through a co-incubation at 38 °C for up to 24 hours using various EV concentrations from HF and RF males.

## 2. Material and methods

### 2.1. Ethics statement

The sperm samples in commercial AI semen doses were provided by the AI center AIM Ibérica, León, Spain (Topigs Norsvin Spain SLU) in accordance with European (2010–63-EU; ES13RS04P, July 2012) and Spanish regulations (ES300130640127, August 2006) on the commercialization of seminal AI-doses, ensuring compliance with animal welfare and health standards.

### 2.2. Animal handling and sperm collection and manipulation

Commercial AI doses from the landrace boar breed were used. Ejaculates were obtained using the double-glove semen collection technique, collecting the sperm-rich fraction. Samples were extended in a commercial extender from MAGAPOR (Spain) to standardize them to a final concentration of 30 x 10^6^ spermatozoa/mL in a total volume of 80 mL. Semen doses were stored at 17 °C until analysis. Each batch represented a single ejaculate obtained from breeding boars with known and compared fertility and semen quality, characterized by exceeding 80 % objective motility and having less than 15 % total sperm abnormalities, performed by the AI center.

Animal samples were selected depending on their fertility status and were classified in HF and RF samples following Broekhuijse et al. statistical analysis model (M. L. W. J. Broekhuijse et al., 2012). To classify the males, we attended to the farrowing rate and litter size values (p < 0.05), data provided by the insemination center after the analysis of more than 250 artificial inseminations per male. Animals with values > 0 were classified as HF males and animals with values < 0 as RF males.

### 2.3. Experimental design

The aim of this work was to try to get closer to the molecular explanation behind differential fertility rates in boar. Focusing on the cargo of seminal plasma EVs, we carried three different studies. *Experiment 1*. Proteome profile and differentially expressed proteins (DEPs) in seminal plasma EVs from HF (n = 6) and RF males (n =6). *Experiment 2*. miRNA transcriptome profile and differentially expressed miRNAs (DEMs) in seminal plasma EVs (n = 6 HF, 6 RF), spermatozoa (n = 6 HF, 6 RF) and free transcripts in seminal plasma (n = 6 HF, 6 RF). *Experiment* 3. Coincubation of spermatozoa pooled from four RF males with seminal EVs pooled from four HF and four RF males was performed in five independent replicates (n = 5). Coincubation was set to 38 °C for 24 hours to mimics the female genital tract conditions.

### 2.4. Isolation of seminal EVs

Seminal EVs were isolated from both HF and RF samples. For *Experiment 1* and *Experiment 2*, EVs were isolated from 20 mL of commercial AI doses from individual males. For *Experiment 3*, 60 mL of doses were obtained from a pool of four males. Diluted seminal plasma was first centrifuged at 1500 × g, 30 min to remove the sperm pellet. The supernatant was then centrifuged at 14000 × g, 10 min at 4 °C to eliminate debris and remaining membrane fragments. The resulting seminal plasma was ultrafiltrated using 20 mL PES Pierce™ Protein Concentrators (MWCO 100K, 5–20 mL volume; Thermo Scientific, US) by centrifugation at 3130 × g at 22 °C until a final volume of 1 mL was obtained. EVs were further enriched by Size Exclusion Chromatography (SEC) using an agarose resin column (Izon Science, New Zealand). 1 mL of concentrated seminal plasma was loaded onto the column, and 0.22 μm-filtered PBS 1X was used as elution buffer. The first 4 mL of elute were discarded, and fractions eluted between 4 to 7.5 mL were collected and re-concentrated using PES Pierce™ Protein Concentrators (MWCO 100K, 2–6 mL volume; Thermo Scientific) to a final volume of 0.5 mL of highly concentrated and enriched EVs. The SEC procedure was repeated three times for each fertility pool group, yielding a total volume of EVs for both HF and RF samples.

### 2.5. Seminal EVs characterization

Seminal EVs were characterized according to the Minimal Information for Studies of Extracellular Vesicles 2023 (MISEV2023) requirements (Welsh et al., 2024).

#### 2.5.1. Nanoparticle Tracking Analysis (NTA) of EVs

To determine EV concentration (EVs/mL) and size distribution (nm), samples were characterized by NTA at IdiPAZ NTA Service (Madrid, Spain). NTA measures particle size and quantity of particles in liquid suspension based on light scattering and Brownian motion principles. EV analysis was performed using a NanoSight LM10 instrument (Malvern Panalytical, Malvern, UK) equipped with a 653 nm laser. EV samples were resuspended in PBS and three 60-second videos were recorded for each sample. Particle concentration (particles/mL) and size distribution (nm) were analyzed using NanoSight software version 3.2.

#### 2.5.2. Transmission Electron Microscopy (TEM) of EVs

EV morphology and structural integrity were assessed by TEM at Centro Nacional de Biotecnología (Madrid, Spain). For negative staining, 15 μL of the sample was placed on a 400-mesh collodion-carbon coated copper grids subjected to glow discharge treatment and allowed to adsorb for 2 minutes. Grids were washed and stained with 2 % aqueous uranyl acetate solution (Electron Microscopy Sciences, US). Samples were visualized using a 100 kV JEOL JEM 1400 Flash transmission electron microscope (Jeol Ltd., Japan) equipped with a Gatan OneView digital camera (Gatan Inc., US).

#### 2.5.3. Western Blot characterization of EVs

Western blotting (WB) was performed following our previous protocol (Alvarez-Rodriguez et al., 2021) including some modifications to detect ALIX marker and to assess sample purity by measuring albumin levels. EV samples were lysed using commercial RIPA buffer (RIPA 10X, Cell Signalling Technology, US) at a final concentration of 2X + Triton 1X (Triton 100X, Sigma-Aldrich) + protease inhibitor 1X (Halt™ Protease Inhibitor Cocktail 100X, Thermofisher Scientific). Samples were sonicated with 3 pulses of 10 seconds at 60 % of amplitude (Fisherbrand FB120, Fisher Scientific) and incubated in ice 15 minutes. Protein concentration before and after extraction was determined using the DC^TM^ Bradford Protein Assay Kit (BIO-RAD Laboratories, US).

For sample purity assessment by measuring albumin levels, serial dilutions starting of a concentration of 40 μg/mL to 6.12 μg/mL were made loading a final amount of 0.25 μg to 0.00397 μg of albumin and 25 μg of protein from each sample in each well following Charette at al., 2010 methodology for protein quantification by Western blotting (Charette et al., 2010). For electrophoresis, each sample were denatured in 1X LDS sample buffer (4X LDS sample buffer, Thermofisher Scientific) and DTT for 5 min at 95 °C and loaded onto Mini-PROTEAN TGX Stain-Free gels (BIO-RAD Laboratories). Electrophoresis was conducted at 120 V for 1 hour, after which proteins were transferred to a nitrocellulose membrane (BIO-RAD Laboratories). Non-specific binding sites were blocked for 1 hour at room temperature using TBS 1X (LI-COR, US) + 5 % BSA. Membranes were incubated overnight at 4 °C with the following primary antibodies prepared in TBS 1X + 1 % BSA + 0.1 % Tween-20: anti-ALIX antibody 1:1000 (EPR15314, Abcam, UK) and anti-Pig albumin 1:1000 (A100-110A, Bethyl Laboratories Inc., US). Following three 10 min washes in TBS 1X + 0.1 % Tween-20, membranes were incubated for 1 hour at room temperature with the corresponding secondary antibodies: HRP mouse anti-Goat IgG 1:10000 (205-035-108, Jackson Immuno Research Laboratories, US) and HRP Goat anti-Rabbit IgG 1:10000 (926-80011, LI-COR), and revealed with substrate WesternSure® Premium LI-COR® chemiluminescent substrate (LI-COR) for 5 min. Signal detection was performed with the C-DIGIT Blot Scanner (LI-COR).

#### 2.5.4. Flow Cytometer characterization of EVs

Seminal doses EVs detection was performed using the flow cytometry (CytoFLEX, Beckman Coulter, US), equipped with violet (405 nm), blue (488 nm) and red (638 nm) lasers and multiple photodetectors (UV450, B525, B585, B610, R660). The side scatter (SSC) detector was configured to collect scatter from the 405-nm (violet) laser. Flow cytometry beads of known size (200 nm – 2 µm) (F13839, Thermo Fisher Scientific) were used to establish the detection threshold and to assess the cytometer sensitivity for detecting small particles. The detection threshold was set up to V-SSC > 5000 and recombinant green fluorescent protein EVs (exosome standards, fluorescent, SAE0193, Sigma-Aldrich, US) at a concentration of 5 x 10^7^ particles/mL in filtered-PBS were used to gate the EVs population. Seminal EVs were stained using CellTrace Violet (Excitation: 405 nm/ Emission: 450 nm; C34557, Invitrogen) with a concentration of 5 µM at 38 °C for 30 min and were detected with the UV450 photodetector. To immunophenotype the EVs, 25 µL of samples (aproximately 2.0 x 10^9^ EVs/mL) were incubated 30 min at 38 °C with conjugated antibodies against specific EVs markers: human anti CD63-PE diluted 1:50 (Ref. 130-118-150, Miltenyi Biotec, Germany), human anti CD81-APC 1:50 (Ref. 130-119-825, Miltenyi Biotec) and human anti Hsp70-FITC 1:50 (Ref. 130-124-700, Miltenyi Biotec). Then, 5 µL of stained EVs were diluted in 345 µL of filtered-PBS before detection. The flow rate was set to 10 µL/s and data were collected for at least 30000 particles.

To ensure that the events detected corresponded to EVs and not to artifacts or the emission of the dyes employed, we used different controls: i) filtered PBS, ii) CellTrace Violet (5 µM) without EVs (incubated 30 min at 38 °C), iii) unstained EVs, iv) Stained EVs lysated with the combination of Triton 1X (Triton 100X, Sigma-Aldrich) + 0.1% SDS (Sigma-Aldrich) incubating 30 min at 38 °C to ensure that only intact EVs were analyzed. Data analysis was performed using Kaluza Analysis Software version 2.1 (Beckman Coulter). Cytometry plots for EVs analysis are provided in **Supplementary Figure 1**.

### 2.6. Protein characterization by DDA and DIA analysis by nano-LC-MS/MS

For Experiment 1 and 2, enriched EVs from six HF and six RF animals were sent to the company BGI (BGI Genomics Co., China). A summary of the procedures performed were as follows:

#### 2.6.1. Proteolysis

Protein concentration was determined using the Bradford assay, with absorbance measured at 595 nm. For SDS-PAGE analysis, 10 μg of protein were mixed with loading buffer, heated at 95 °C for 5 min, centrifuged at 25000 × g for 5 min, and the supernatant was loaded into a 12 % SDS-polyacrylamide gel. Electrophoresis was performed at 120 V for 120 min, after which gels were stained with Coomassie Brilliant Blue for 2 hours and subsequently de-stained in 40 % ethanol + 10 % acetic acid for 3–5 cycles of 30 min each. For enzymatic digestion, 100 μg of total protein were transferred to a 10 kDa ultrafiltration tube and centrifuged for 20 min at 12000 × g at 20 °C to remove buffer components. Then, 100 μL of 0.5 M Triethylammonium bicarbonate was added, and samples were centrifuged again under the same conditions; this step was repeated three times. Trypsin was added at a 1:20 (enzyme:protein) ratio, and samples were incubated at 37 °C for 4 h, followed by centrifugation for 15 min at 12000 × g at 20 °C. The digested peptides were collected, washed with 100 μL of 0.5 M TEAB, centrifuged again, and the resulting peptide fractions were freeze-dried.

#### 2.6.2. DDA and DIA analysis by nano-LC-MS/MS

Dried peptide samples were reconstituted with a mobile phase A (100 % H2O, 0.01 % FA), centrifuged at 20000 × g for 10 min, and the supernatant was injected for LC-MS/MS. Chromatographic separation was performed on a Bruker nanoElute system. Samples were enriched on a trap column, desalted and subsequently separated on a self-packed C18 analytical column (75 μm i.d., 1.8 μm particles, 25 cm length) at a flow rate of 300 nL/min using the following gradient: 0 min, 2 % mobile phase B (100 % acetonitrile, 0.1 % formic acid); 0-45 min, mobile phase B linearly increased from 2 % to 22 %; 45-50 min, mobile phase B rose from 25 % to 35 %; 50-55 min, mobile phase B rose from 35 % to 80 %; 55-60 min, 80 % mobile phase B. The nanoliter LC separation output was directly coupled to the mass spectrometer.

##### 2.6.2.1. DDA Mass Spectrometry Detection

Peptides were ionized by nano-ESI and analyzed on a timsTOF Pro mass spectrometer operating in Data-Dependent Acquisition (DDA) mode. The main parameters were set: ion source voltage to 1.6 kV, MS1 scan range to 100∼1700 m/z; ion mobility range to 0.6-1.60 V.S/cm^2^; MS2 scan range to 100∼1700 m/z; The precursor ion screening conditions for MS2 fragmentation: charge 0 to 5+, top 10 ions with peak intensity >10000 and detectable intensity >2500. MS1 cumulative scan time to 100 ms; MS2 cumulative scan time to 100 ms. The ion fragmentation mode was CID, and the fragmentions were detected in TOF. The dynamic exclusion time was set to 30 seconds.

##### 2.6.2.2. DIA Mass Spectrometry Detection

For Data-Independent Acquisition (DIA), peptides were ionized via nano-ESI and analyzed on the timsTOF Pro with the following settings: ion source voltage set to 1.6 kV, ion mobility range to 0.6-1.60 V.S/cm^2^; MS1 mass spectrometer scanning range to 302∼1077 m/z (ions with intensity >2500 were detected); The 302∼1077 m/z range was divided into 4 steps, each divided in 8 windows (total 32 windows). The fragmentation mode was CID with an energy of 10 eV, the window width was 25 m/z, and the DIA cycle time was 3.3 seconds.

#### 2.6.3. Bioinformatics Analysis

Bioinformatic processing was performed on data generated by the high-resolution mass spectrometer. For large-scale DIA datasets, computational algorithms were used for analytical quality control and to obtain reliable quantitative values. The pipeline included Gene Ontology (GO), COG, KEGG pathway annotation, and time-series analyses. Based on the quantitative results, differentially expressed proteins (DEPs) between comparison groups were identified and functional enrichment analysis of the differential proteins were performed using ShinyGO (v. 0.85) and different databases. The databases used were: NR Database (non-redundant database from the National Center for Biotechnology Information (NCBI)), Swiss-Prot Database (from European Bioinformatics Center (EBI)), KOG/COG (Clusters of Orthologous Groups of proteins), and KEGG.

### 2.7. miRNA transcripts characterization

miRNA transcripts were sequenced not only in HF and RF seminal EVs, but also in spermatozoa and in free miRNA transcripts from seminal plasma. To separate spermatozoa from seminal plasma, 10 mL of commercial semen doses (containing a total of 300 x 10^6^ sperm cells) from six HF and six RF individual males were centrifuged for 5 min at 1500 × g at 22 °C. The supernatant (seminal plasma) was transferred to a new tube and the sperm pellet was washed with 1 mL of PBS 1X followed by centrifugation (5 min at 1500 × g at 22 °C). EVs samples were obtained as previously described.

To investigate the potential regulatory roles of miRNAs in fertility, differential expression analysis was performed for each sample type: EVs, seminal plasma, and spermatozoa comparing HF and RF groups.

The 49nt sequence obtained from sequencing were processed through adapter trimming, removal of low-quality reads and contamination removal to obtain reliable high-quality sequences for backup analysis (small RNA of 18-30nt). Quality control and filtering were performed using SOAPnuke (v1.5.6). Length distribution and shared sequence statistics were subsequently calculated. Clean reads were aligned to the reference genome using Bowtie2 (v2.2.9) for the prediction of miRNAs and other non-coding RNAs through comparison with miRBase and Rfam databases. Novel miRNAs were predicted using miRDeep2 (v. 0.1.3).

Differential miRNA expression was assessed using DESeq2 (v. 1.40.2) and clustering analysis were performed using pheatmap (v. 1.0.12). Downstream bioinformatic analyses were conducted in RStudio (v4.5.2). For novel differentially expressed miRNAs (DEMs), seed sequences (positions 2–8) were extracted, and putative human target genes were predicted using the multiMiR database (v1.32.0). For known miRNAs, target prediction was also performed. In both cases, orthologous genes in pig were identified using biomaRt (v2.66.0) and ENSEMBL orthology databases. Reproduction-related Gene Ontology (GO) biological processes were identified and categorized into development, signaling pathways, membrane regulation, oxidative stress, immune system regulation, and sperm-related functions. GO annotation and KEGG pathway enrichment (pvalueCutoff = 0.05; qvalueCutoff = 0.2) were performed using clusterProfiler (v. 4.18.2). Redundant GO terms were reduced using simplify with a semantic similarity cutoff of 0.7, retaining the most significant term based on adjusted p value.

#### 2.7.1. miRNA homology

Sequences of novel DEMs transcripts were subjected to homology analysis using BLAST (Kozomara et al., 2019) against closely related taxa. The following criteria were applied: E-Value: <10, Identity: 100 % (including same seed sequence), Query Coverage: > 90 %, Target Coverage: > 95 % and no gaps. miRNA target genes were retrieved from the miRTarBase database (Cui et al., 2025). GO enrichment analyses were performed in RStudio using the clusterProfiler package (v4.16.0), in combination with org.Hs.eg.db (v3.21.0) and enrichplot (v1.28.4).

### 2.8. Coincubation of seminal EVs with spermatozoa

Coincubation was performed using RF sperm. Five sperm pools were prepared, each composed of semen dose samples with the same spermatozoa concentration (30 × 10⁶ sperm cells/mL) from four different animals (v/ 1:1:1:1) per fertility group (n = 5). A total of 20 mL from each RF sperm pool (30 × 10⁶ sperm cells/mL) was centrifuged at 1500 × g for 5 min at 22 °C. The seminal plasma was discarded to ensure that sperm cells remained free from seminal plasma components, including endogenous EVs, prior to coincubation. The sperm pellet was resuspended in 20 mL of the maintenance extender Duragen (Magapor) and stored at 17 °C until further processing. For coincubation, 1 mL of sperm suspension was distributed into six tubes and centrifuged again (5 min, 1500 *×* g, 22 °C). After discarding the supernatant, sperm pellets were resuspended in the following treatments: i) Control (CTL): 1 mL of a 1:1 dilution of Duragen and 0.22 μm-filtered PBS 1X (EVs elution buffer); ii) 0.75 mL Duragen + 0.25 mL RF EVs (1X RF_EVs; 1.9 x 10^10^ ± 3.3 x 10^6^ EVs/mL); iii) 0.50 mL Duragen + 0.50 mL RF EVs (2X RF_EVs; 3.9 x 10^10^ ± 6.5 x 10^6^ EVs/mL); iv) 0.75 mL Duragen + 0.25 mL HF EVs (1X HF_EVs; 2.2 x 10^10^ ± 8.1 x 10^6^ EVs/mL); v): 0.50 mL Duragen + 0.50 mL HF EVs (2X HF_EVs; 4.4 x 10^10^ ± 1.6 x 10^7^ EVs/mL). Samples were incubated for up to 24 hours at 38 °C and sperm motility and physiological parameters were evaluated before incubation (0 h) and after 1, 3, 6, 12, and 24 h.

### 2.9. Sperm evaluation

#### 2.9.1. Motility and morphology anomalies analysis

For motility assessment, 5 μL of each sample was loaded on a pre-warmed Makler chamber at 37°C and analyzed using the CASA system (SPERMTECH® AI Station 1.2.24, Spain) connected to a Nikon Eclipse Si RS phase-contrast microscope at 100X magnification. A minimum of five video fields and at least 200 spermatozoa per sample were evaluated. Fifty frames per field were recorded at 50 fps. CASA settings included: minimum cell size 10 μm², maximum cell size 69 μm², 50 % straightness threshold for progressive motility, and velocity thresholds of 15 μm/s (slow), 30 μm/s (medium), and 50 μm/s (fast). A connectivity value of 12 μm was applied. The following kinematic parameters were quantified: total motility (TM, %), progressive motility (PM, %), curvilinear velocity (VCL, μm/s), straight-line velocity (VSL, μm/s), average path velocity (VAP, μm/s), linearity (LIN, %), straightness (STR, %), and wobble coefficient (WOB, %). The presence of morphology anomalies, proximal and distal abnormalities and coiled tails, was also analyzed using the CASA system.

#### 2.9.2. Physiological sperm parameters analysis

Physiological parameters related to sperm quality were analyzed by flow cytometry (CytoFLEX, Beckman Coulter, US) equipped with violet (405 nm), blue (488 nm), and red (638 nm) lasers and multiple photodetectors (UV450, B525, B585, B610, R660). Forward (FSC) and side scatter (SSC) were used to identify sperm and exclude debris, while FSC-height vs. FSC-area plots were used to select singlet events. Spermatozoa were analyzed at 400–800 cells/s, and at least 10000 sperm-like events were recorded per treatment. Staining solutions were prepared in 1X PBS from fluorochrome stock solutions at the required working concentrations. For each assay, 30 μL of sperm (30 × 10⁴ cells) were mixed with 220 μL of staining solution and incubated for 10 min at 38 °C in darkness. Hoechst 33342 (H; Ex/Em 350/461 nm; Ref. H1399, Thermo Fisher Scientific) was used in every staining at a concentration of 0.41 µM from a stock solution of 1.62 mM to identify nucleated sperm and exclude debris from the analysis. Data analysis was performed using Kaluza Analysis Software version 2.1 (Beckman Coulter). Cytometry plots for spermatozoa analysis are provided in **Supplementary Figure 2**.

##### 2.9.2.1. Sperm viability and acrosome integrity (PI/PNA-FITC)

Propidium iodide (PI; 535/617 nm) was used to assess plasma membrane integrity and fluorescein isothiocyanate-conjugated peanut (*Arachis hypogaea*) agglutinin (PNA-FITC; 579/603 nm) to assess acrosome status (Martín-San Juan et al., 2024). Working solutions of 0.37 µM PI and 2.27 µM PNA-FITC were prepared from stock solutions of PI (1.5 mM, Thermo Fisher Scientific) and PNA-FITC (9.09 mM, Thermo Fisher Scientific). Four sperm populations were distinguished: live sperm with intact acrosome (%, PI-/PNA-FITC-), live sperm with reacted acrosome (%, PI-/PNA-FITC+), dead sperm with intact acrosome (%, PI+/PNA-FITC-), and dead sperm with reacted acrosome (%, PI+/PNA-FITC+).

##### 2.9.2.2. Viability and mitochondrial activity and oxidation (YO-PRO-1/MTDR/MSox)

Mitochondrial activity was evaluated using Mitotracker Deep Red (MTDR; Ex/Em 644/665 nm), and YO-PRO-1 (491 nm/ 509 nm) was used to evaluate membrane integrity linked to sperm viability (Sargiacomo et al., 2021). Working solutions with concentrations of 0.25 µM YO-PRO-1 and 0.062 µM MTDR were prepared from stock solutions of YO-PRO-1 (1.0 mM, Thermo Fisher Scientific) and MTDR (1.0 mM, Thermo Fisher Scientific). Four sperm populations were differentiated: live spermatozoa with non-mitochondrial activity (%, YO-PRO-1-/MTDR-), live spermatozoa with mitochondrial activity (%, YO-PRO-1-/MTDR+), dead spermatozoa with non-mitochondrial activity (%, YO-PRO-1+/MTDR-), and dead spermatozoa with mitochondrial activity (%, YO-PRO-1+/MTDR+).

Mitochondrial oxidation was assessed using MitoSOX Red (MSox; 396/610 nm) in the same staining solution (Riley et al., 2021). Working solution of 0.625 µM MSox was prepared from a stock solution of 5.0 mM (Thermo Fisher Scientific). Four sperm populations were differentiated: live spermatozoa with non-mitochondrial oxidation (%, YO-PRO-1-/MSox-), live spermatozoa with mitochondrial oxidation (%, YO-PRO-1-/MSox+), dead spermatozoa with non-mitochondrial oxidation (%, YO-PRO-1+/MSox-), and dead spermatozoa with mitochondrial oxidation (%, YO-PRO-1+/MSox+).

##### 2.9.2.3. Viability and membrane fluidity (YO-PRO-1/M540)

Plasma membrane fluidity was evaluated using Merocyanine 540 (M540; 560 nm/ 578 nm) (Harrison et al., 1996), which binds to the outer cell membrane and is internalized when there is an increase of the membrane fluidity, potentially linked with capacitation status. Working solution of 0.25 µM YO-PRO-1 and 0.25 µM of M540 were prepared from stock solutions of YO-PRO-1 (1.0 mM, Thermo Fisher Scientific) and M540 (1.0 mM, Thermo Fisher Scientific). Four sperm populations were differentiated: live and non-capacitated spermatozoa (%, YO-PRO-1-/M540-), live and capacitated spermatozoa (%, YO-PRO-1-/M540+), dead and non-capacitated spermatozoa (%, YO-PRO-1+/M540-), and dead and capacitated spermatozoa (%, YO-PRO-1+/M540+).

##### 2.9.2.4. Viability and sperm oxidation (YO-PRO-1/DHE)

Cellular oxidative status was evaluated using dihydroethidium (DHE; 518/606 nm) (Martínez-Pastor et al., 2010) with YO-PRO-1. DHE reacts with cytosolic free superoxide radicals, oxidizing itself and binding to the DNA, emitting fluorescence. Working solution of 0.125 µM DHE was prepared from a stock solution of 1.0 mM (Thermo Fisher Scientific). Four sperm populations were differentiated: live and non-oxidized spermatozoa (%, H+/YO-PRO-1-/DHE-), live and oxidized spermatozoa (%, H+/YO-PRO-1-/DHE+), dead and non-oxidized spermatozoa (%, H+/YO-PRO-1+/DHE-), and dead and oxidized spermatozoa (%, H+/YO-PRO-1+/DHE+).

### 2.10. Data analysis and statistics

#### 2.10.1. Experiment 1: Proteomics

Quantitative protein measurements obtained from multiple sample replicates were analyzed using Welch’s t-test to compare the designated groups and calculate fold changes. Proteins with a fold change > 1.2 and a p-value < 0.05 were considered significantly differentially expressed. Significantly enriched GO terms and KEGG pathways of DEPs were defined as those with p < 0.05.

#### 2.10.2. Experiment 2: Transcriptomics

Statistical analyses were incorporated into the bioinformatics pipeline following miRNA sequencing. Sequence length distributions and read counts were summarized statistically to ensure data reliability. Differential expression between HF and RFgroups was analyzed using DESeq2, which applies a Wald test based on a negative binomial generalized linear model. Resulting p-values were adjusted for multiple comparisons using the Q-value method to control the False Discovery Rate (FDR). DEMs were defined using thresholds of Log2 FC ≥ 1 and Q-value < 0.05.

When biological replicates were unavailable, differential analysis was performed using a Poisson-based test following the method of Audic and Claverie. Predicted target genes of DEMs were subsequently analyzed through GO and KEGG pathway enrichment analyses. Enrichment significance was evaluated using a hypergeometric test and p-values were corrected using the Q-value method. GO terms and KEGG pathways with Q-value ≤ 0.05 were considered significantly enriched.

#### 2.10.3. Experiment 3: Coincubation

Normality and homoscedasticity of the data were tested using the Shapiro-Wilk test (p < 0.05). Non-parametric variables were transformed using Y=1/Y or by logarithmic-transformation. Two-way ANOVA was used to assess the effects of incubation time and fertility group as fixed factors, and Tukey’s test for multiple comparisons (p < 0.05). For variables that remained non-parametric after transformation, Friedman test was applied, followed by Dunn’s post-hoc multiple comparisons test (p < 0.05). Results are reported as mean ± standard error of the mean (SEM). All analyses were conducted using GraphPad Prism software v. 10.2.3 (GraphPad, USA).

## 3. Results

### 3.1. EVs characterization

Ultrastructural analysis of EV samples by TEM imaging confirmed that isolated and purified EVs from commercial AI doses showed regular cup-shaped morphology with intact membranes, without observable anomalies in either fertility group (**Figure 1A**). NTA analysis revealed a high concentration of EVs in HF samples (6.4 x 10^10^ ± 8.3 x 10^9^ VEs/mL) and in RF samples (7.8 x 10^10^ ± 5.53 x 10^9^ VEs/mL) (**Figure 1B**). Size distribution profiles were similar between groups, with no significant differences observed (HF: 251.47 ± 6.16 nm vs. RF: 244.1 ± 3.9 nm, p > 0.05) (**Figure 1C**). Soluble proteins concentrations measured after EV isolation and purification were 0.5 ± 0.2 mg/mL in HF samples and 0.4 ± 0.2 mg/mL in RF EVs (p > 0.05). Western Blot (**Figure 1D**) and flow cytometry analysis (**Supplementary Figure 1**) revealed EVs specific proteins as ALIX, CD81, CD63 and Hsp70. On the other hand, no albumin contamination were detected in EVs samples by Western blotting (**Figure 1E**).

**Figure 1.**
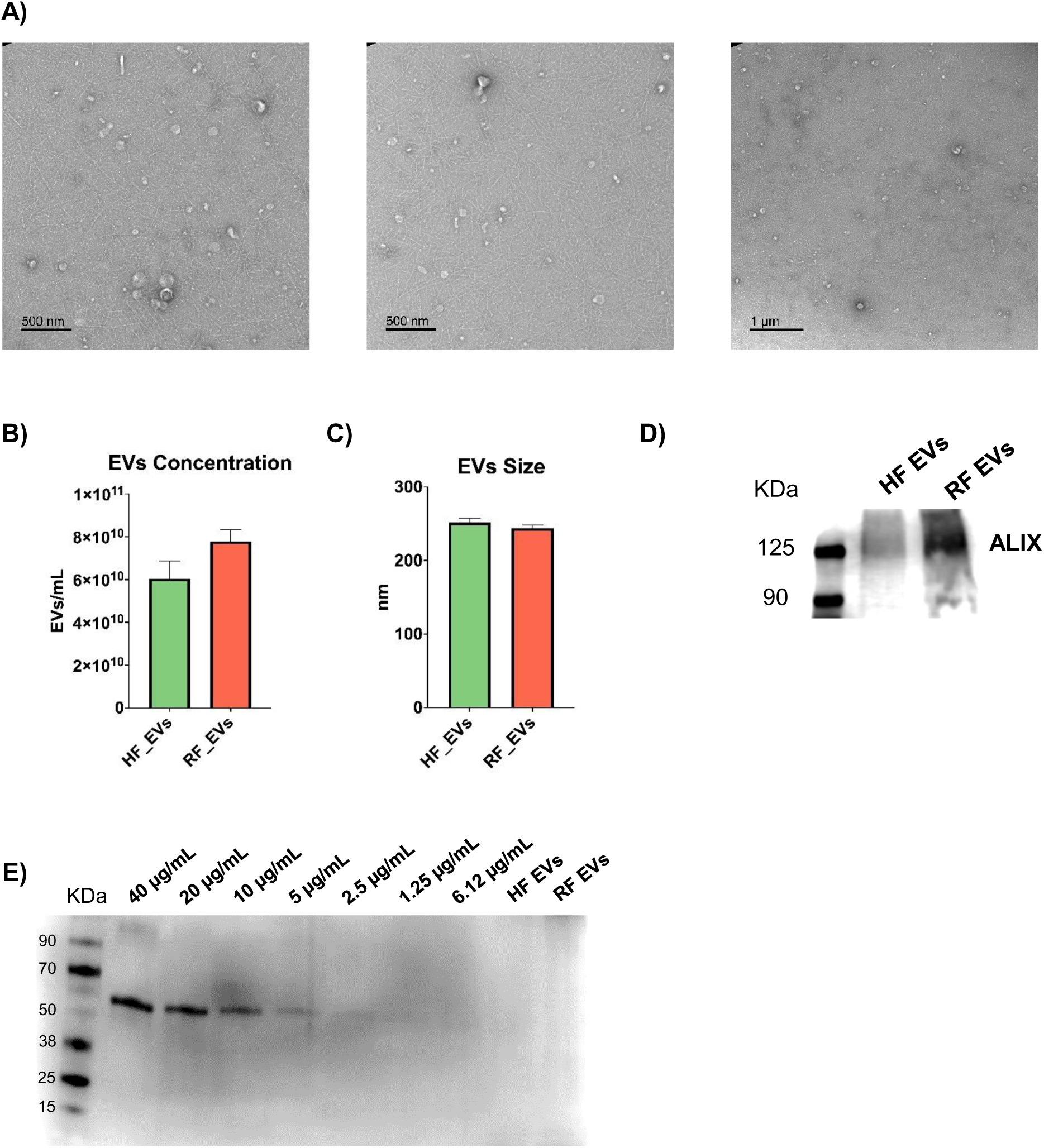
**A)** Extracellular vesicles (EVs) characterization by transmission electronic microscopy (TEM) isolated from boar seminal plasma. Scale bar: 500 nm and 1 µm. **B)** Concentration of high fertility (HF) and reduced fertility (RF) EVs in EVs/mL and **C)** Size in nm determined by Nanoparticle Tracking Analysis (NTA). **D)** Western blotting analysis testing positive for EVs protein marker ALIX in HF and RF EVs. **E)** Albumin quantification by Western blotting.

### 3.2. Protein cargo

DIA mass spectrometry detection identified a total of 10070 peptides and 1943 proteins across the twelve EV samples analyzed (HF, n = 6; RF, n = 6) (**Supplementary Table 1**). Most detected proteins (87.75 %) had a molecular weight between 1 and 100 kDa (**Supplementary Figure 3**). The distribution of unique peptides per protein showed that 72.5 % of proteins were identified with two or more unique peptides, providing high confidence in the proteomic dataset (**Supplementary Figure 4**). Established EV protein markers including CD44, CD81, CD9, ALIX, TSG101, FLOT1, FLOT2, HSP70, and HSP90 were detected. Importantly, no commonly used negative EV markers, such as Calnexin (CANX), GM130, or GRP94, were present, indicating high purity of the isolated EVs (Welsh et al., 2024).

Statistical analysis revealed 108 DEPs between HF and RF EVs (Q-value < 0.05). Among them, 97 were overexpressed in HF EVs (Log2 Fold Change > 0), whereas 11 proteins were overexpressed in RF EVs (Log2 Fold Change < 0) (**Figure 2A, 2B**). Functional annotation of DEPs was characterized by GO and KEGG enrichment analyses. In HF EVs, GO biological process enrichment highlighted pathways associated with catabolic processes, vesicle-mediated transport, and proteolysis. Four DEPs were specifically linked to sperm–egg recognition and zona pellucida binding (CCT3, CCT7, ACR, and ARSA). GO cellular component analysis indicated enrichment in proteins associated with the endomembrane system, organelle membranes, cell junctions, and vesicles. Molecular function enrichment revealed strong representation of proteins involved in ion and anion binding, hydrolase activity, and general catalytic functions (**Figure 2C**). KEGG pathway analysis identified enrichment in metabolic pathways, Rap1 signaling, biosynthesis of cofactors, and lysosome-related functions (**Figure 2D**). No GO or KEGG terms were significantly enriched (FDR < 0.05) among DEPs in RF EVs.

**Figure 2.**
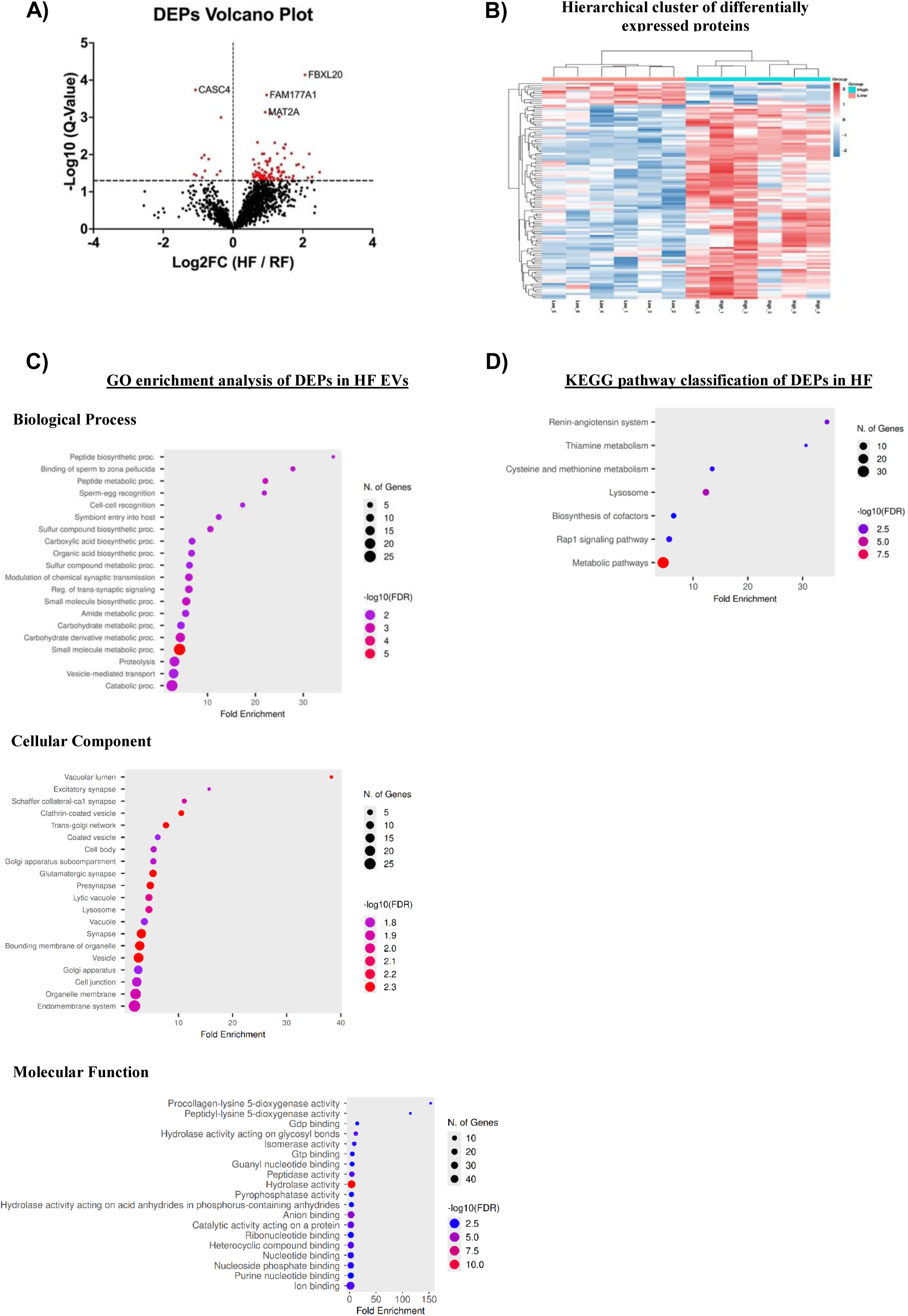
**A)** Volcano plot of differentially expressed protein (DEPs) in seminal extracellular vesicles (EVs) from high fertility (HF) and reduced fertility (RF) males. X axis: Log2 Fold Change (FC); Y axis: -Log10 (Q-Value). **B)** Heat map of DEPs. **C)** Gen Ontology (GO) enrichment analysis of DEPs in HF EVs. **D)** Kyoto Encyclopedia of Genes and Genomes (KEGG) pathway annotation of DEPs in HF EVs.

To highlight key findings, the top five most upregulated proteins in HF and RF EVs based on fold change and Q-value are listed in **Table 1**. Notably, proteins such as MMP7, ISYNA1 and NUDT2 were strongly overexpressed in HF EVs, whereas RNASE4, CASC4 and ACTB were overexpressed in RF EVs. The complete list of DEPs is provided in **Supplementary Table 2**.

**Table 1.**
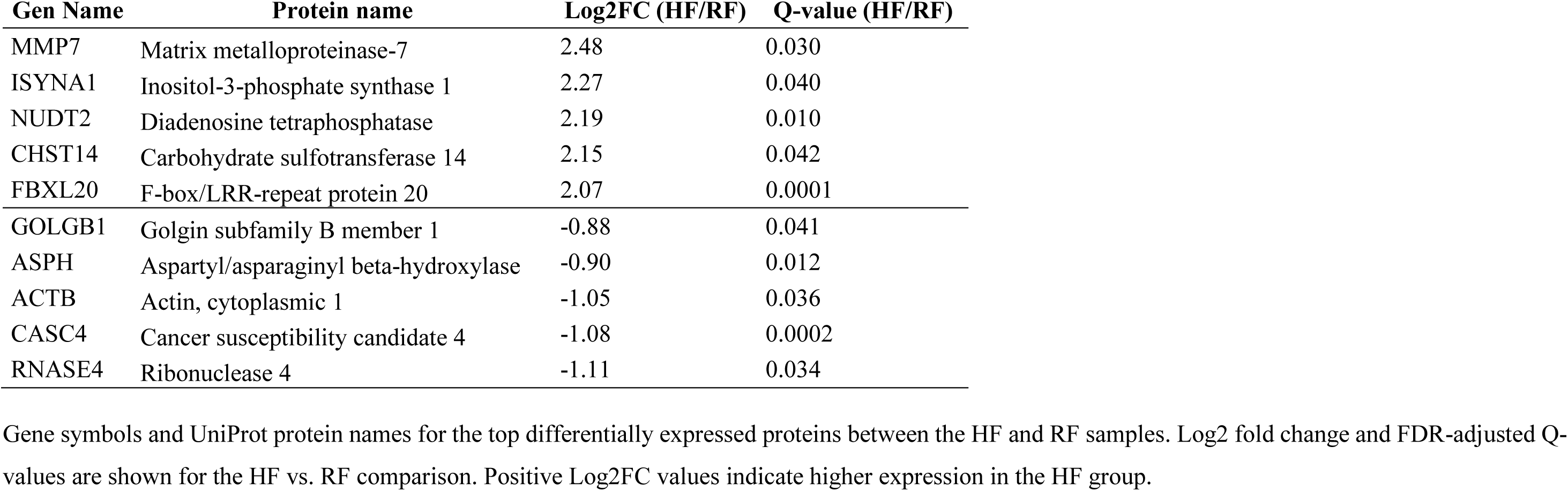
Top Differentially Expressed Proteins overexpressed in high fertility (HF) and reduced fertility (RF) extracellular vesicles.

| Gen Name | Protein name | Log2FC (HF/RF) | Q-value (HF/RF) |
| --- | --- | --- | --- |
| MMP7 | Matrix metalloproteinase-7 | 2.48 | 0.030 |
| ISYNA1 | Inositol-3-phosphate synthase 1 | 2.27 | 0.040 |
| NUDT2 | Diadenosine tetraphosphatase | 2.19 | 0.010 |
| CHST14 | Carbohydrate sulfotransferase 14 | 2.15 | 0.042 |
| FBXL20 | F-box/LRR-repeat protein 20 | 2.07 | 0.0001 |
| GOLGB1 | Golgin subfamily B member 1 | -0.88 | 0.041 |
| ASPH | Aspartyl/asparaginyl beta-hydroxylase | -0.90 | 0.012 |
| ACTB | Actin, cytoplasmic 1 | -1.05 | 0.036 |
| CASC4 | Cancer susceptibility candidate 4 | -1.08 | 0.0002 |
| RNASE4 | Ribonuclease 4 | -1.11 | 0.034 |

Gene symbols and UniProt protein names for the top differentially expressed proteins between the HF and RF samples. Log2 fold change and FDR-adjusted Q-values are shown for the HF vs. RF comparison. Positive Log2FC values indicate higher expression in the HF group.

### 3.3. miRNA cargo

Small RNA sequencing was performed on EVs, seminal plasma, and spermatozoa samples from HF and RF boars. An average of 26.7 ± 0.1 million raw reads per sample were generated, and after quality filtering, an average of 91.0 ± 0.9 % of the raw reads were retained as clean tags. Q20 values exceeded 99 %, indicating high sequencing quality. Base composition analysis of clean reads showed expected nucleotide biases, including enrichment of thymine (T) and cytosine (C) at multiple positions, especially near the 5′ and 3′ ends, without detectable contamination or sequence artifacts (low N base content). These features are consistent with known miRNA characteristics and support the overall quality of the sequencing libraries.

Mapping efficiency differed among sample types. In EVs, 55.1 ± 3.0 % of clean reads aligned to the *Sus scrofa* reference genome, while free transcripts in seminal plasma showed slightly higher alignment rates (59.1 ± 3.7 %). Spermatozoa exhibited markedly higher alignment efficiency, with 95.3 ± 0.4 % of reads mapping successfully (**Supplementary Table 3**). Differential expression analysis identified 124 DEMs across all sample types (Q-value < 0.05). EVs contained the highest number of DEMs (80), followed by seminal plasma (52), and spermatozoa (3).

#### 3.3.1. EVs miRNA cargo

EVs presented the largest number of fertility-associated miRNAs. In total, 59 miRNAs were upregulated in HF EVs (Log2 FC > 0; Q-value < 0.05), whereas 21 miRNAs were upregulated in RF EVs (Log2 FC < 0; Q-value < 0.05).

GO enrichment analysis of target genes for HF-upregulated miRNAs showed strong enrichment in biological processes associated with development, positive regulation of transcription by RNA polymerase II, and responses to stimuli. Several target genes were involved in embryo development, morphogenesis, and hormone response. Enriched cellular components included vesicles, the microtubule-organizing center, nuclear bodies, and membranes. Molecular functions such as protein kinase activity, acyltransferase activity, and calcium ion binding were also overrepresented. KEGG pathway enrichment revealed significant involvement of the PI3K–Akt, MAPK, and IgSF CAM signaling pathways. Reproduction-related GO terms included 2060 genes involved in embryo and cell development, 1653 in signaling pathways, 1588 in membrane regulation, 537 in immune system processes, 413 in oxidative stress, and 39 in sperm function. Top terms (based on Q-value) included embryonic morphogenesis, gland development, Wnt signaling, regulation of cellular response to stress, membrane lipid regulation, interleukin-12 production, and DNA damage checkpoint signaling (**Figure 3A**).

**Figure 3.**
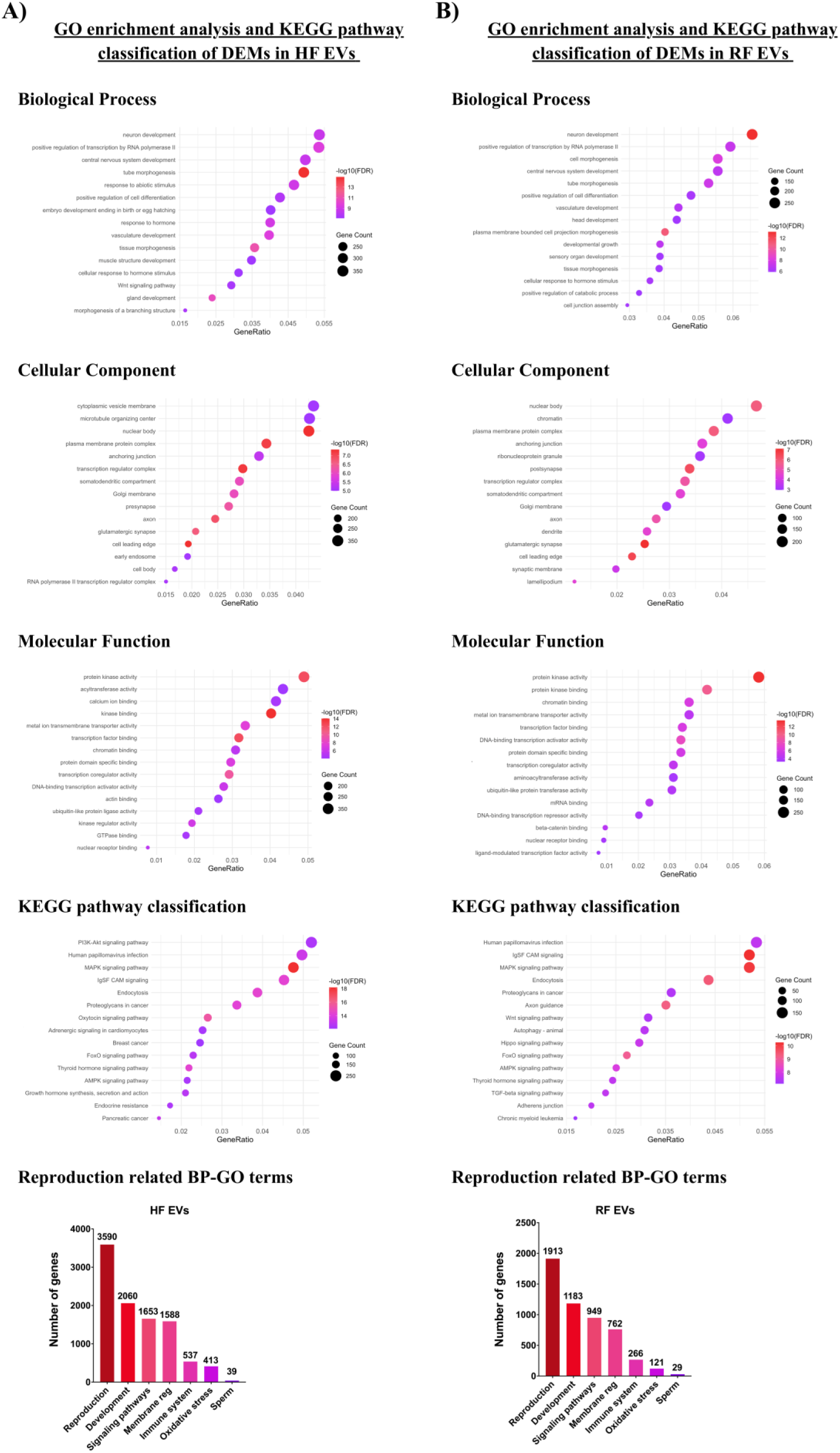
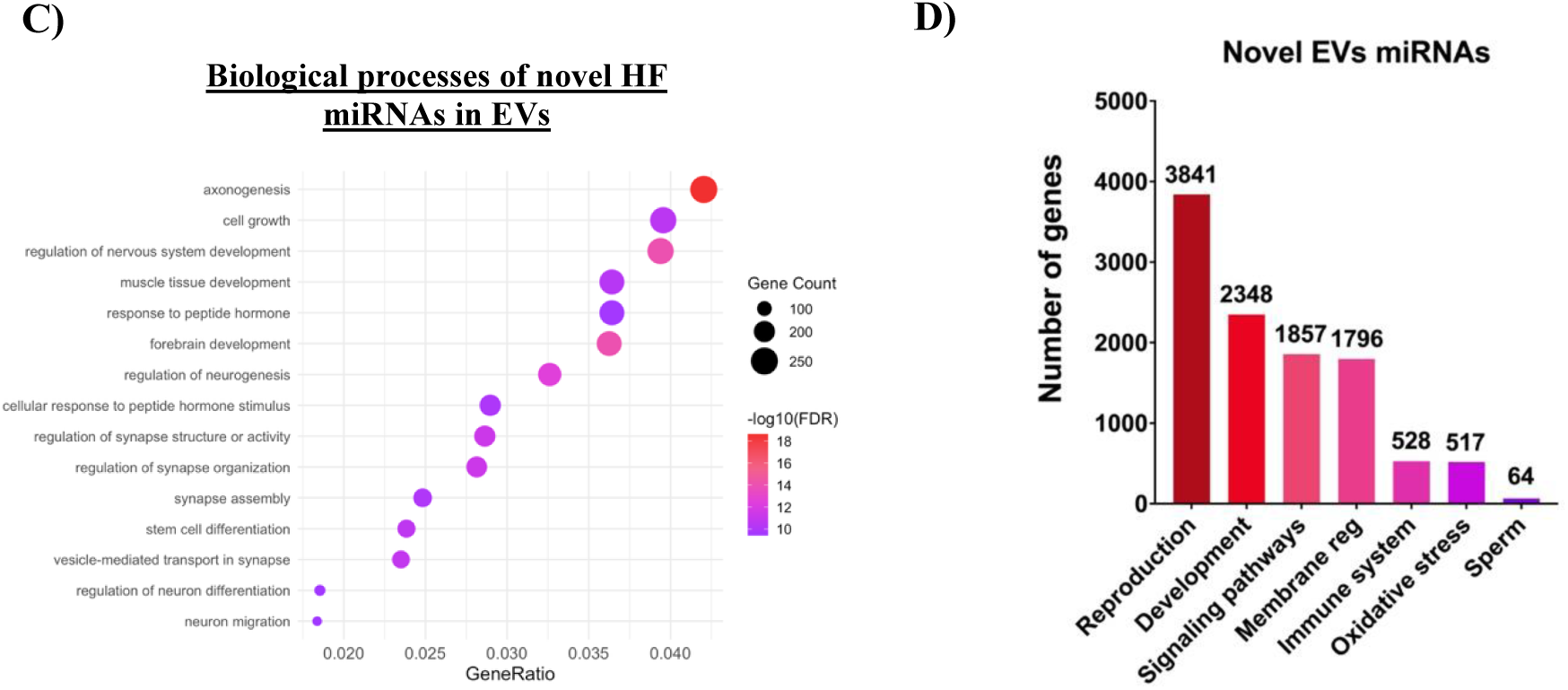
Analysis of the miRNA cargo in seminal extracellular vesicles (EVs). **A)** Gene Ontology (GO) enrichment analysis and KEGG pathway classification of the target genes of the upregulated miRNAs in high fertility (HF) EVs. **B)** GO enrichment analysis and KEGG pathway classification of the target genes of the upregulated miRNAs in reduced fertility (RF) EVs. **C)** Biological processes of GO enrichment analysis in novel *Sus scrofa* miRNAs found in EVs. **D)** Principal reproductive functions of the novel miRNAs.

Target genes of RF-upregulated EV miRNAs showed enrichment in biological processes such as neuron development, positive regulation of transcription by RNA polymerase II, and cell morphogenesis. Enriched cellular components included chromatin, nuclear bodies, and the plasma membrane, and molecular functions were dominated by protein kinase activity and chromatin binding. KEGG analysis highlighted MAPK and IgSF CAM signaling pathways. Reproduction-related categories included 1183 genes linked to development, 949 to signaling pathways, 762 to membrane regulation, 266 to immune system processes, 121 to oxidative stress, and 29 to sperm function (**Figure 3B**).

Notably, 36 DEMs in HF EVs and 14 DEMs in RF EVs corresponded to previously undescribed porcine miRNAs. 8 novel HF sequences showed high sequence homology with human miRNAs, including miR-25-3p-like, miR-203a-3p-like, miR-33a-5p-like (two occurrences), miR-188-5p-like, miR-378d-like, and miR-93-5p-like (**Table 2**). GO enrichment analysis of the novel HF miRNAs indicated that target genes were enriched for axonogenesis, cell growth, and hormone response (**Figure 3C**). Reproduction-related functions included 2348 genes involved in cell and embryo development, 1857 in signaling pathways (e.g., Wnt, PI3K/Akt, Hippo), 1796 in membrane regulation, 528 in immune regulation, 522 in oxidative stress, and 64 in sperm-related processes (e.g., androgen receptor signaling, Sertoli cell development) (**Figure 3D**).

**Table 2.**
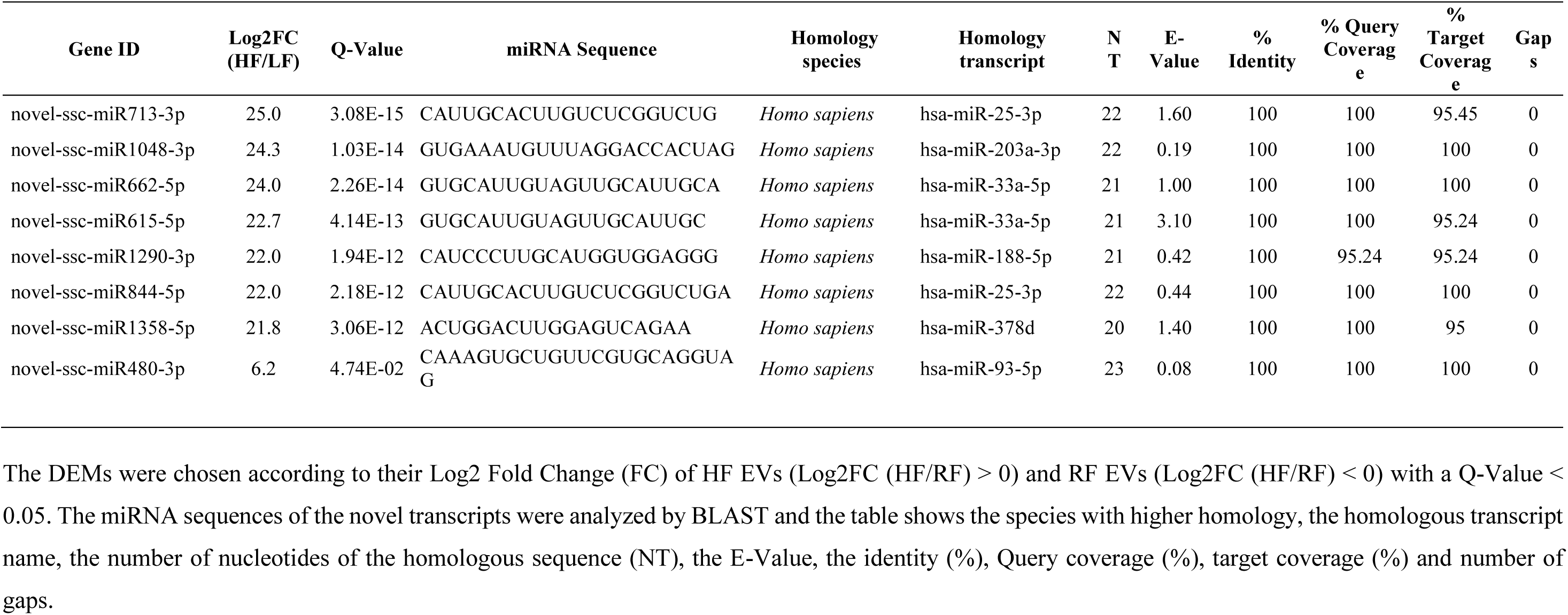
Novel differentially expressed miRNAs (DEMs) in seminal extracellular vesicles (EVs) of *Sus scrofa* by sequence homology.

Among known DEMs, miR-582-5p, miR-713-3p-like, and miR-7-3p were the most upregulated in HF EVs, while miR-3613, miR-362, and miR-23b were the most upregulated in RF EVs. The complete list is provided in **Supplementary Table 4**.

#### 3.3.2. Seminal plasma miRNA cargo

Analysis of seminal plasma revealed 33 miRNAs upregulated in HF samples (Log2 FC > 0; Q-Value < 0.05) and 19 upregulated in RF samples (Log2 FC < 0; Q-Value < 0.05). GO enrichment showed that target genes of HF-upregulated miRNAs were enriched in negative regulation of RNA biosynthesis and biogenesis, cell morphology and differentiation, and processes associated with the plasma membrane and embryonic development. Enriched cellular components included endosomes, nuclear bodies, and ribonucleoprotein granules. Molecular functions included protein kinase activity, kinase binding, and metal ion transmembrane transporter activity. KEGG enrichment identified PI3K–Akt signaling, MAPK signaling, and IgSF CAM signaling. Reproduction-related GO terms revealed 2432 target genes potentially contributing to reproductive processes, including 1423 involved in development (cell morphogenesis), 1129 in membrane regulation (response to lipid, aminoacid import or membrane docking), 1058 in signaling pathways (apoptosis, nuclear-receptor pathways), 264 in oxidative stress (autophagy, response to stress and oxidative stress), 199 in immune regulation (leukocyte and macrophage differentiation and cytokinesis), and 21 in sperm function (related with prostate) (**Figure 4A**).

**Figure 4.**
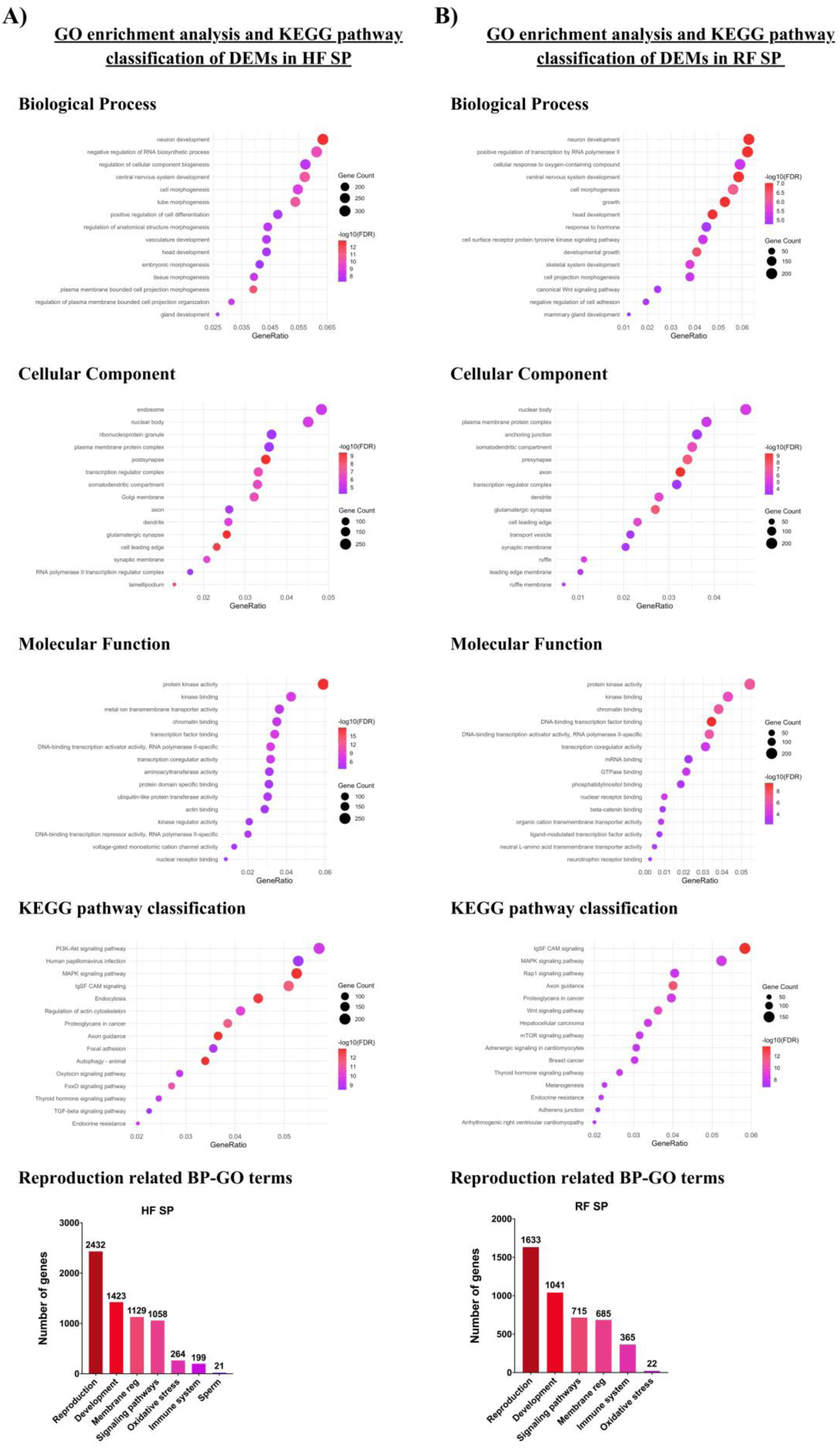
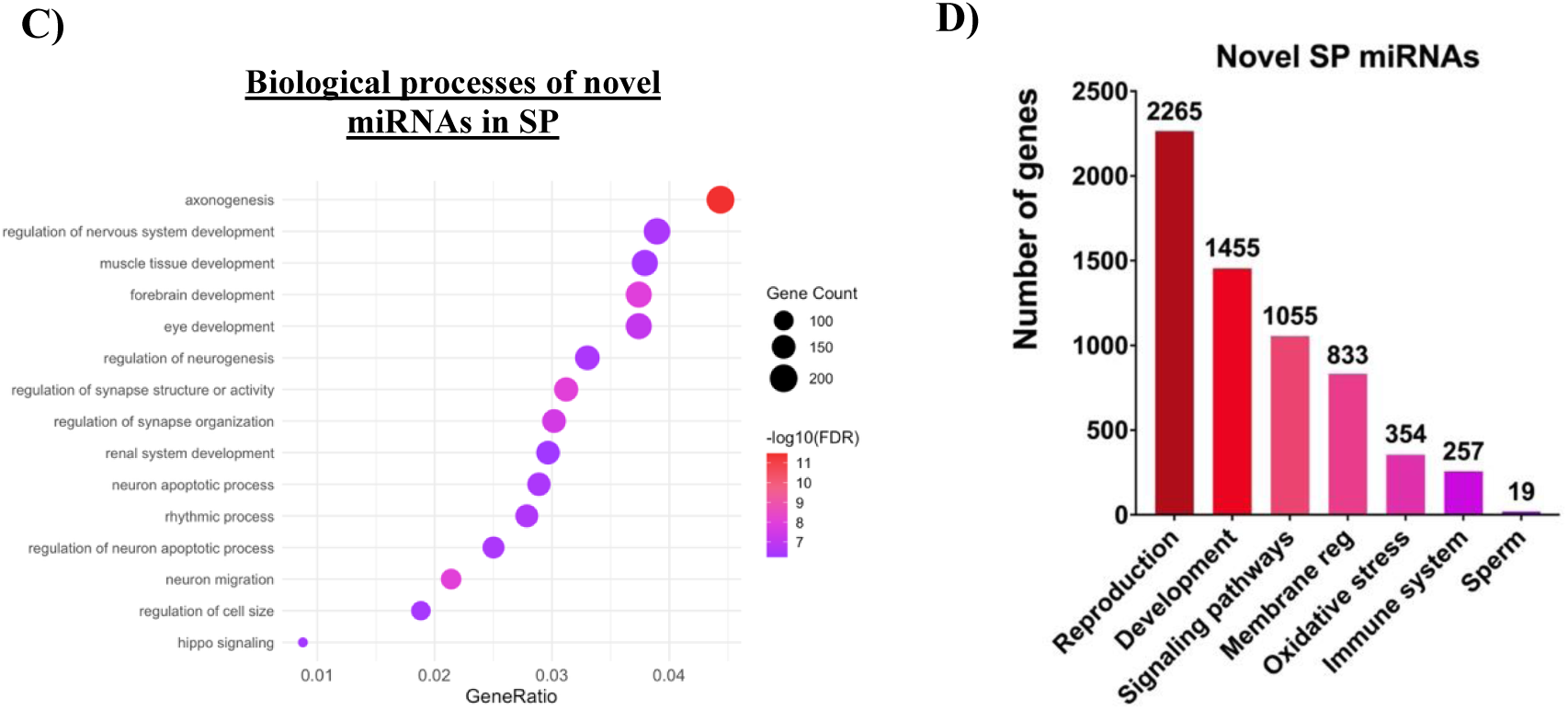
Analysis of the miRNA cargo in seminal plasma (SP). **A)** Gene Ontology (GO) enrichment analysis and KEGG pathway classification of the target genes of the upregulated miRNAs in high fertility (HF) SP. **B)** GO enrichment analysis and KEGG pathway classification of the target genes of the upregulated miRNAs in reduced fertility (RF) SP. **C)** Biological processes of GO enrichment analysis in novel *Sus scrofa* miRNAs found in SP. **D)** Principal reproductive functions of the novel miRNAs.

In RF seminal plasma, biological processes included cellular response to oxygen-containing compounds, growth, and hormone response. Enriched cellular components were the nuclear body, plasma membrane, and anchoring junction. Molecular functions with the highest representation were protein kinase activity and chromatin binding. KEGG pathways included IgSF CAM signaling, MAPK signaling, and Rap1 signaling. Reproduction-related terms included 1633 genes: 1041 in development (growth, cell morphogenesis), 715 in signaling pathways (Wnt, receptor tyrosine kinase), 685 in membrane regulation, 365 in immune processes, and 22 associated with sperm (**Figure 4B**).

Among all DEMs, 25 miRNAs in HF and 11 in RF were newly identified transcripts. Two showed high homology with human sequences: miR-93-5p-like (HF; E-value = 0.079) and miR-200a-3p-like (RF; E-value = 0.25) (**Table 3**). GO enrichment of these novel transcripts showed that target genes were related to development, regulation of cell size, and Hippo signaling (**Figure 4C**). Reproduction-related functions identified 2265 genes: 1455 linked to development, 1055 to signaling pathways, 833 to membrane regulation, 354 to oxidative stress, 257 to immune regulation, and 19 to sperm-related functions (**Figure 4D**).

**Table 3.**
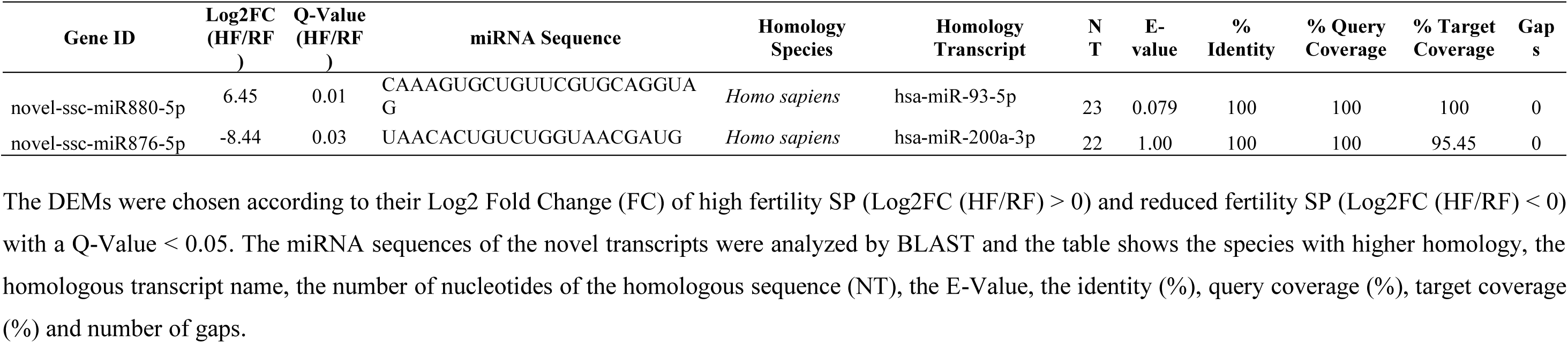
Novel differentially expressed miRNAs (DEMs) in seminal plasma (SP) of *Sus scrofa* by sequence homology.

Among all described miRNAs, the most upregulated in HF seminal plasma were miR-1249, miR-1271-5p, and miR-135, whereas miR-141, miR-423-5p, and miR-1306-3p were most upregulated in RF samples. Full results are provided in **Supplementary Table 5**.

#### 3.3.3. Spermatozoa miRNA cargo

Only spermatozoa from HF boars showed upregulated miRNAs, with three novel DEMs identified (Log2 FC > 0; Q-value < 0.05). No miRNAs were upregulated in RF spermatozoa. GO analysis of predicted target genes revealed enrichment in positive regulation of transcription, cellular growth, and secretion. Molecular functions were dominated by protein serine/threonine kinase activity and GTP binding. KEGG pathway enrichment highlighted MAPK, calcium, and Wnt signaling pathways. Reproduction-related GO terms included 148 genes: 118 associated with development and 40 with membrane regulation (**Figure 5**). None of these novel transcripts showed strong homology to known sequences in closely related species. Full results are provided in **Supplementary Table 6**.

**Figure 5.**
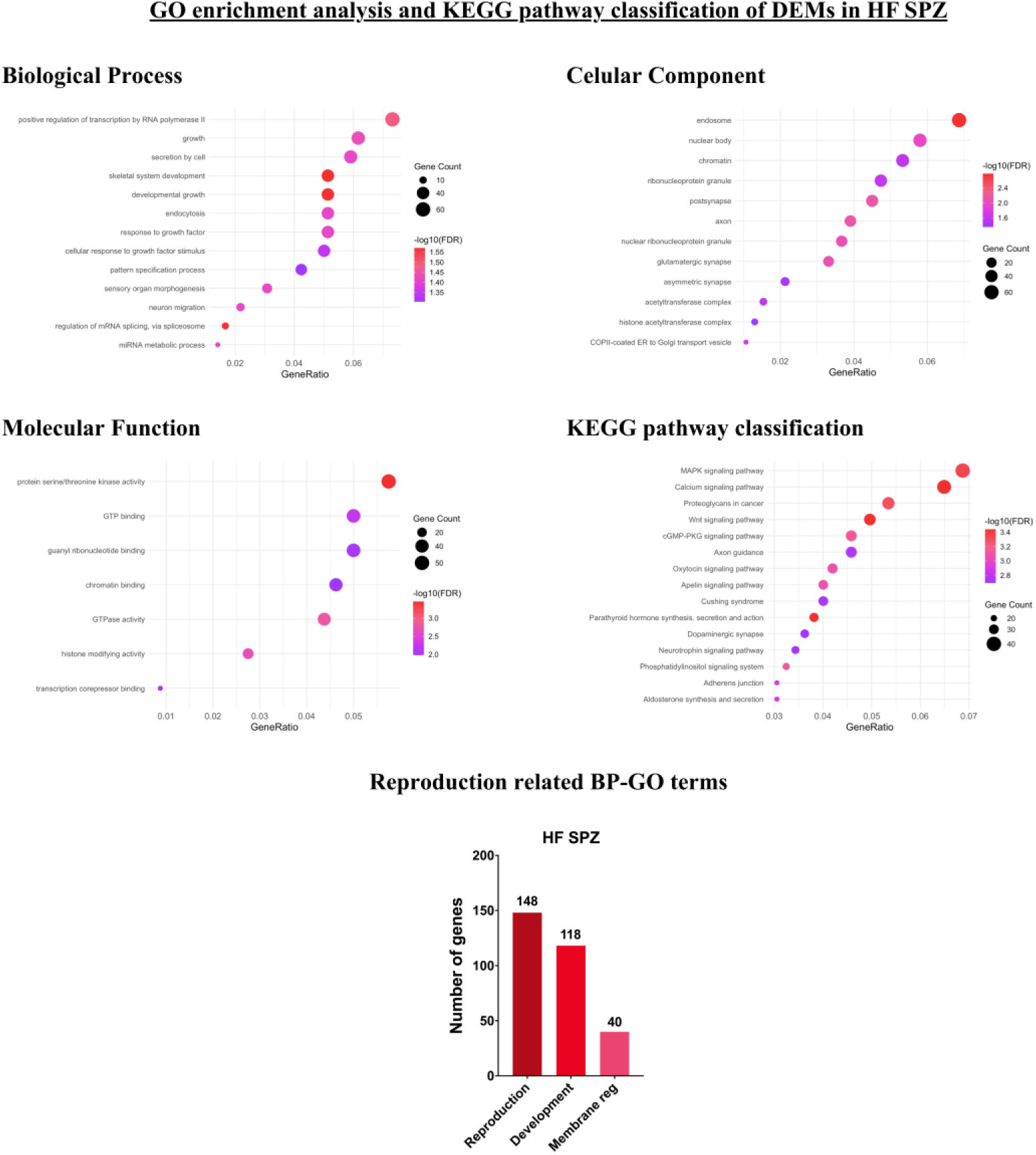
Analysis of the miRNA cargo in spermatozoa. Gene Ontology (GO) enrichment analysis and KEGG pathway classification of the target genes of the upregulated miRNAs in high fertility (HF) spermatozoa.

#### 3.3.4. Overlapping DEMs

VENN analysis showed that among the 80 DEMs identified in EVs, 10 overlapped with seminal plasma DEMs (miR-127, miR-1306-3p, miR-139-3p, miR-93-5p-like, plus six novel transcripts), and one overlapped with sperm DEMs, leaving 69 unique EV-specific DEMs. Seminal plasma and spermatozoa did not share any DEMs; seminal plasma contained 42 unique DEMs, and spermatozoa had 2 unique DEMs. No miRNAs were shared across all three sample types (**Figure 6A**). Interestingly, the four known miRNAs shared between EVs and seminal plasma displayed opposite expression patterns. miR-127 was upregulated in RF EVs but upregulated in HF seminal plasma, whereas miR-1306-3p, miR-139-3p, and miR-93-5p-like were upregulated in HF EVs but downregulated in HF seminal plasma (**Figure 6B**).

**Figure 6.**
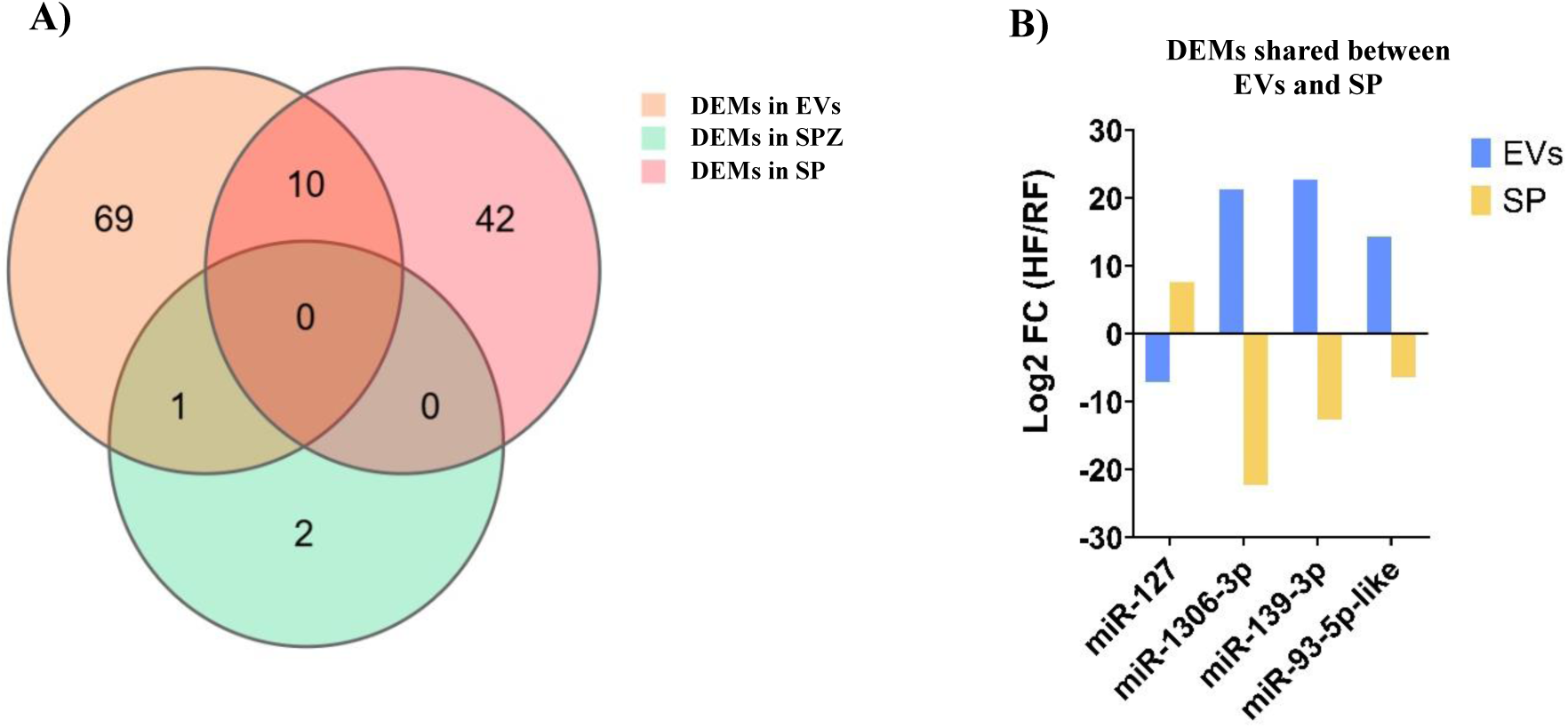
**A)** VENN representation of the shared differentially expressed miRNAs (DEMs) found between high (HF) and reduced fertility (RF) seminal extracellular vesicles (EVs), spermatozoa (SPZ) and free transcripts in seminal plasma (SP). **B)** Expression in Log2 Fold Change (FC) (HF/RF) of shared DEMs between EVs and SP.

### 3.4. Coincubation experiment

#### 3.4.1. Motility and sperm morphology

The effects of EV supplementation on sperm kinematic parameters were evaluated across treatments and incubation times. At time 0 (immediately after medium change), the percentage of TM was significantly higher in all EV-treated samples compared with the control (CTL: 12.9 ± 4.3 % vs. 1X RF_EVs: 44.9 ± 9.1 %; 2X RF_EVs: 41.4 ± 7.0 %; 2X HF_EVs: 40.8 ± 8.2 %; p < 0.05). This effect persisted after 3 hours of incubation, with the lower concentration of RF EVs exerting the strongest enhancement compared with the CTL (CTL: 16.7 ± 2.8 % vs. 1X RF_EVs: 36.4 ± 2.6 %; 2X RF_EVs: 44.3 ± 6.5 %; p < 0.05).

Similarly, PM was significantly increased in RF EV–treated samples at time 0 (CTL: 5.0 ± 2.1 % vs. 1X RF_EVs: 22.2 ± 5.3 %; 2X RF_EVs: 17.6 ± 3.2 %; p < 0.05) and after 3 hours of incubation (CTL: 9.4 ± 2.6 % vs. 1X RF_EVs: 27.9 ± 5.1 %; 2X RF_EVs: 31.6 ± 5.0 %; p < 0.05). VCL decreased during the first 6 hours of incubation in the 2X HF EV group (except at 1 hour), and at 1 hour in the 2X RF EV group (p < 0.05). VAP was significantly reduced in RF-supplemented samples at 1 hour (CTL: 83.1 ± 15.9 µm/s vs. 1X RF_EVs: 62.3 ± 2.3; 2X RF_EVs: 67.6 ± 1.3 µm/s; p < 0.05). At 6 hours, VAP was significantly reduced in the 1X HF EV group (CTL: 73.7 ± 5.4 µm/s vs. 1X HF_EVs: 49.3 ± 5.4 µm/s; p < 0.05). No significant differences were observed among treatments for VSL, WOB or LIN. STR was higher at time 0 in samples treated with 2X HF EVs (CTL: 39.9 ± 5.2 % vs. 2X HF_EVs: 57.5 ± 3.7 %; p < 0.05) and at 1 hour in the 1X HF EV group (CTL: 64.1 ± 2.0 % vs. 1X HF_EVs: 72.8 ± 1.3 %; p < 0.05) (**Figure 7**).

**Figure 7.**
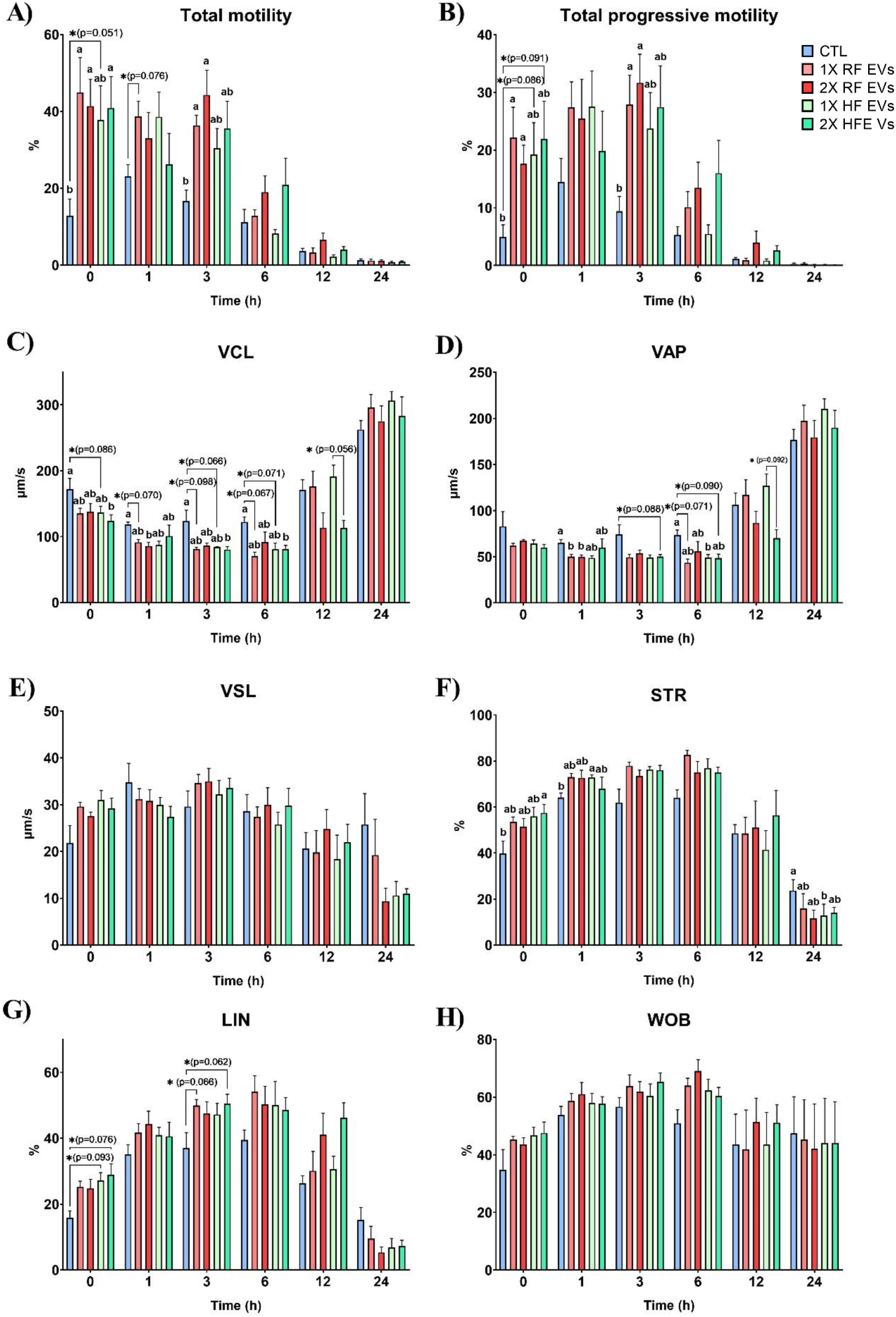
Sperm motility parameters analyzed at different time points in samples incubated 24 hours at 38 °C with high (HF) and reduced fertility (HF) seminal extracellular vesicles (EVs) at low (1X) and high concentration (2X) compared to a non-supplemented control group (CTL). Results are depicted as mean ± SEM (n = 5). The parameters analyzed were: **A)** total motility (%), **B**) total progressive motility (%), **C)** curvilinear velocity (VCL, µm/s), **D)** average path velocity (VAP, µm/s), **E)** straight line velocity (VSL, µm/s), **F)** straightness (STR, %), **G)** linearity (LIN, %) and **H)** wobble (WOB, %). Different lowercase letters indicate significant differences (p < 0.05) among different treatments in the same time point. * indicates a tendency towards significance.

Regarding sperm morphology, significant differences were observed only between RF EV concentrations. Proximal abnormalities were more frequent in 2X RF EVs compared with 1X RF EVs (2.6 ± 0.4 % vs. 3.6 ± 0.6 %; p < 0.05). Conversely, after 3 hours of incubation, distal abnormalities were lower in 2X RF EVs than in 1X RF EVs (4.4 ± 0.5 % vs. 2.5 ± 0.3 %; p < 0.05) (**Figure 8**).

**Figure 8.**
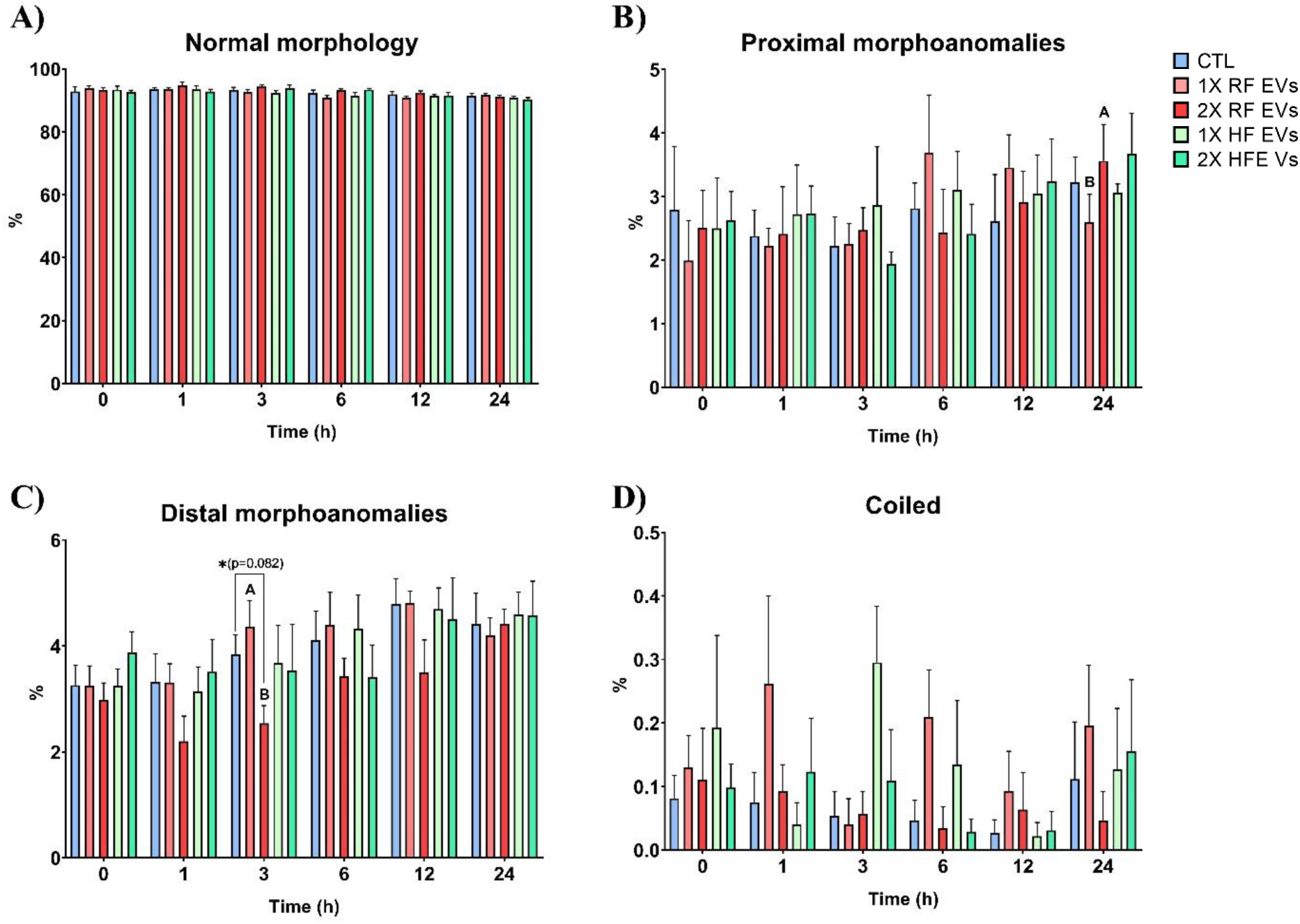
Sperm morphology analysis parameters analyzed at different time points in samples incubated 24 hours at 38 °C with high (HF) and reduced fertility (HF) seminal extracellular vesicles (EVs) at low (1X) and high concentration (2X) compared to a non-supplemented control group (CTL). Results are depicted as mean ± SEM (n = 5). The parameters analyzed were: **A)** normal morphology (%), **B)** proximal morphoanomalies (%), **C)** distal morphoanomalies (%), **D)** coiled sperm (%). Different uppercase letters indicate significant differences (p < 0.05) between low and high concentration of same fertility EVs. * indicates a tendency towards significance.

#### 3.4.2. Physiological sperm parameters analysis

##### 3.4.2.1. Sperm viability and acrosome integrity (H/PI/PNA-FITC)

After 1 hour of incubation, samples supplemented with the highest concentration of EVs exhibited a significant increase in viable sperm with intact acrosomes (H⁺/PI⁻/PNA⁻) compared with the control (CTL: 21.4 ± 1.6 % vs. 2X RF_EVs: 39.5 ± 2.8 %; 2X HF_EVs: 37.6 ± 3.0 %; p < 0.05).

Low EV concentrations showed a trend toward significance. After 3 hours, only the highest concentration of HF EVs maintained a significantly higher proportion of live, non-reacted sperm (CTL: 32.7 ± 2.8 % vs. 2X HF_EVs: 46.0 ± 2.4 %; p < 0.05) (**Figure 9A**). The proportion of live sperm with reacted acrosomes (H⁺/PI⁻/PNA⁺) increased in the 2X HF EV group after 12 h (CTL: 0.7 ± 0.1 % vs. 2X HF_EVs: 1.6 ± 0.4 %; p < 0.05) (**Figure 9B**). Dead sperm with intact acrosomes (H⁺/PI⁺/PNA⁻) showed significant differences between RF EV concentrations at 3 hours (1X RF_EVs: 26.6 ± 4.8 % vs. 2X RF_EVs: 19.4 ± 3.4 %; p < 0.05) and between CTL and 1X RF_EVs at 24 hours (CTL: 21.6 ± 1.5 % vs. 1X RF_EVs: 36.4 ± 2.7 %; p < 0.05) (**Figure 9C**). Dead sperm with reacted acrosomes (H⁺/PI⁺/PNA⁺) significantly decreased in both RF EV treatments after 1 hour (CTL: 51.9 ± 3.9 % vs. 1X RF_EVs: 35.8 ± 2.2 %; 2X RF_EVs: 34.6 ± 2.5 %; p < 0.05). After 3 hours, the lowest EV concentrations (1X RF_EVs and 1X HF_EVs) showed the lowest percentages (p < 0.05). At 6 and 12 hours, 1X RF EVs maintained significantly lower levels vs. CTL (p < 0.05). Between EV concentrations of the same group, 2X HF EVs showed higher percentages than 1X HF EVs at 6 and 24 hours (p < 0.05) (**Figure 9D**).

**Figure 9.**
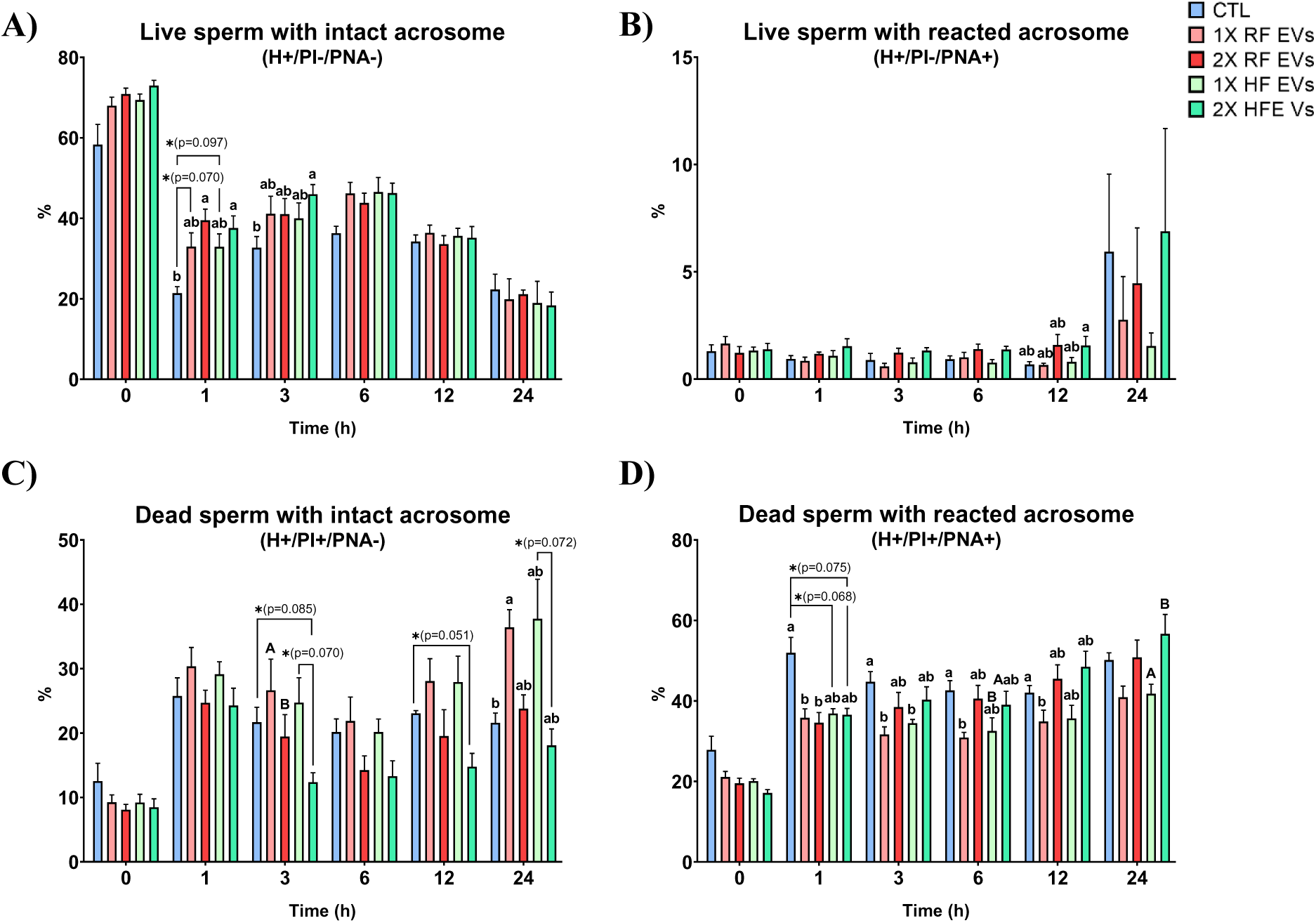
Analysis of sperm viability and acrosome integrity (H/PI/PNA-FITC) at different time points in samples incubated 24 hours at 38 °C with high (HF) and reduced fertility (HF) seminal extracellular vesicles (EVs) at low (1X) and high concentration (2X) compared to a non-supplemented control group (CTL). Results are depicted as mean ± SEM (n = 5). The parameters analyzed were: **A)** Live sperm with intact acrosome (H+/PI-/PNA-, %), **B)** Live sperm with reacted acrosome (H+/PI-/PNA+, %), **C)** Dead sperm with intact acrosome (H+/PI+/PNA-, %), **C**) Dead sperm with reacted acrosome (H+/PI+/PNA+, %). Different lowercase letters indicate differences between CTL vs. and treatments and uppercase letters indicate significant differences (p < 0.05) between low and high concentration of same fertility EVs. * indicates a tendency towards significance.

##### 3.4.2.2. Viability and mitochondrial oxidation (H/YO-PRO-1/MSox)

The high concentration of HF EVs increased the proportion of live sperm without mitochondrial oxidation (H⁺/YO-PRO-1⁻/MSox⁻) after 1 hour (CTL: 14.0 ± 2.6 % vs. 2X HF_EVs: 36.3 ± 2.5%; p < 0.05) (**Figure 10A**). The same treatment also reduced the percentage of dead sperm with oxidized mitochondria (H⁺/YO-PRO-1⁺/MSox⁺) (CTL: 64.4 ± 4.6 % vs. 2X HF_EVs: 41.7 ± 3.7%; p < 0.05) (**Figure 10D**). After 3 hours, all treatments showed reduced levels of dead sperm with mitochondrial oxidation, whereas after 6 hours, only low EV concentrations maintained this effect (p < 0.05). Among concentrations of the same fertility group, 1X RF EVs exhibited higher proportions of dead sperm with non-oxidized mitochondria at 3 and 24 hours (**Figure 10C**) and lower levels of dead sperm with oxidized mitochondria compared with HF EVs (p < 0.05) (**Figure 10D**).

**Figure 10.**
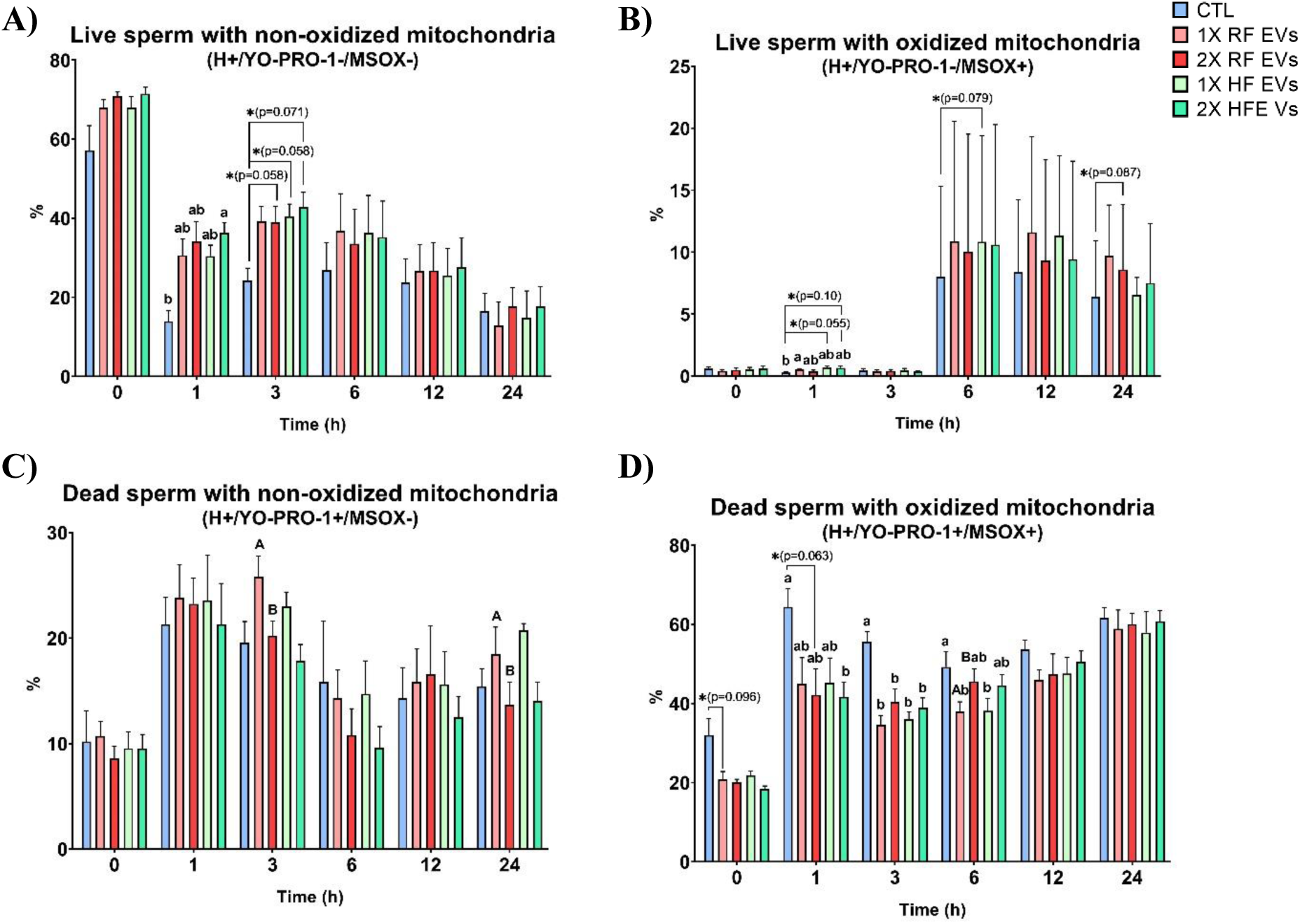
Analysis of sperm viability and mitochondrial oxidation (H/YO-PRO-1/MSox) at different time points in samples incubated 24 hours at 38 °C with high (HF) and reduced fertility (HF) seminal extracellular vesicles (EVs) at low (1X) and high concentration (2X) compared to a non-supplemented control group (CTL). Results are depicted as mean ± SEM (n = 5). The parameters analyzed were: **A)** Live sperm with non-oxidized mitochondria (H+/YO-PRO-1-/MSOX-, %), **B)** Live sperm with oxidized mitochondria (H+/YO-PRO-1-/MSOX+, %), **C)** Dead sperm with non-oxidized mitochondria (H+/YO-PRO-1+/MSOX-, %), **D)** Dead sperm with oxidized mitochondria (H+/YO-PRO-1+/MSOX+, %). Different lowercase letters indicate differences between CTL vs. and treatments and uppercase letters indicate significant differences (p < 0.05) between low and high concentration of same fertility EVs. * indicates a tendency towards significance.

##### 3.4.2.3. Viability and mitochondrial activity (H/YO-PRO-1/MTDR)

Live sperm with active mitochondria (H⁺/YO-PRO-1⁻/MTDR⁺) increased after 1 hour in HF EV–treated samples (CTL: 13.8 ± 2.6 % vs. 1X HF_EVs: 30.8 ± 2.8 %; 2X HF_EVs: 36.3 ± 2.6 %; p < 0.05). After 3 hours, 1X RF EVs showed significantly higher values (CTL: 24.0 ± 3.1 % vs. 1X RF_EVs: 39.2 ± 3.9 %; p < 0.05), while HF EVs showed a trend toward significance (p < 0.1) (**Figure 11B**). Dead sperm with inactive mitochondria (H⁺/YO-PRO-1⁺/MTDR⁻) decreased in both HF EV treatments after 1 hour (CTL: 83.5 ± 2.9 % vs. 1X HF_EVs: 63.6 ± 3.5 %; 2X HF_EVs: 59.5 ± 2.0 %; p < 0.05) and after 3 hours (CTL: 72.1 ± 3.2 % vs. 1X HF_EVs: 53.2 ± 3.1 %; 2X HF_EVs: 51.6 ± 3.4 %; p < 0.05). At 6 hours, this reduction was observed only in 1X RF EVs (CTL: 62.5 ± 1.9 % vs. 1X RF_EVs: 50.1 ± 2.3 %; p < 0.05) (**Figure 11C**).

**Figure 11.**
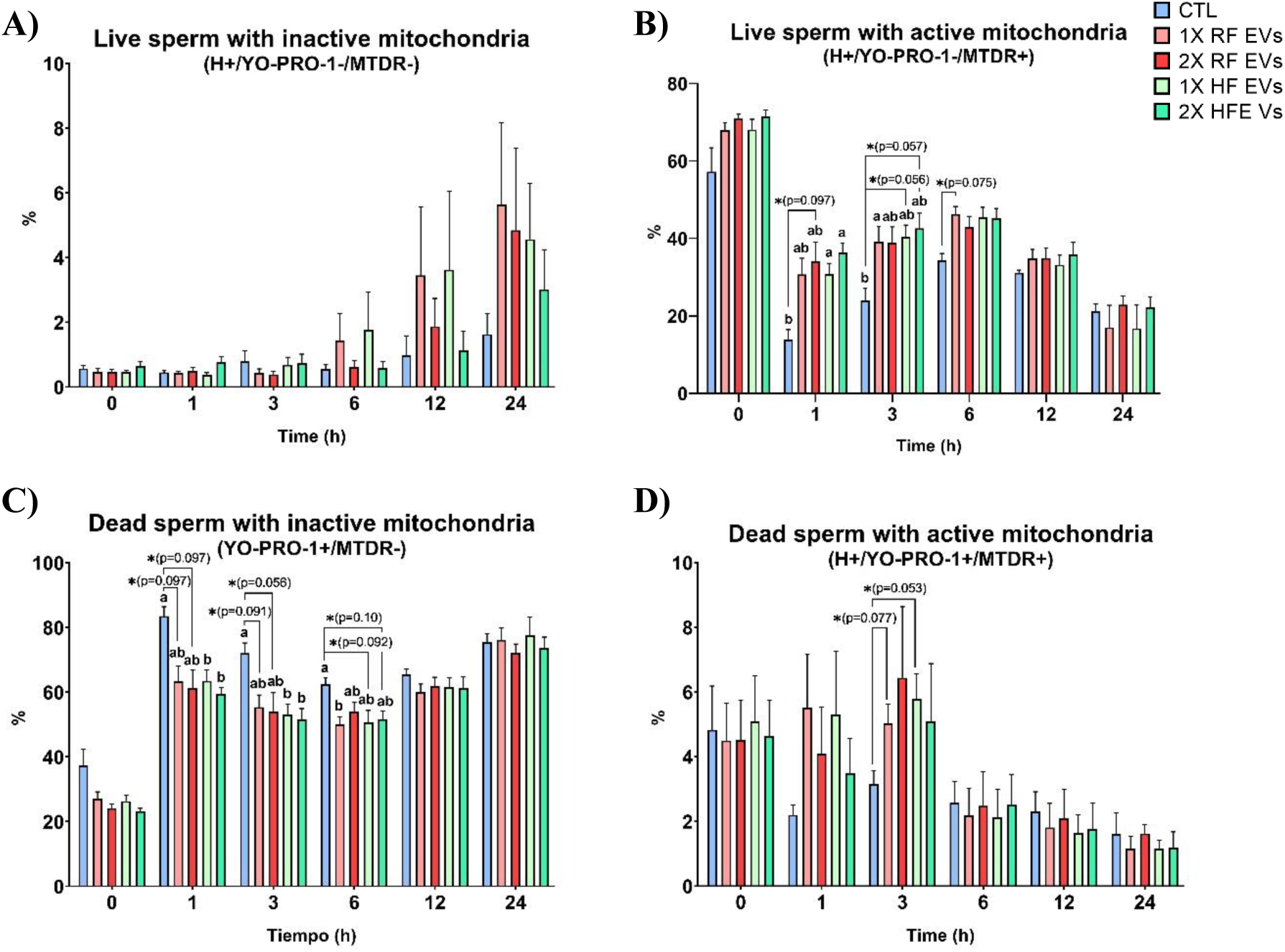
Analysis of sperm viability and mitochondrial activity (H/YO-PRO-1/MTDR) at different time points in samples incubated 24 hours at 38 °C with high (HF) and reduced fertility (HF) seminal extracellular vesicles (EVs) at low (1X) and high concentration (2X) compared to a non-supplemented control group (CTL). Results are depicted as mean ± SEM (n = 5). The parameters analyzed were: **A)** Live sperm with inactive mitochondria (H+/YO-PRO-1-/MTDR-, %), **B)** Live sperm with active mitochondria (H+/YO-PRO-1-/MTDR+, %), **C)** Dead sperm with inactive mitochondria (H+/YO-PRO-1+/MTDR-, %), **D)** Dead sperm with active mitochondria (H+/YO-PRO-1+/MTDR+, %). Different lowercase letters indicate differences between CTL vs. treatments (p < 0.05). * indicates a tendency towards significance.

##### 3.4.2.4. Viability and capacitation status (H/YO-PRO-1/M540)

After 1 hour of incubation, the percentage of live, non-capacitated sperm (H⁺/YO-PRO-1⁻/M540⁻) increased significantly in low-concentration RF and HF EV treatments (CTL: 16.6 ± 2.2 % vs. 1X RF_EVs: 23.3 ± 2.6 %; 1X HF_EVs: 25.1 ± 3.4 %; p < 0.05), while high EV concentrations showed a trend towards significance (p < 0.1) (**Figure 12A**). Live capacitated sperm (H⁺/YO-PRO-1⁻/M540⁺) tended to increase in 1X RF EVs (CTL: 2.9 ± 0.4 % vs. 1X RF_EVs: 6.7 ± 2.1%) (**Figure 12B**). Dead, non-capacitated sperm (H⁺/YO-PRO-1⁺/M540⁻) decreased in 1X RF EVs after 1 hour (CTL: 6.9 ± 1.7 % vs. 1X RF_EVs: 4.5 ± 1.8 %; p < 0.05). After 12 hours, the lowest HF EV concentration showed an increase compared with CTL and with the highest HF EV dose (CTL: 6.0 ± 0.4 % vs. 1X HF_EVs: 11.1 ± 0.9 %, p < 0.05; vs. 2X HF_EVs: 6.3 ± 1.1 %, p < 0.05) (**Figure 12C**). Dead capacitated sperm (H⁺/YO-PRO-1⁺/M540⁺) were significantly lower after 1 hour in both high-concentration EV treatments (CTL: 73.3 ± 1.8 % vs. 2X RF_EVs: 55.7 ± 2.6 %; 2X HF_EVs: 57.9 ± 3.6 %; p < 0.05). After 3 hours, only 2X HF EVs maintained this reduction (CTL: 63.2 ± 3.7 % vs. 2X HF_EVs: 49.2 ± 3.7%; p < 0.05) (**Figure 12D**).

**Figure 12.**
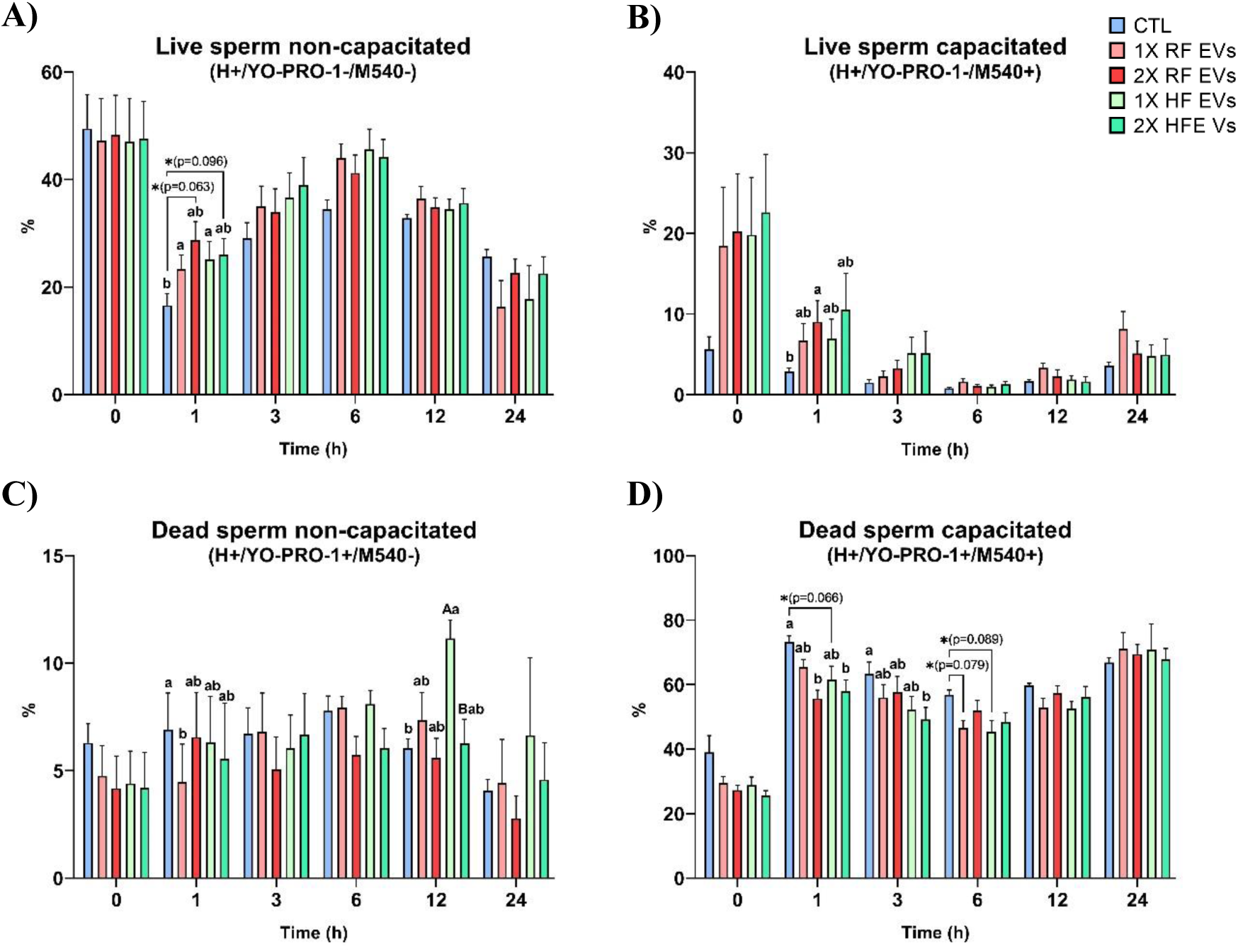
Analysis of sperm viability and capacitation status (H/YO-PRO-1/M540) at different time points in samples incubated 24 hours at 38 °C with high (HF) and reduced fertility (HF) seminal extracellular vesicles (EVs) at low (1X) and high concentration (2X) compared to a non-supplemented control group (CTL). Results are depicted as mean ± SEM (n = 5). The parameters analyzed were: **A)** Live sperm non-capacitated (H+/YO-PRO-1-/M540-, %), **B)** Live sperm capacitated (H+/YO-PRO-1-/M540+, %), **C)** Dead sperm non-capacitated (H+/YO-PRO-1+/M540-, %), **D)** Dead sperm capacitated (H+/YO-PRO-1+/M540+, %). Different lowercase letters indicate differences between CTL vs. treatments and uppercase letters indicate significant differences (p < 0.05) between low and high concentration of same fertility EVs. * indicates a tendency towards significance.

##### 3.4.2.5. Viability and sperm oxidation (H/YO-PRO-1/DHE)

At 1 hour, both HF EV concentrations significantly increased the proportion of live, non-oxidized sperm (H⁺/YO-PRO-1⁻/DHE⁻) (CTL: 16.5 ± 2.1 % vs. 1X HF_EVs: 30.3 ± 2.5 %; 2X HF_EVs: 35.5 ± 2.7 %; p < 0.05). At 3 hours, this effect was maintained in the high RF EV concentration and both HF EV treatments (CTL: 25.8 ± 2.7 % vs. 2X RF_EVs: 36.4 ± 3.8 %; 1X HF_EVs: 41.0 ± 2.9 %; 2X HF_EVs: 45.6 ± 2.5 %; p < 0.05) (**Figure 13A**). Consistent with these results, dead oxidized sperm (H⁺/YO-PRO-1⁺/DHE⁺) decreased significantly in all EV-treated samples at 1 hour (p < 0.05). After 6 hours, only the lowest HF EV concentration maintained this reduction (CTL: 57.0 ± 2.6 % vs. 1X HF_EVs: 44.1 ± 3.0 %; p < 0.05) (**Figure 13D**). No significant differences were observed at later time points.

**Figure 13.**
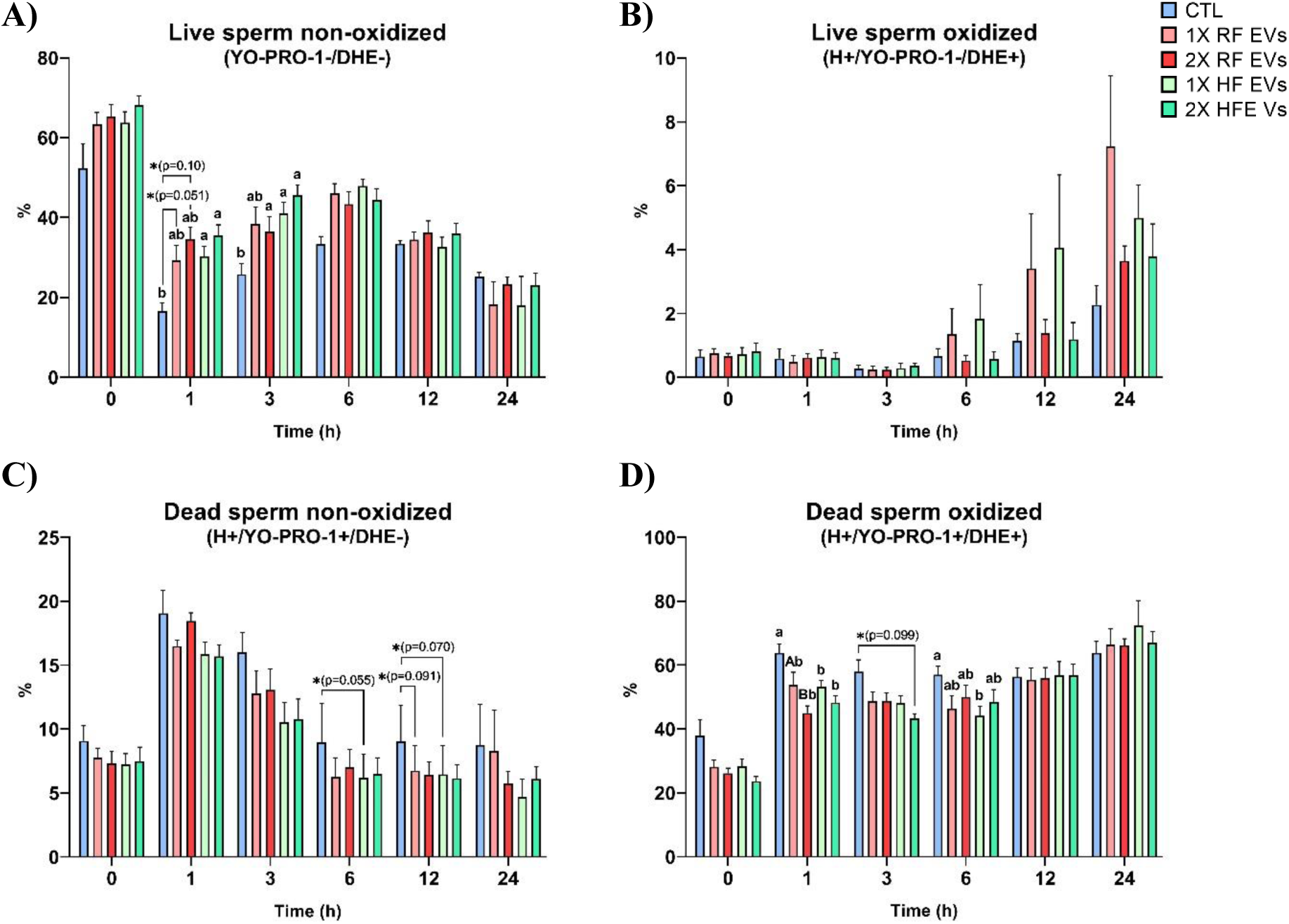
Analysis of sperm viability and oxidation (H/YO-PRO-1/DHE) at different time points in samples incubated 24 hours at 38 °C with high (HF) and reduced fertility (HF) seminal extracellular vesicles (EVs) at low (1X) and high concentration (2X) compared to a non-supplemented control group (CTL). Results are depicted as mean ± SEM (n = 5). The parameters analyzed were: **A)** Live sperm non-oxidized (H+/YO-PRO-1-/DHE-, %), **B)** Live sperm oxidized (H+/YO-PRO-1-/DHE+, %), **C)** Dead sperm non-oxidized (H+/YO-PRO-1+/DHE-, %), **D)** Dead sperm oxidized (H+/YO-PRO-1+/DHE+, %). Different lowercase letters indicate differences between CTL vs. and treatments and uppercase letters indicate significant differences (p < 0.05) between low and high concentration of same fertility EVs. * indicates a tendency towards significance.

## Discussion

Worldwide porcine production has increased in the last decade (USDA Foreign Agricultural Service), in part due to global population growth. This trend has driven the evolution of reproductive procedures, aiming to increase productivity and reduce costs. In this context, the early detection of subfertile males before entering the production chain is crucial. The use of diluted seminal doses for AI has expanded considerably (M.L.W.J. Broekhuijse et al., 2012). They are usually optimized for sperm concentration, while other seminal components, such as EV cargo, receive less attention despite their potential roles as key fertility regulators. In this work, we examined the morphology and integrity of seminal EVs in porcine semen doses, their protein and miRNA cargo and their functional effects after high dilution.

Seminal EVs are produced in the sexual glands and carry proteins such as integrins, tetranspanins and lectins on their membranes, enabling interactions with sperm cells and modulation of their functional state (Jankovičová et al., 2020). Although the precise mechanisms governing EV uptake by spermatozoa are not fully understood, they have been proposed to involve lipid raft–mediated endocytosis (Murdica et al., 2020) or receptor-mediated processes (Zhou et al., 2019). Different biomolecules, including proteins and miRNAs carried by seminal EVs, have been associated with male fertility across species, suggesting that EVs may convey specialized cargo that influences sperm maturation, fertilization, and early embryo development (Aalberts et al., 2014; Ding et al., 2021; Dlamini et al., 2023).

To better understand the role of seminal EVs in male fertility, we isolated EVs from commercial semen doses. TEM confirmed a normal cup-shaped morphology, and NTA indicated high particle concentration and a typical size distribution after EV enrichment, supporting that seminal EVs remain structurally intact despite ejaculate dilution. During storage, which can last several days, these EVs remain in intimate contact with sperm cells and therefore may influence their function. Our proteomic characterization further supports the potential use of seminal EVs as biomarkers of boar fertility status.

Proteomic analysis (as well as Western Blot and flow cytometry analysis) identified classical EV markers, including tetraspanins and luminal proteins (CD44, CD81, CD9, ALIX, TSG101, FLOT1, FLOT2, HSP70, HSP90), and importantly, no classical negative markers of cellular or plasma contamination (Calnexin (CANX), GM130, GRP94 and albumin) were detected. Differential expression analysis revealed 108 DEPs between HF and RF EVs, of which 97 were upregulated in HF EVs and 11 in RF EVs, indicating clear fertility-associated differences in EV protein cargo that may influence reproductive outcomes. GO enrichment analysis for RF DEPs had limited statistical power due to the low number of proteins; however, several RF-enriched proteins have known roles in fertility. Toll-like receptor 2 (TLR2) participates in sperm-triggered uterine inflammatory responses at insemination (Mansouri et al., 2023) and can accelerate the acrosome reaction (Fujita et al., 2011). Carbohydrate-binding protein AQN-1, also known as acrosin inhibitor acceptor molecule, is a spermadhesin implicated in sperm–zona pellucida recognition and binding (Sanz et al., 1992), but it also has decapacitating effects (Caballero et al., 2012), which could lead to a reduction of the fertility of these animals. The dickkopf-related protein 1 (DKK1) has been described as a regulator of bovine embryo development by inhibiting WNT signaling (Amaral et al., 2022; Denicol et al., 2015). Several RF DEPs were related to vesicular trafficking, including SMIM22, GOLIM4, GOLGB1, and ASPH (Luo et al., 2021; Papadopoulou et al., 2019; Ramalho-Santos et al., 2001; Zhao et al., 2024). Actin (ACTB), also upregulated in RF EVs, modulates sperm capacitation (Breitbart et al., 2005) and is abundant in oviductal fluid in post-ovulatory stages (Lamy et al., 2018). Excessive exposure of spermatozoa to ACTB before reaching the appropriate site and time for fertilization could promote premature capacitation and reduce their fertilizing capacity in the oviduct. RNASE4, the most upregulated RF protein, has been detected in seminal plasma and proposed as part of host defense against pathogens (Gupta et al., 2016), but it also affects the integrity of miRNAs in RF EVs. Overall, most of these proteins seem to be related to the capacitation process and could affect negatively to the oocyte fertilization, potentially contributing to the reduced fertility of these animals

In contrast, the larger number of DEPs upregulated in HF EVs allowed robust GO enrichment, revealing biological processes related not only to metabolism (catabolic processes, proteolysis, carbohydrate metabolism) but also to reproduction, including sperm–egg recognition, zona pellucida binding, and vesicle-mediated transport. Several HF enriched proteins had already been implicated in key sperm functions. Acrosin (ACR), an acrosomal protease, is essential for zona pellucida penetration and oocyte fertilization (Hirose et al., 2020). MMP7, a matrix metalloproteinase, has been associated with sperm differentiation, capacitation, viability, fertilization, inflammatory modulation and sperm DNA fragmentation (Asgari et al., 2021). ATP6V1A, the catalytic subunit of the V-ATPase complex, regulates acrosomal acidification, a prerequisite for enzyme activation (Sun-Wada et al., 2002; Visconti et al., 2011). FURIN has been proposed as a fertility biomarker in boars (Pérez-Patiño et al., 2018), likely through its role in epididymal fluid and sperm membrane regulation. Similarly, IL4I1, a ROS-producing enzyme in the acrosomal region (Houston et al., 2015), has been suggested as a fertility biomarker in bovine sperm (D’Amours et al., 2019). Collectively, these proteins seem to participate in proper acrosome development and the fertilization process, which could explain the higher fertility rate of these animals.

Several DEPs identified here are associated with cytoskeletal organization, especially actin filament dynamics, which are central to capacitation. Capacitation involves extensive cytoskeletal remodeling, including actin polymerization between the outer acrosomal and plasma membranes to stabilize the acrosome and prevent premature exocytosis. Subsequent F-actin depolymerization is required for the acrosome reaction (Breitbart and Finkelstein, 2018). In this context, multiple DEPs detected in EVs are known modulators of actin turnover. Notably, CNN3 and WDR1, two regulators of actin dynamics, were differentially expressed. CNN3 promotes F-actin depolymerization through phosphorylation-dependent mechanisms, supporting membrane flexibility and progression of capacitation (Shen et al., 2021). WDR1, an actin-interacting protein, facilitates filament disassembly and has been linked to structural transitions associated with the acrosome reaction (Staicu et al., 2021). Their differential abundance suggests that actin remodeling is actively modulated by EV cargo and may be critical during fertilization. The identification of CCT3 and CCT7, subunits of the TRiC chaperonin complex, further highlights cytoskeletal regulation. The TRiC complex has been detected on the surface of capacitated sperm and proposed to participate in zona pellucida binding (Dun et al., 2011), suggesting that its presence in EVs could influence sperm–oocyte interactions and fertilization efficiency.

Other DEPs with roles in cytoskeletal and structural regulation include SIRT2, ARL3 and DAG1. SIRT2 is involved in spermatogenesis (Feng et al., 2024), has antioxidant functions (Tatone et al., 2018), and its inhibition is associated with reduced blastocyst cell numbers (Kwak et al., 2012), linking it to both sperm function and early embryo development. ARL3 is critical for manchette formation during spermiogenesis and participates in ciliary trafficking and intraflagellar transport (Qi et al., 2013; Umer et al., 2022), suggesting that its presence in EVs may reflect transfer from epididymosomes during early sperm maturation. DAG1, reported in bull seminal plasma but absent from epididymal sperm, appears to be acquired at ejaculation and may facilitate cell–cell adhesion events, relevant to sperm–oocyte interactions (Zoca et al., 2023).

Interestingly, GO analysis indicated that some HF DEPs (e.g., MMP7, GBA, ST3GAL1, GGT1, PAFAH1B1, S100A14) may have immune regulatory functions. Seminal plasma triggers a controlled immune response in the female tract, inducing an inflammatory reaction that promotes embryo implantation while establishing immune tolerance (Schjenken et al., 2021).

Beyond proteins, seminal plasma also carries miRNAs both in solution and within EVs. The precise mechanisms by which these transcripts exert their effects remain unclear, but miRNAs have been implicated in post-mating immune modulation in the endometrium (Barranco et al., 2020). Our small RNA-seq data showed high mapping quality, particularly for sperm-derived miRNAs, supporting the robustness of the downstream analyses and aligning well with the proteomic findings. Notably, a significantly higher number of DEMs were detected in HF EVs compared with RF EVs, which may reflect a richer or more functionally diverse cargo in HF EVs. GO analysis revealed fertility-related terms enriched in HF EVs, including “embryo development ending in birth or egg hatching”. Seminal EVs have previously been linked to embryo development (Rodriguez-Martinez and Roca, 2022), and some HF EV miRNAs may target key genes such as BMP4, NOTCH1, and WNT5A (Hayward et al., 2008; Li and Parast, 2014), potentially supporting improved embryonic development. Reproduction-related enrichment was more pronounced in HF EVs, particularly for pathways associated with membrane regulation, suggesting that EV-borne miRNAs could contribute to critical reproductive processes. Several miRNAs identified here have already been linked to reproductive function. For instance, miR-10a-3p modulates immune activity by suppressing inflammatory responses (Kiran et al., 2023). Additional DEMs identified in this study also have documented links to reproductive tissues or functions. miR-1343 has been detected in follicular fluid (Pasquariello et al., 2020), whereas miR-149 is present in Sertoli cell–derived EVs (Tan et al., 2022) and cumulus cells (Wu et al., 2024), pointing to a potential role in gamete maturation. miR-15b is upregulated in HF boar sperm (Alvarez-Rodriguez et al., 2020), and miR-193b-5p is enriched in EVs from fertile bucks (Sakr et al., 2023), both supporting their association with male fertility. miR-215-5p, secreted in endometrial EVs (Cai et al., 2025), may mediate communication between the uterine environment and the embryo. Mir-27a and miR-296-5p, upregulated in HF bulls, have been associated with the acrosome reaction (Kasimanickam and Kasimanickam, 2024). Together, these observations reinforce the emerging view that EV-associated miRNAs participate in the molecular communication underpinning sperm quality, fertilization, implantation and early embryo development.

Given the high genomic similarity between pigs and humans, we performed BLAST analyses of novel DEMs in EVs and identified new *Sus scrofa* miRNAs not previously described. Some of these novel miRNAs have established roles in human reproduction. For example, miR-188-5p participates in embryonic development (Pu et al., 2022), while miR-203a-3p has been linked to embryo implantation (Yu et al., 2025) and is present in uterine EVs (Deng et al., 2022). miR-33a-5p, upregulated in seminal EVs from fertile bucks (Sakr et al., 2023), regulates immune responses and cholesterol efflux (Oladosu et al., 2024), a key process during sperm capacitation (Dudkiewicz et al., 2024). Altogether, these data suggest that the novel *Sus scrofa* miRNAs identified here may play important roles in reproduction and warrant further functional investigation in this species.

Free (non–EV-encapsulated) miRNAs in seminal plasma were also differentially expressed. We identified 33 miRNAs upregulated in HF and 19 in RF seminal plasma, mirroring the pattern observed in EVs. GO analysis revealed both shared terms (e.g., cell morphogenesis, gland development) and distinct patterns between fertility groups. Whereas HF showed enrichment for negative regulation of RNA biosynthetic processes, RF showed positive regulation of transcription by RNA polymerase II. The functional consequences of this contrast are complex, as seminal plasma miRNAs can act on both the female and male sides, but proper transcriptional regulation is known to underpin normal cell and organ function and is therefore linked to fertility (Saftić Martinović et al., 2024). In addition, HF seminal plasma was enriched for terms such as regulation of cellular component biogenesis, positive regulation of cell differentiation and embryonic/tissue morphogenesis, whereas RF was associated with growth, Wnt signaling and cellular responses to oxygen. The molecular function terms were broadly similar between HF and RF SP, with HF-specific enrichment in metal ion transmembrane transporter activity, which could affect Ca²⁺ fluxes in sperm via channels such as TRPC4 (de Mercado et al., 2026), and in actin binding, consistent with its known role in capacitation (Breitbart and Finkelstein, 2018).

Several HF-upregulated DEMs in seminal plasma have been linked to reproductive functions. miR-100 appears to promote spermatogonial stem cell proliferation (Huang et al., 2017) and exerts immunoregulatory effects (Long et al., 2021), while miR-326 secretion has been detected in granulosa cells (Chaurasiya et al., 2020). In contrast, although miR-29b was upregulated in HF SP in our study, it has been associated with low fertility in bulls (Menezes et al., 2020), highlighting potential species- or context-specific differences. Some RF-upregulated DEMs are associated with impaired male fertility, such as miR-141 (Senousy et al., 2025) and miR-423-5p, which has been linked to oxidative stress and reduced sperm motility (Zhang et al., 2021).

Following the BLAST analysis of novel miRNAs in EVs, we performed a similar approach for seminal plasma DEMs and found two novel transcripts with high homology to human miR-93-5p and miR-200a-3p, both implicated in immune regulation and embryo development (Saha et al., 2015).

Interestingly, the Venn diagram showed that three known miRNAs and one novel transcript were shared between EVs and SP but displayed opposite expression patterns. miR-127 was upregulated in RF EVs but upregulated in HF seminal plasma, whereas miR-1306-3p, miR-139-3p and miR-93-5p-like were upregulated in HF EVs but upregulated in the seminal plasma of RF males. This inverse relationship may be functionally relevant. Due to the EV lipid bilayer, miRNA cargo is protected from RNases present in seminal plasma (D’Alessio et al., 1972), enhancing their stability and facilitating targeted delivery to recipient cells.

Comparing GO analysis between HF seminal plasma and HF EVs revealed common terms such as neuron development, positive regulation of cell differentiation, vasculature development, tissue morphogenesis and gland development. However, HF seminal plasma showed additional terms such as embryonic morphogenesis and plasma membrane regulation and, in contrast to HF EVs, negative regulation of RNA biosynthetic processes. RF seminal plasma exhibited terms (e.g., growth, cellular response to oxygen-containing compounds, cell surface receptor tyrosine kinase signaling, canonical Wnt signaling) that were absent in RF EVs, again supporting a complementary but distinct regulatory role of free vs EV-associated miRNAs.

Differential miRNA expression between sperm from HF and low fertility males has been described previously (Alvarez-Rodriguez et al., 2020). Although transcription and translation are largely silent in mature sperm, sperm-delivered miRNAs can be transferred to the zygote and modulate maternal mRNAs (Liu et al., 2012). In our study, three novel miRNAs (one of them shared with EV DEMs) were upregulated in HF spermatozoa. GO analysis indicated roles in growth, endocytosis, and signaling pathways such as MAPK, calcium and Wnt, all of which are highly relevant to fertilization and early embryo development.

Based on the combined transcriptomic and proteomic evidence for differential EV cargo, we next investigated whether EVs from semen doses are still functionally active and capable of modifying sperm quality. To this end, we coincubated sperm from RF males with different concentrations of HF and RF EVs and assessed changes in sperm quality parameters over time. Overall, our results indicate a beneficial effect of EVs on many sperm quality traits, particularly during the first hours of incubation, during which HF EVs improved sperm viability and reduced oxidative stress.

Sperm motility and kinematic parameters are positively correlated with male fertility (Bae et al., 2024). In our coincubation experiments, RF EVs increased total and progressive motility, whereas HF EVs did not markedly improve motility parameters. This may reflect a degree of “compatibility” between EVs and sperm from the same fertility group, as both originate from RF males and may share cargo that better supports RF sperm function (Budden et al., 2021; Haghighitalab et al., 2023). In contrast, velocity parameters such as VCL and VAP tended to decrease in all treatments. Increases in these parameters are associated with hyperactivation during capacitation (Ramió et al., 2008); thus, their reduction suggests that EVs may help modulate or restrain premature capacitation. Indeed, M540 staining revealed decreased capacitation in live and dead sperm after 1 hour with low EV doses from both fertility groups. The upregulation in HF EVs of proteins such as MMP7 and CNN3, which regulate capacitation-related processes, supports this interpretation. Conversely, the high concentration of RF EVs increased the proportion of capacitated sperm after 1 hour, possibly reflecting the presence of pro-capacitation miRNAs and proteins such as ACTB, which promotes capacitation (Breitbart et al., 2005).

Acrosome integrity, mitochondrial function and oxidative status were also affected by coincubation. The proportion of live sperm with intact acrosomes increased after 1 hour when using the highest EV concentration from both groups. Although some studies reported no effect of EVs on acrosome status (Barranco et al., 2024; Piehl et al., 2013), likely due to methodological differences, our findings align with previous work demonstrating inhibitory effects on acrosome reaction in human (Pons-Rejraji et al., 2011) and bull sperm (Pal et al., 2025). After 3 hours, this protective effect persisted only with the highest HF EV dose, suggesting a stronger and more sustained protective capacity of HF EVs. This is consistent with reports showing that EV proteins and miRNAs from low-fertility males can negatively affect sperm viability (Chen et al., 2024). RF EVs also reduced the proportion of dead sperm with reacted acrosomes after 1 hour, and this effect was most evident at lower EV concentrations. However, after 3 hours the protective effect appeared to diminish, possibly reflecting a reduced EV–sperm interaction over time.

GO analysis also highlighted metabolic and oxygen-related pathways, which is consistent with the observed effects on mitochondrial activity. HF EVs increased the percentage of sperm with active mitochondria and decreased the proportion with inactive mitochondria after 1 and 3 hours, in agreement with previous reports describing ATP-generating enzymes in boar seminal EVs that enhance mitochondrial membrane potential and activity (Guo et al., 2019). This enhanced mitochondrial function is likely linked to reduced oxidative damage. Both HF EV doses lowered sperm oxidative stress, and the higher dose in particular reduced mitochondrial oxidation. The presence of antioxidant proteins in seminal EVs (Guo et al., 2019) and miRNAs that activate anti-oxidative pathways likely contributes to decreased ROS production in sperm and mitochondria. Excessive ROS are deleterious metabolic by-products associated with sperm damage and infertility (O’Flaherty and Matsushita-Fournier, 2017), and reduced ROS after EV supplementation has also been reported in recent studies (Pal et al., 2025; Parra et al., 2025).

Overall, this study demonstrates that seminal plasma EVs isolated from commercial semen doses preserve their structural integrity, display fertility-associated differences in protein and miRNA cargo between HF and RF males, and show inverse relationships with free miRNAs in seminal plasma. Moreover, we demonstrate that these EVs remain functionally active, improving sperm quality by enhancing viability and mitochondrial activity, reducing sperm and mitochondrial oxidation, and modulating the timing of capacitation. These findings support the potential of seminal EVs as biomarkers and as functional modulators that could be exploited to improve reproductive technologies. Although implementation in commercial AI schemes may be technically challenging, the addition or modulation of seminal EVs in semen doses should be considered as a promising strategy to optimize boar fertility.

## Conclusions

This study demonstrates that seminal EVs from porcine AI doses retain structural integrity after dilution, carry fertility-associated molecular cargo, and exert measurable functional effects on spermatozoa. Proteomic and miRNA profiling revealed clear differences between HF and RF males. HF EVs were enriched in proteins and miRNAs related to vesicle transport, cytoskeletal regulation, acrosome function, metabolic activity and embryo development, whereas RF EVs contained molecules potentially linked to premature capacitation, oxidative stress or impaired immune modulation. Several novel *Sus scrofa* miRNAs with homology to human reproductive miRNAs were identified, highlighting conserved regulatory pathways. Functionally, HF EVs improved sperm viability, preserved acrosome integrity, enhanced mitochondrial activity and reduced oxidative stress, while RF EVs mainly increased motility and influenced capacitation status. These findings confirm that seminal EVs actively modulate sperm physiology and that their cargo could be used to increase RF spermatozoa quality. Overall, seminal EVs emerge as promising biomarkers of boar fertility and potential tools to optimize sperm quality and reproductive performance in swine production systems.

## Author contribution

**Adrián Martín-San Juan**: writing and editing– original draft, investigation, conceptualization, methodology, software, bioinformatics and data analysis. **Claudia Cerrato Martín-Hinojal**: review & editing, investigation, software, methodology, data analysis. **Maria José Martinez-Alborcia**: review & editing, investigation, methodology. **Helena Nieto-Cristóbal**: review & editing, conceptualization, software, methodology. **Eduardo de Mercado**: Writing and editing – original draft, software, project administration, methodology, funding acquisition, investigation, conceptualization. **Manuel Álvarez-Rodríguez**: Writing and editing – original draft, supervision, software, project administration, methodology, funding acquisition, conceptualization.

## Aknowledgments

To Mariano José Rangil Escribano, Sandra Blanco López, and AIM Ibérica (TOPIGS Norsvin) for the commercial AI doses used to conduct these experiments. To Paco Blasco from SPERMTECH, for our collaborative agreement to establish AIStation analysis software in our Animal Reproduction Lab. To Maria Teresa Vallejo Cremades of the Nanoparticle Tracking Analysis service located at IdiPAZ (Madrid, Spain), and to Beatriz Martín Jouve of the Transmission Electron Microscopy service located at Centro Nacional de Biotecnología (CNB, Madrid, Spain).

## Data Availability Statement

The data that support the findings of this study are available from the corresponding author upon reasonable request.

## Controls

**Supplementary Figure 1.**
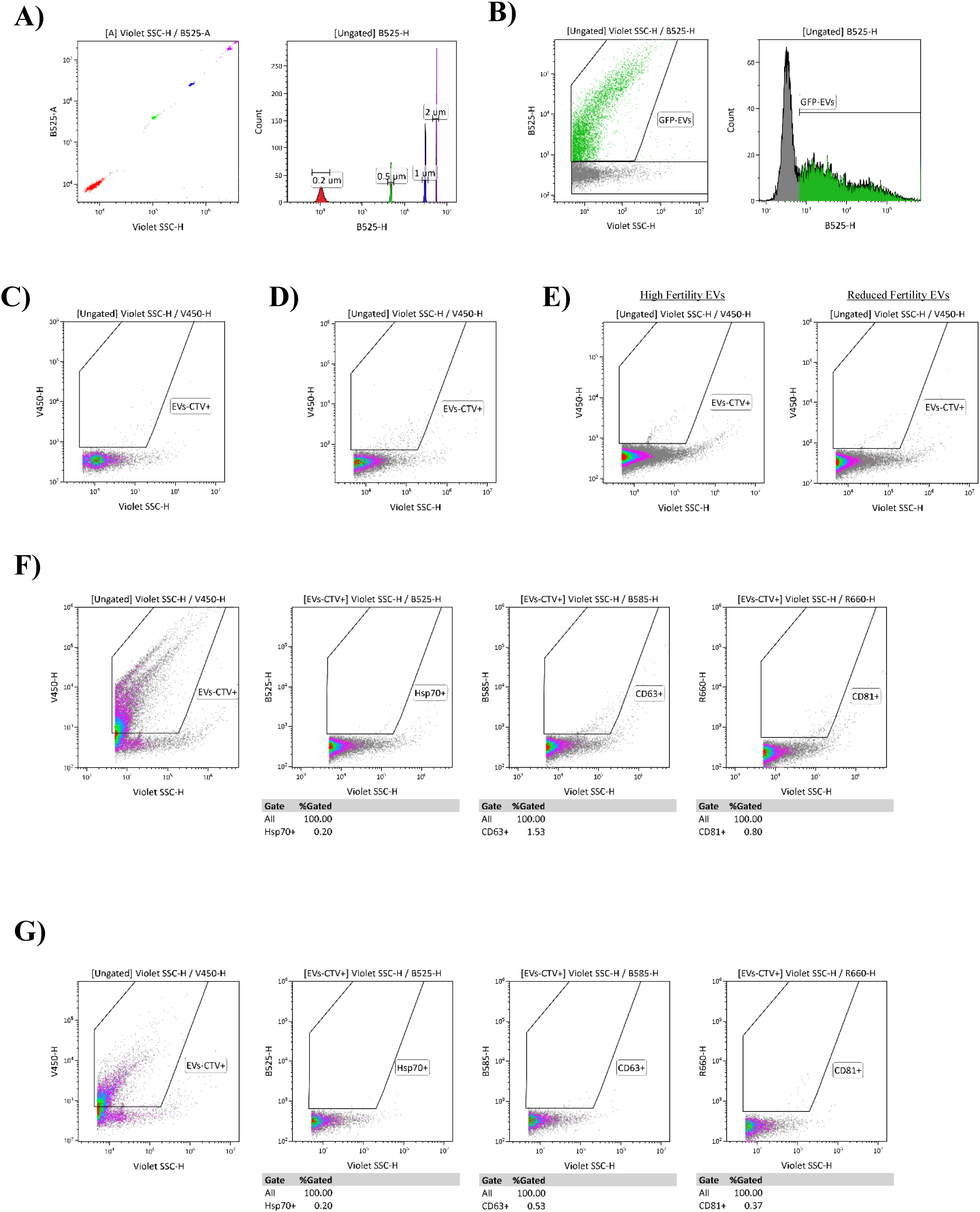
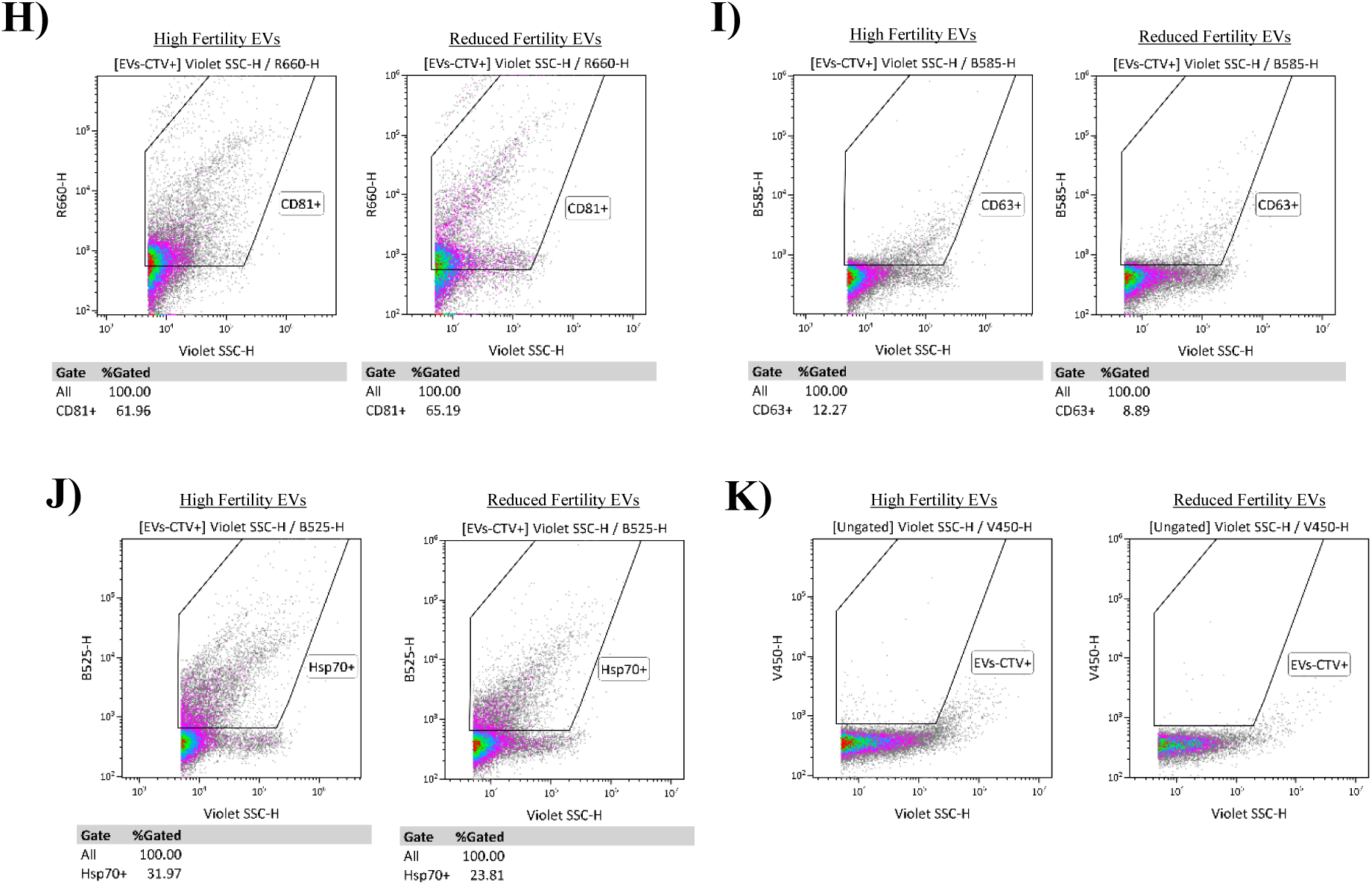
Cytometry plots and histograms of seminal extracellular vesicles (EVs) characterization. **A)** Flow cytometry beads of know size (200 nm – 2 µm). **B)** GFP EVs were used to identify EVs population. **C)** PBS 1X 0.22 μM filtered. **D)** Negative control of CellTrace Violet (CTV) staining diluted in PBS. **E)** High fertility (HF) and reduced fertility (RF) unstained EVs. **F)** HF CTV-stained EVs. **G)** RF CTV-stained EVs. **H)** CTV-stained EVs + human anti CD81-APC antibody (1:50). **I)** CTV-stained EVs + human anti CD63-PE antibody (1:50). **J)** CTV-stained EVs + human anti Hsp70-FITC antibody (1:50). **K)** Lysated EVs with Triton 1X + 0.1% SDS. Photodetectors used: Violet Side Scatter – Height (Violet SSC-H), Blue 525-Area (B525-A), Blue 525-Height (B525-H), Violet 450-Height (V450-H), Blue 585-Height (B585-H) and Red 660-Height (R660-H).

**Supplementary Figure 2.**
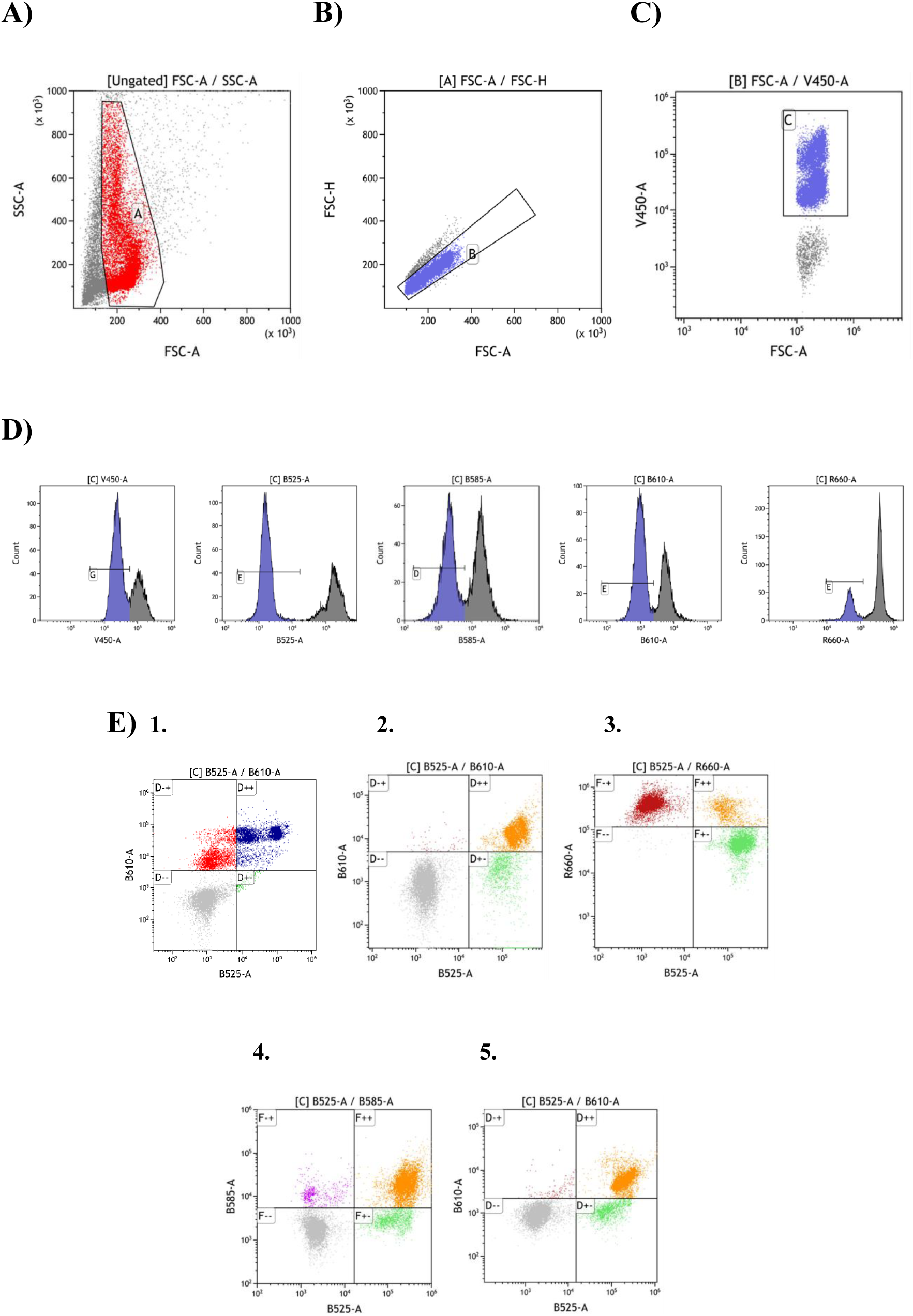
Cytometry plots and histograms for physiological sperm parameters analysis. **A)** Forward (FSC) and side scatter (SSC) plot was used to identify sperm population and exclude debris. **B)** FSC-height and FSC-area plot was used to select singlet events. **C)** FSC-area and V450-area plot was used to identify nucleated sperm and exclude debris from the analysis. **D)** Representative histograms of the photodetectors used in the analysis. The gated region corresponds to the negative population of unstained cells, whereas events outside the gate represent positively fluorescent cells (Hoechst 33342: V450-A; fluorescein isothiocyanate-conjugated peanut (*Arachis hypogaea*) agglutinin (PNA-FITC) and YO-PRO-1: B525-A; Propidium iodide (PI), MitoSOX Red (MSox) and dihydroethidium (DHE): 610-A; Mitotracker Deep Red (MTDR): 660-A; Merocyanine 540 (M540): 585-A). **E)** Histograms of the combined stainings divided in the four population for each staining method (see *2.9.2. Physiological sperm parameters analysis*). **1)** B525-A/B610-A for viability and acrosome integrity (PI/PNA-FITC) analysis. **2)** B525-A/B610-A for viability and mitochondrial oxidation (YO-PRO-1/MSox). **3)** B525-A/B660-A for viability and mitochondrial activity (YO-PRO-1/MTDR). **4)** B525-A/B585-A for viability and membrane fluidity (YO-PRO-1/M540). **5)** B525-A/610-A for viability and sperm oxidation (YO-PRO-1/DHE).

**Supplementary Figure 2.**
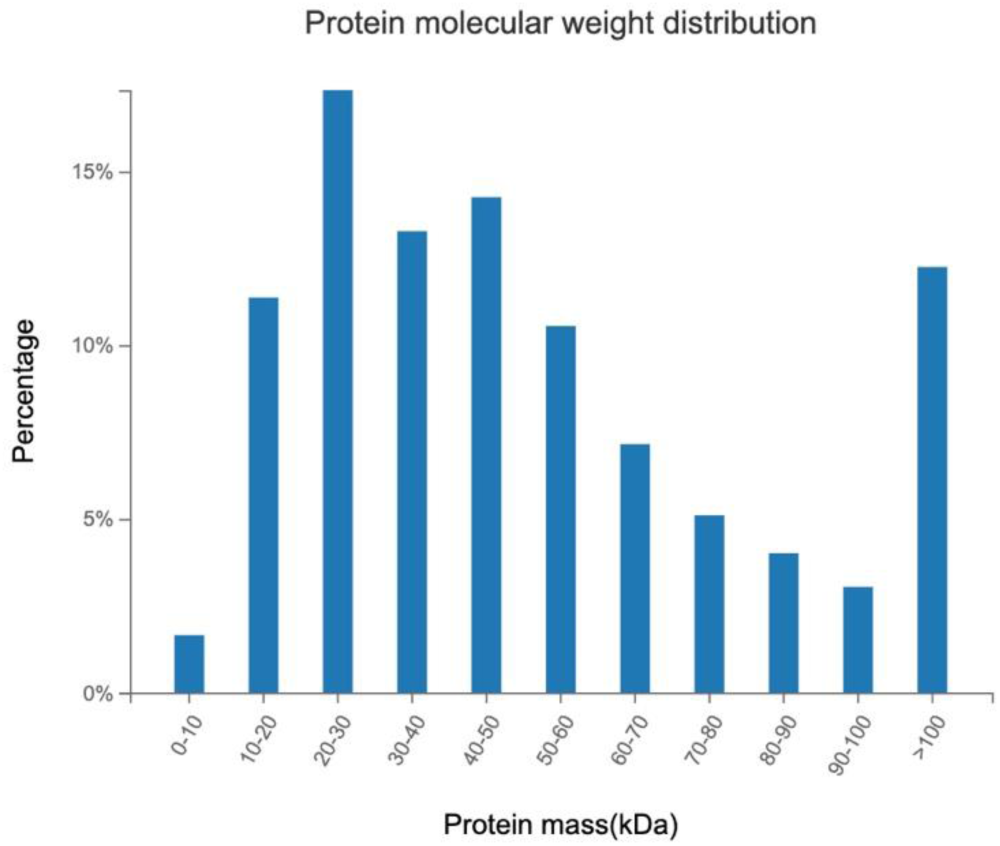
Protein molecular weight (kDa) distribution of proteins sequenced in seminal extracellular samples from high (n = 6) and reduced fertility (n = 6) animals by DIA Mass Spectrometry Detection.

**Supplementary Figure 3.**
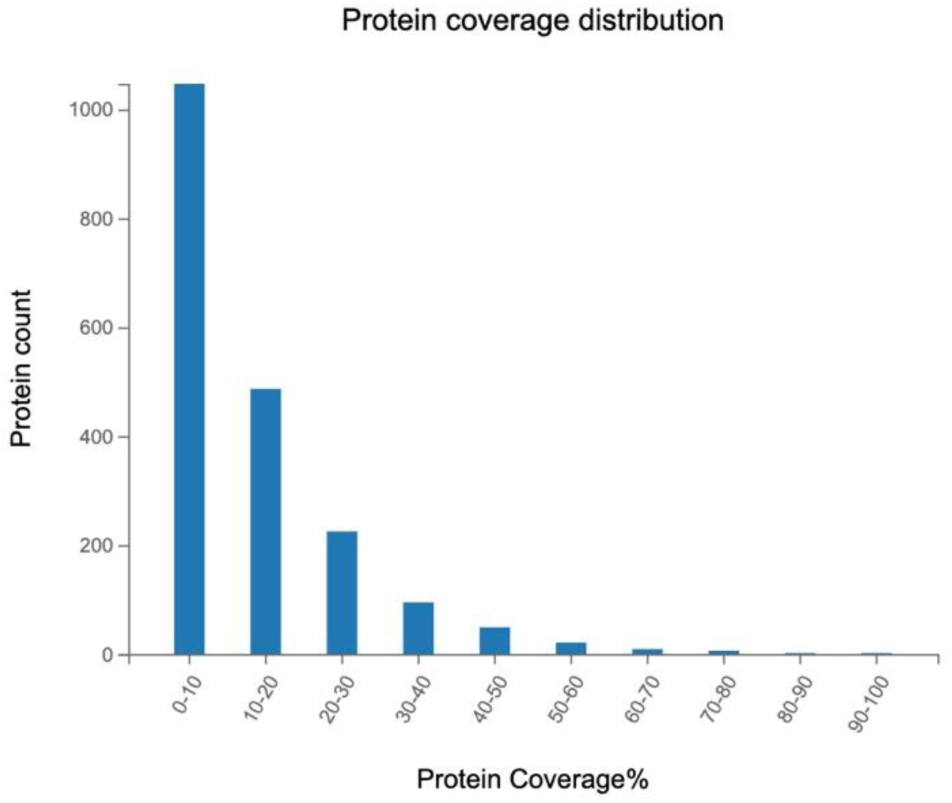
Protein coverage distribution (%) of proteins sequenced in seminal extracellular samples from high (n = 6) and reduced fertility (n = 6) animals by DIA Mass Spectrometry Detection.

---

**Supplementary Table 1.** Total number of peptides and proteins sequenced in seminal extracellular vesicles from high (HF, n = 6) and reduced fertility (RF, n = 6) animals by DIA Mass Spectrometry Detection.

| Sample Name | Peptides | Proteins Identified |
| --- | --- | --- |
| HF_EVs_1 | 7838 | 1756 |
| HF_EVs_2 | 5156 | 1283 |
| HF_EVs_3 | 8740 | 1783 |
| HF_EVs_4 | 5671 | 1416 |
| HF_EVs_5 | 6735 | 1596 |
| HF_EVs_6 | 6606 | 1523 |
| RF_EVs_1 | 5815 | 1446 |
| RF_EVs_2 | 6116 | 1447 |
| RF_EVs_3 | 7259 | 1576 |
| RF_EVs_4 | 6018 | 1444 |
| RF_EVs_5 | 5593 | 1454 |
| RF_EVs_6 | 5929 | 1431 |

**Supplementary Table 2.**
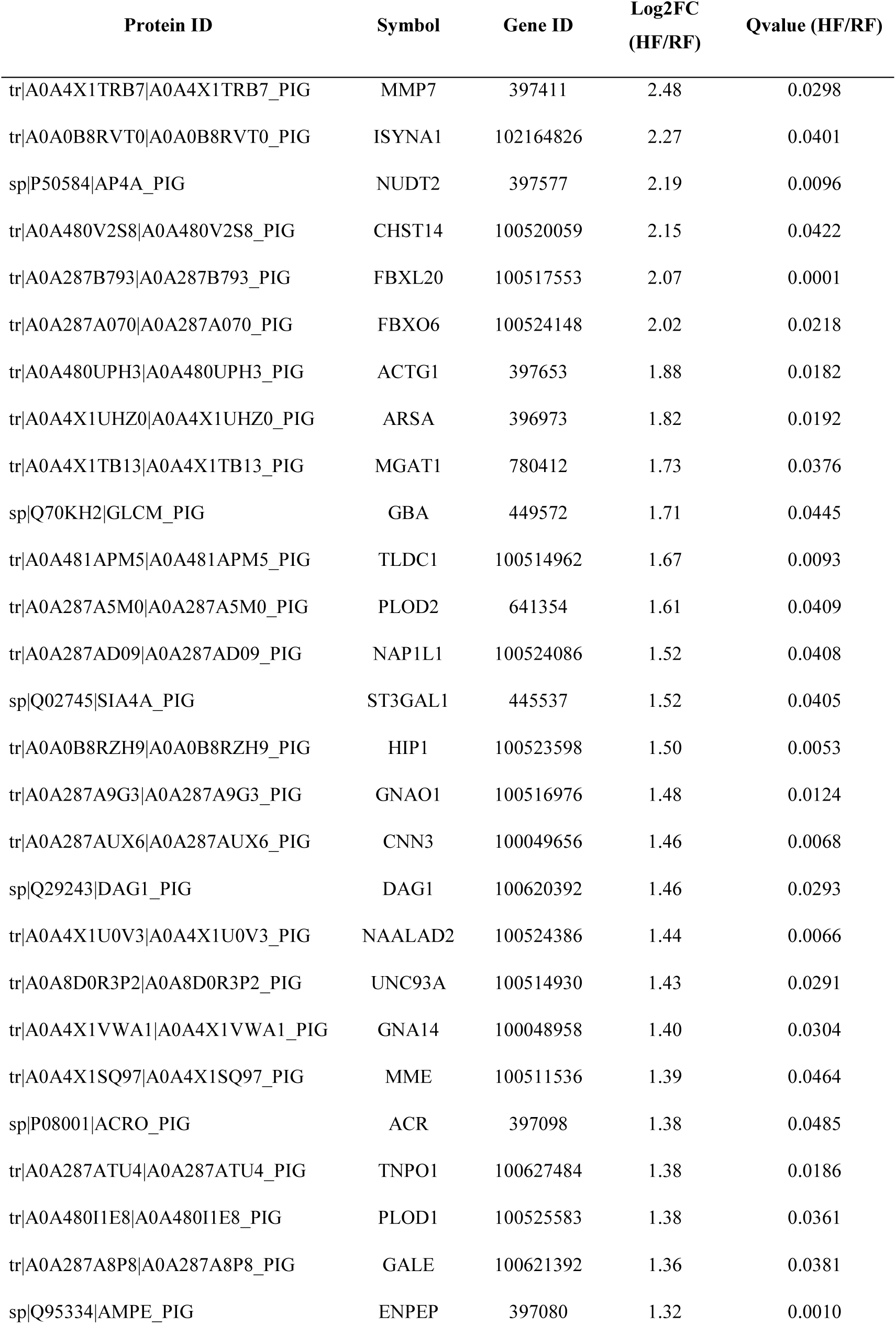
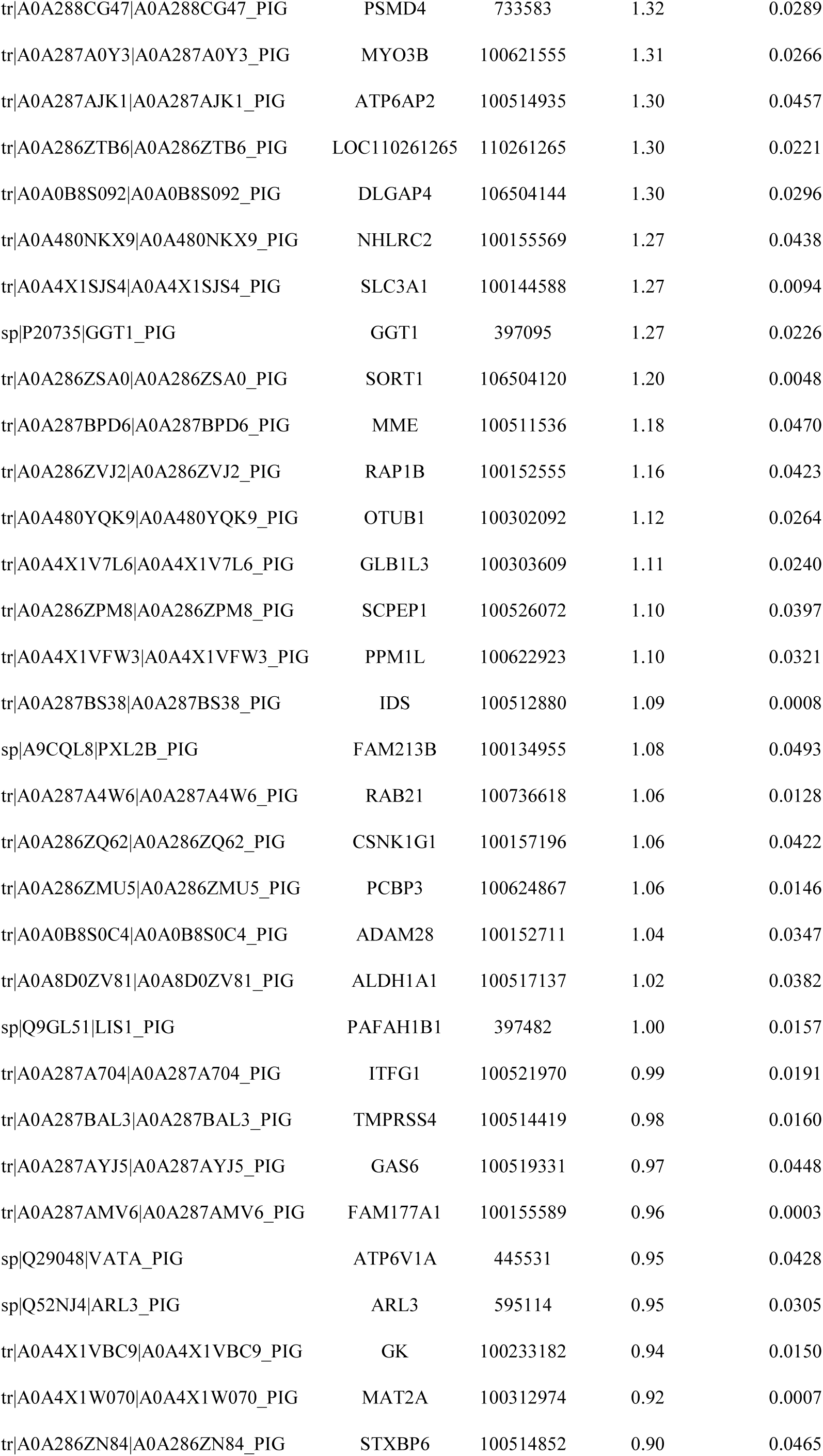
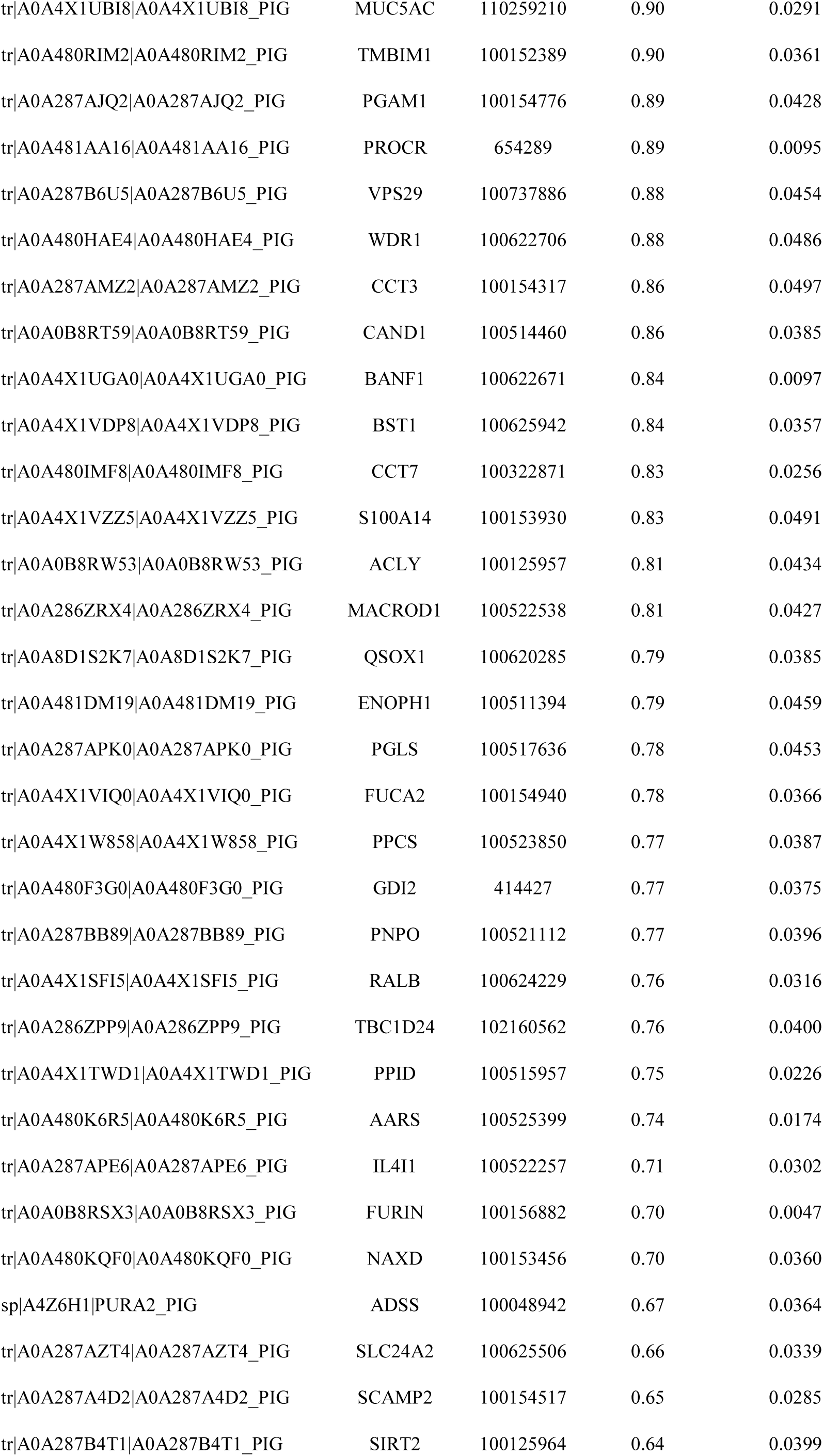
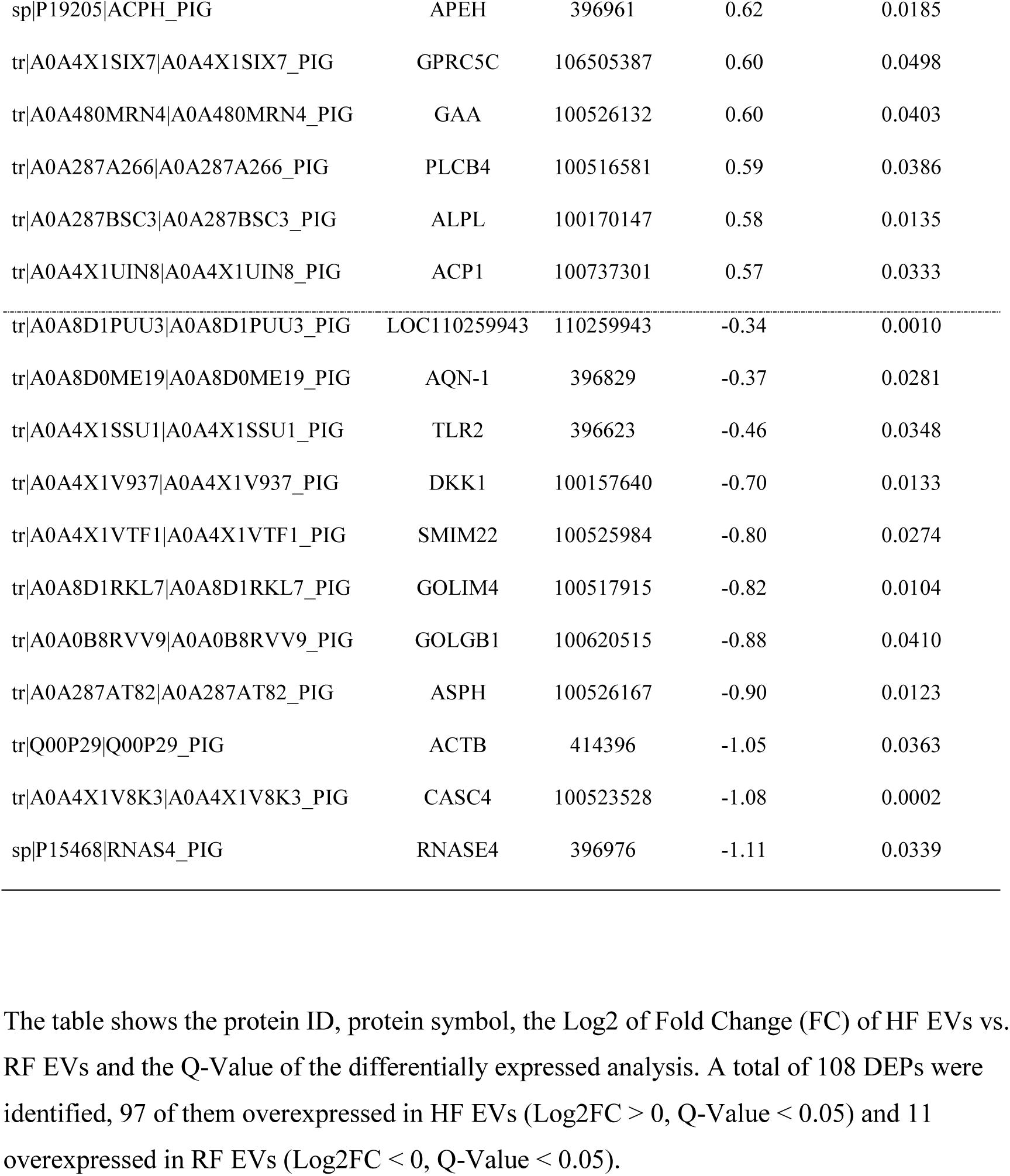
Total differentially expressed proteins (DEPs) identified in seminal extracellular vesicles (EVs) samples from high (HF, n = 6) and reduced fertility (RF, n = 6) animals.

**Supplementary Table 3.** Alignments of the reads after quality control and processing with *Sus scrofa* reference genome of seminal extracellular vesicles (EVs), free transcripts in seminal plasma (SP) and inside sperm cells (SPZ) from high (HF) and reduced fertility animals (RF). Values are expressed in mean, standard deviation (SD) and standard error of the mean (SEM).

| Sample | Total Clean Reads<br>(M) | Total Mapping (%) |  |  |  |
| --- | --- | --- | --- | --- | --- |
| RF_EVs_1 | 22.8 | 78.1 |  |  |  |
| RF_EVs_2 | 25.5 | 56.6 |  |  |  |
| RF_EVs_3 | 25.1 | 60.3 |  |  |  |
| RF_EVs_4 | 24.6 | 58.3 |  |  |  |
| RF_EVs_5 | 23.2 | 46.8 |  |  |  |
| RF_EVs_6 | 25.3 | 53.8 |  |  |  |
| HF_EVs_7 | 25.0 | 59.7 |  |  |  |
| HF_EVs_8 | 24.6 | 62.4 |  |  |  |
| HF_EVs_9 | 25.0 | 44.9 |  |  |  |
| HF_EVs_10 | 25.4 | 55.9 |  |  |  |
| HF_EVs_11 | 24.8 | 45.6 | Mean | SD | SEM |
| HF_EVs_12 | 25.0 | 38.8 | 55.1 | 10.3 | 3.0 |
| RF_SP_1 | 22.1 | 85.3 |  |  |  |
| RF_SP_2 | 21.9 | 57.1 |  |  |  |
| RF_SP_3 | 22.9 | 74.3 |  |  |  |
| RF_SP_4 | 22.9 | 65.7 |  |  |  |
| RF_SP_5 | 26.2 | 57.3 |  |  |  |
| RF_SP_6 | 24.8 | 55.7 |  |  |  |
| HF_SP_7 | 23.8 | 72.3 |  |  |  |
| HF_SP_8 | 23.5 | 53.8 |  |  |  |
| HF_SP_9 | 26.2 | 46.1 |  |  |  |
| HF_SP_10 | 25.5 | 46.1 |  |  |  |
| HF_SP_11 | 23.9 | 45.4 | Mean | SD | SEM |
| HF_SP_12 | 26.5 | 49.6 | 59.1 | 12.8 | 3.7 |
| RF_SPZ_1 | 24.8 | 95.9 |  |  |  |
| RF_SPZ_2 | 24.2 | 96.2 |  |  |  |
| RF_SPZ_3 | 22.0 | 96.4 |  |  |  |
| RF_SPZ_4 | 22.0 | 97.3 |  |  |  |
| RF_SPZ_5 | 23.2 | 94.9 |  |  |  |
| RF_SPZ_6 | 22.1 | 96.9 |  |  |  |
| HF_SPZ_7 | 25.3 | 95.9 |  |  |  |
| HF_SPZ_8 | 24.4 | 93.6 |  |  |  |
| HF_SPZ_9 | 26.2 | 92.7 |  |  |  |
| HF_SPZ_10 | 25.5 | 94.7 |  |  |  |
| HF_SPZ_11 | 24.6 | 93.2 | **Mean** | **SD** | **SEM** |
| HF_SPZ_12 | 23.7 | 95.5 | 95.3 | 1.5 | 0.4 |

**Supplementary Table 4.**
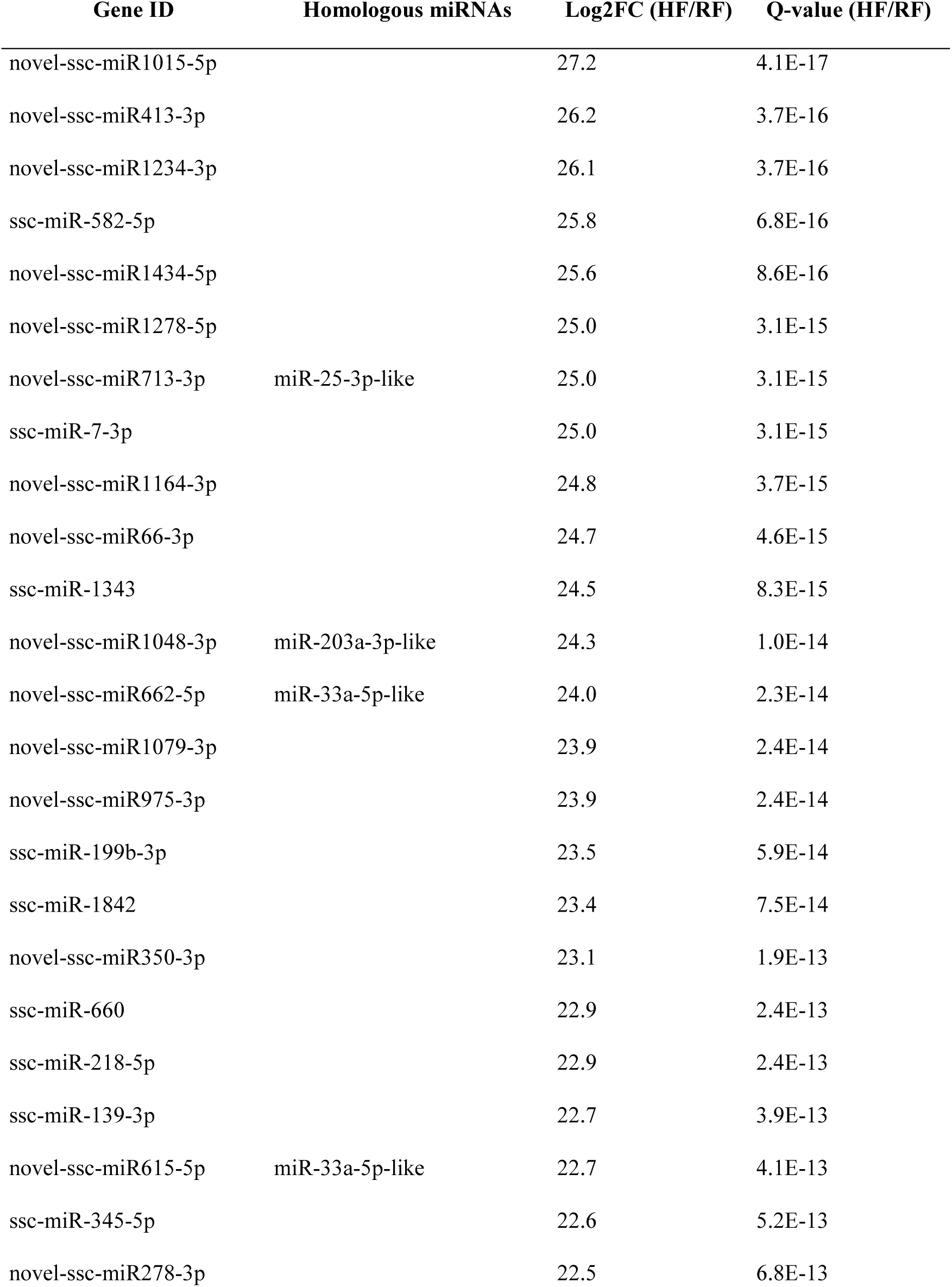
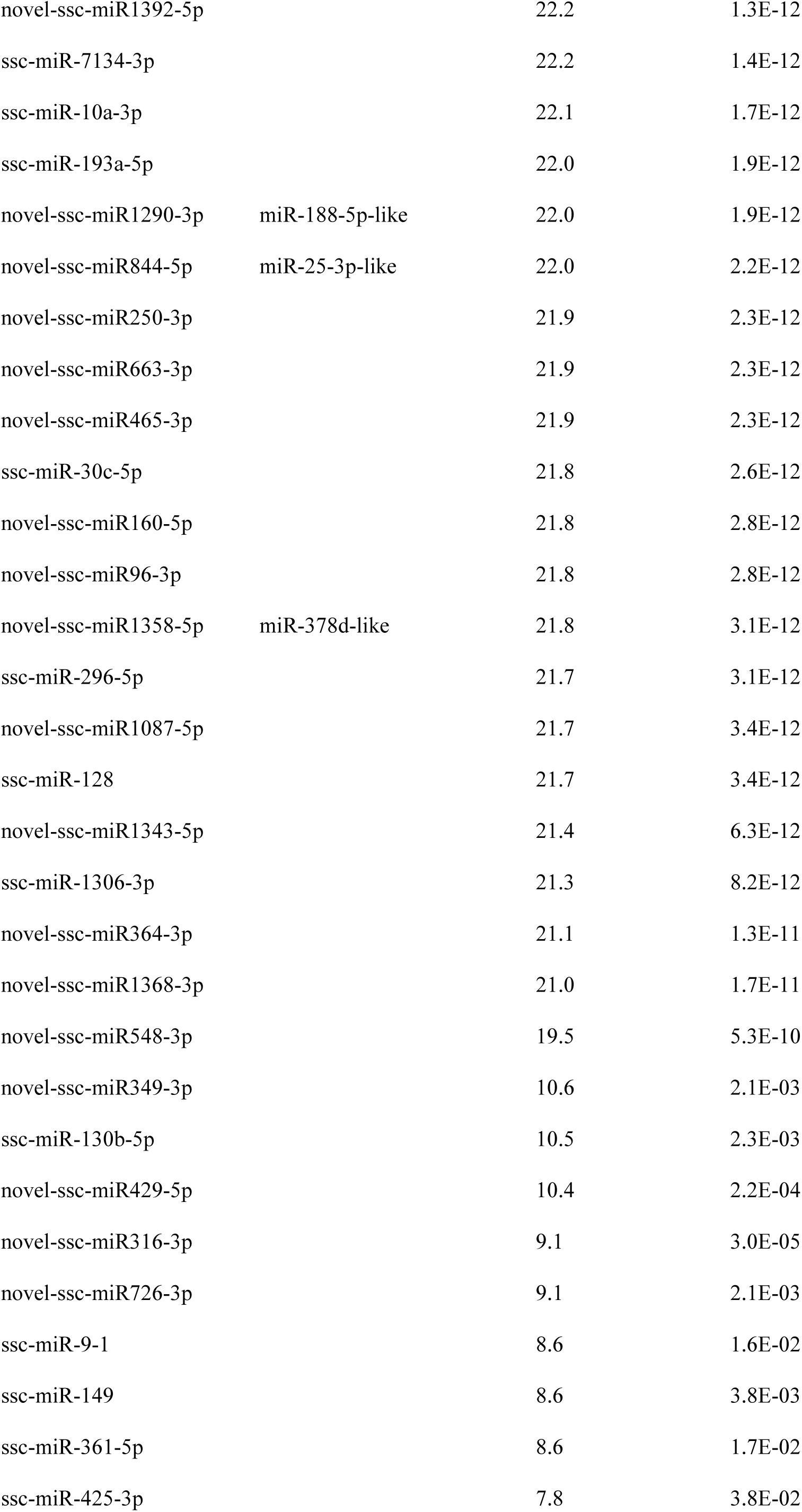
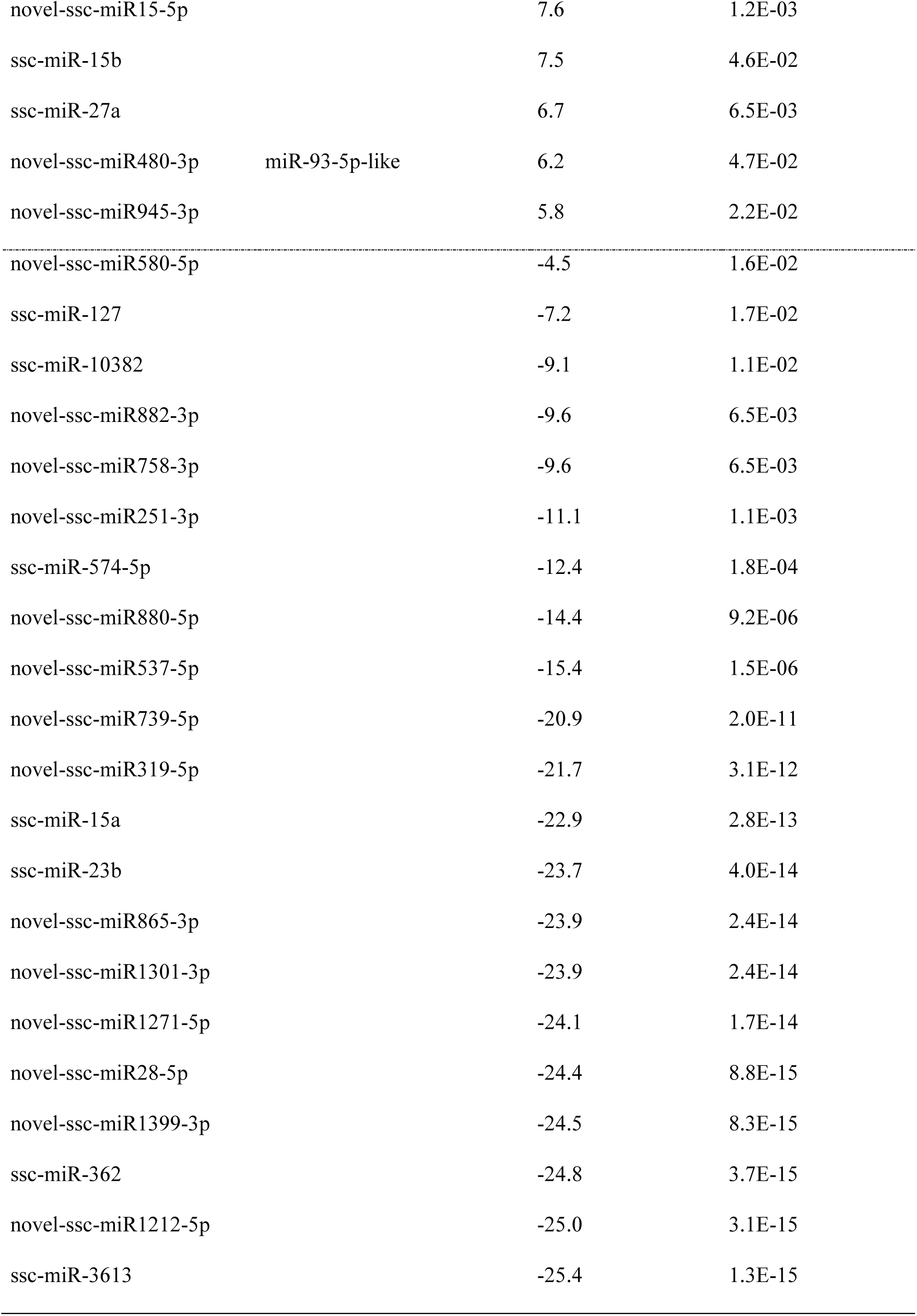
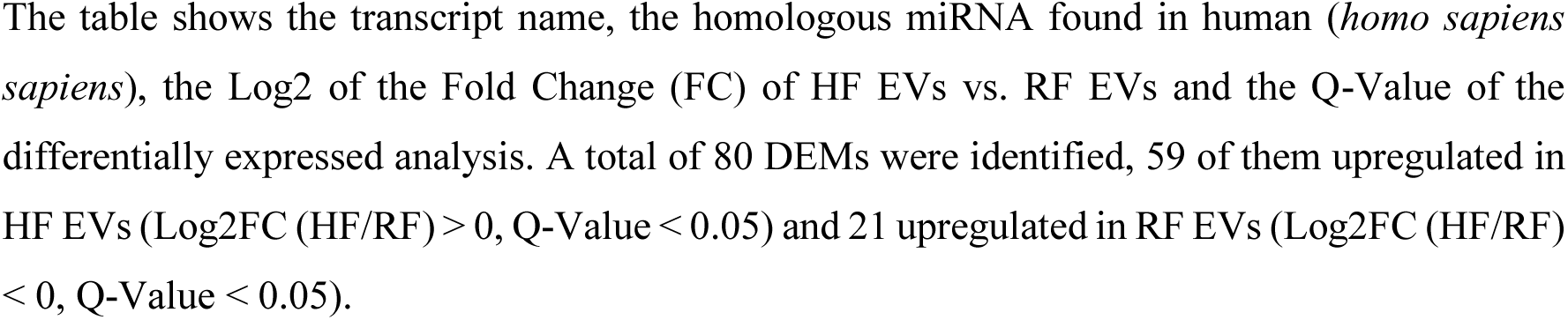
Total differentially expressed miRNAs (DEMs) identified in seminal extracellular vesicles (EVs) samples from high (HF, n = 6) and reduced fertility (RF, n = 6) animals.

| Gene ID | Homologous miRNAs | Log2FC (HF/RF) | Q-value (HF/RF) |
| --- | --- | --- | --- |
| novel-ssc-miR1015-5p |  | 27.2 | 4.1E-17 |
| novel-ssc-miR413-3p |  | 26.2 | 3.7E-16 |
| novel-ssc-miR1234-3p |  | 26.1 | 3.7E-16 |
| ssc-miR-582-5p |  | 25.8 | 6.8E-16 |
| novel-ssc-miR1434-5p |  | 25.6 | 8.6E-16 |
| novel-ssc-miR1278-5p |  | 25.0 | 3.1E-15 |
| novel-ssc-miR713-3p | miR-25-3p-like | 25.0 | 3.1E-15 |
| ssc-miR-7-3p |  | 25.0 | 3.1E-15 |
| novel-ssc-miR1164-3p |  | 24.8 | 3.7E-15 |
| novel-ssc-miR66-3p |  | 24.7 | 4.6E-15 |
| ssc-miR-1343 |  | 24.5 | 8.3E-15 |
| novel-ssc-miR1048-3p | miR-203a-3p-like | 24.3 | 1.0E-14 |
| novel-ssc-miR662-5p | miR-33a-5p-like | 24.0 | 2.3E-14 |
| novel-ssc-miR1079-3p |  | 23.9 | 2.4E-14 |
| novel-ssc-miR975-3p |  | 23.9 | 2.4E-14 |
| ssc-miR-199b-3p |  | 23.5 | 5.9E-14 |
| ssc-miR-1842 |  | 23.4 | 7.5E-14 |
| novel-ssc-miR350-3p |  | 23.1 | 1.9E-13 |
| ssc-miR-660 |  | 22.9 | 2.4E-13 |
| ssc-miR-218-5p |  | 22.9 | 2.4E-13 |
| ssc-miR-139-3p |  | 22.7 | 3.9E-13 |
| novel-ssc-miR615-5p | miR-33a-5p-like | 22.7 | 4.1E-13 |
| ssc-miR-345-5p |  | 22.6 | 5.2E-13 |
| novel-ssc-miR278-3p |  | 22.5 | 6.8E-13 |
| novel-ssc-miR1392-5p |  | 22.2 | 1.3E-12 |
| ssc-miR-7134-3p |  | 22.2 | 1.4E-12 |
| ssc-miR-10a-3p |  | 22.1 | 1.7E-12 |
| ssc-miR-193a-5p |  | 22.0 | 1.9E-12 |
| novel-ssc-miR1290-3p | miR-188-5p-like | 22.0 | 1.9E-12 |
| novel-ssc-miR844-5p | miR-25-3p-like | 22.0 | 2.2E-12 |
| novel-ssc-miR250-3p |  | 21.9 | 2.3E-12 |
| novel-ssc-miR663-3p |  | 21.9 | 2.3E-12 |
| novel-ssc-miR465-3p |  | 21.9 | 2.3E-12 |
| ssc-miR-30c-5p |  | 21.8 | 2.6E-12 |
| novel-ssc-miR160-5p |  | 21.8 | 2.8E-12 |
| novel-ssc-miR96-3p |  | 21.8 | 2.8E-12 |
| novel-ssc-miR1358-5p | miR-378d-like | 21.8 | 3.1E-12 |
| ssc-miR-296-5p |  | 21.7 | 3.1E-12 |
| novel-ssc-miR1087-5p |  | 21.7 | 3.4E-12 |
| ssc-miR-128 |  | 21.7 | 3.4E-12 |
| novel-ssc-miR1343-5p |  | 21.4 | 6.3E-12 |
| ssc-miR-1306-3p |  | 21.3 | 8.2E-12 |
| novel-ssc-miR364-3p |  | 21.1 | 1.3E-11 |
| novel-ssc-miR1368-3p |  | 21.0 | 1.7E-11 |
| novel-ssc-miR548-3p |  | 19.5 | 5.3E-10 |
| novel-ssc-miR349-3p |  | 10.6 | 2.1E-03 |
| ssc-miR-130b-5p |  | 10.5 | 2.3E-03 |
| novel-ssc-miR429-5p |  | 10.4 | 2.2E-04 |
| novel-ssc-miR316-3p |  | 9.1 | 3.0E-05 |
| novel-ssc-miR726-3p |  | 9.1 | 2.1E-03 |
| ssc-miR-9-1 |  | 8.6 | 1.6E-02 |
| ssc-miR-149 |  | 8.6 | 3.8E-03 |
| ssc-miR-361-5p |  | 8.6 | 1.7E-02 |
| ssc-miR-425-3p |  | 7.8 | 3.8E-02 |
| novel-ssc-miR15-5p |  | 7.6 | 1.2E-03 |
| ssc-miR-15b |  | 7.5 | 4.6E-02 |
| ssc-miR-27a |  | 6.7 | 6.5E-03 |
| novel-ssc-miR480-3p | miR-93-5p-like | 6.2 | 4.7E-02 |
| novel-ssc-miR945-3p |  | 5.8 | 2.2E-02 |
| novel-ssc-miR580-5p |  | -4.5 | 1.6E-02 |
| ssc-miR-127 |  | -7.2 | 1.7E-02 |
| ssc-miR-10382 |  | -9.1 | 1.1E-02 |
| novel-ssc-miR882-3p |  | -9.6 | 6.5E-03 |
| novel-ssc-miR758-3p |  | -9.6 | 6.5E-03 |
| novel-ssc-miR251-3p |  | -11.1 | 1.1E-03 |
| ssc-miR-574-5p |  | -12.4 | 1.8E-04 |
| novel-ssc-miR880-5p |  | -14.4 | 9.2E-06 |
| novel-ssc-miR537-5p |  | -15.4 | 1.5E-06 |
| novel-ssc-miR739-5p |  | -20.9 | 2.0E-11 |
| novel-ssc-miR319-5p |  | -21.7 | 3.1E-12 |
| ssc-miR-15a |  | -22.9 | 2.8E-13 |
| ssc-miR-23b |  | -23.7 | 4.0E-14 |
| novel-ssc-miR865-3p |  | -23.9 | 2.4E-14 |
| novel-ssc-miR1301-3p |  | -23.9 | 2.4E-14 |
| novel-ssc-miR1271-5p |  | -24.1 | 1.7E-14 |
| novel-ssc-miR28-5p |  | -24.4 | 8.8E-15 |
| novel-ssc-miR1399-3p |  | -24.5 | 8.3E-15 |
| ssc-miR-362 |  | -24.8 | 3.7E-15 |
| novel-ssc-miR1212-5p |  | -25.0 | 3.1E-15 |
| ssc-miR-3613 |  | -25.4 | 1.3E-15 |

The table shows the transcript name, the homologous miRNA found in human (*homo sapiens*), the Log2 of the Fold Change (FC) of HF EVs vs. RF EVs and the Q-Value of the differentially expressed analysis. A total of 80 DEMs were identified, 59 of them upregulated in HF EVs ( $\text{Log2FC (HF/RF)} > 0$ , $\text{Q-Value} < 0.05$ ) and 21 upregulated in RF EVs ( $\text{Log2FC (HF/RF)} < 0$ , $\text{Q-Value} < 0.05$ ).

**Supplementary Table 5.**
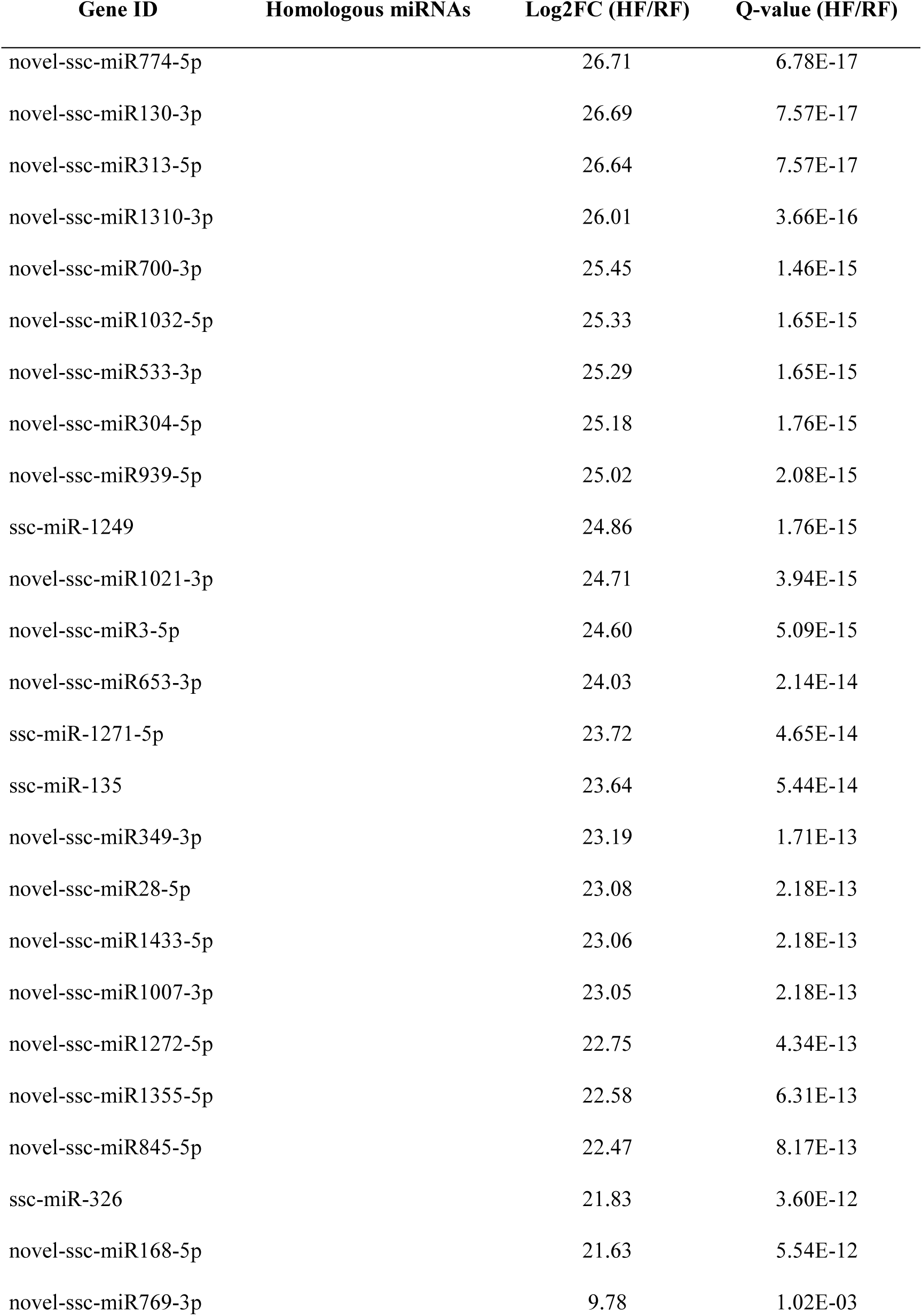
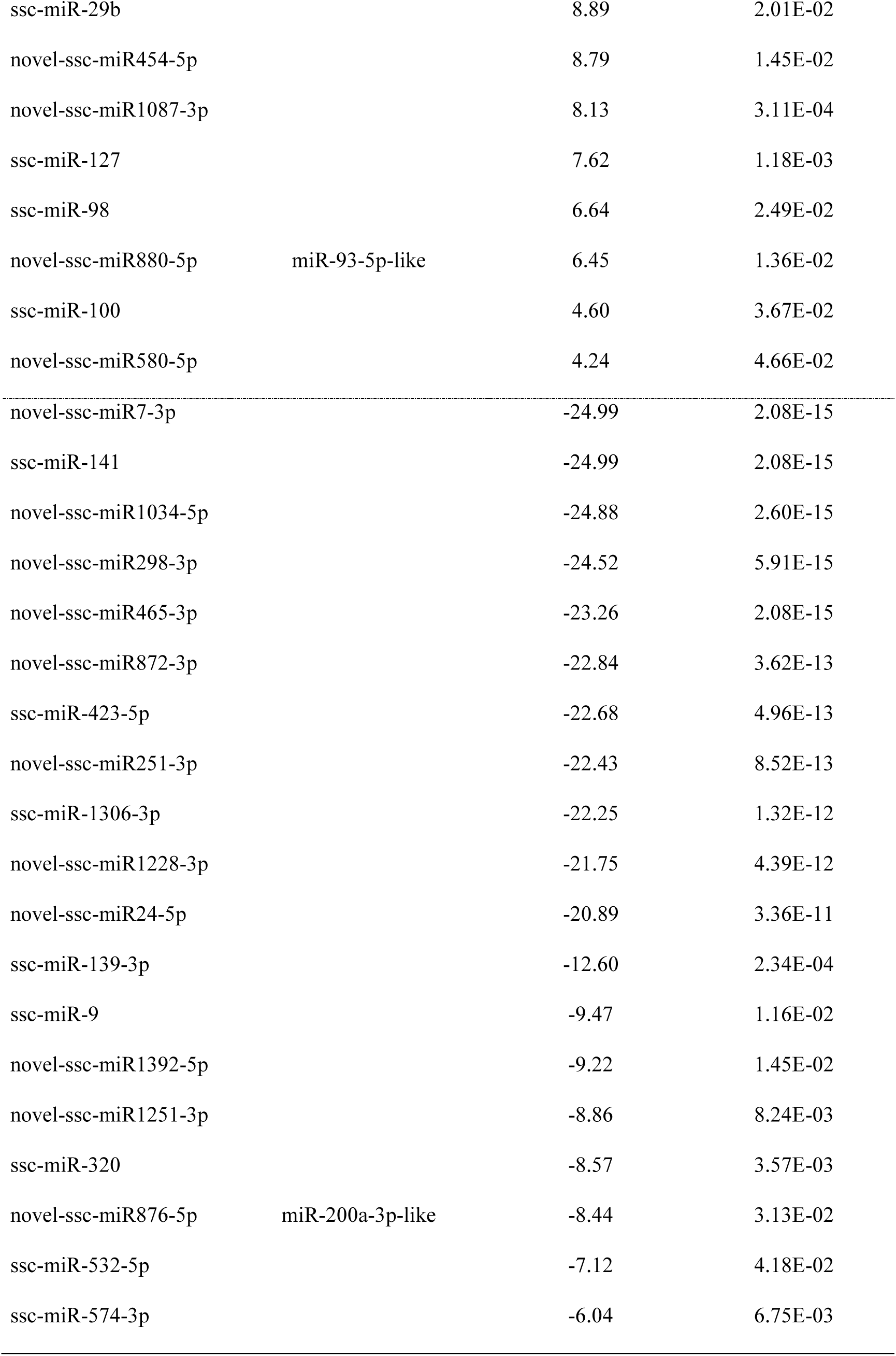
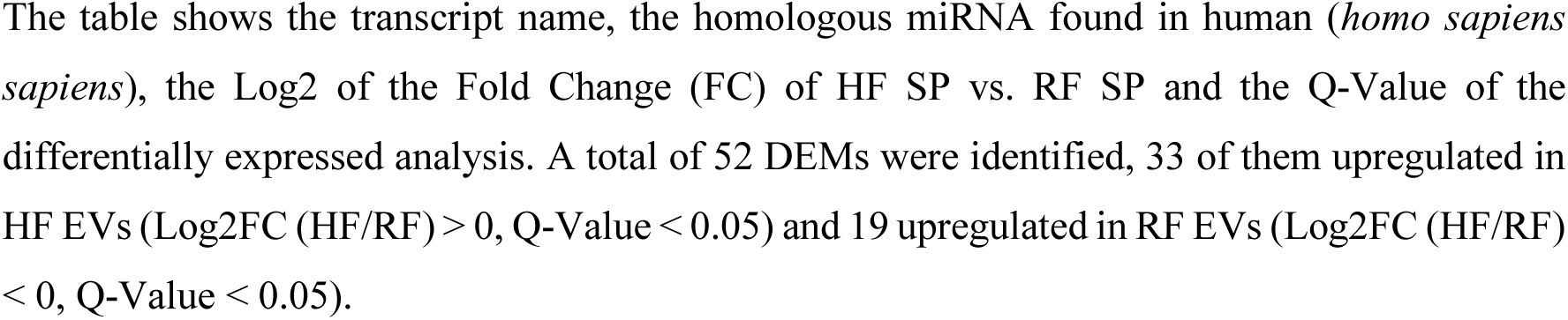
Total differentially expressed miRNAs (DEMs) identified in the seminal plasma (SP) from high (HF, n = 6) and reduced fertility (RF, n = 6) animals.

| Gene ID | Homologous miRNAs | Log2FC (HF/RF) | Q-value (HF/RF) |
| --- | --- | --- | --- |
| novel-ssc-miR774-5p |  | 26.71 | 6.78E-17 |
| novel-ssc-miR130-3p |  | 26.69 | 7.57E-17 |
| novel-ssc-miR313-5p |  | 26.64 | 7.57E-17 |
| novel-ssc-miR1310-3p |  | 26.01 | 3.66E-16 |
| novel-ssc-miR700-3p |  | 25.45 | 1.46E-15 |
| novel-ssc-miR1032-5p |  | 25.33 | 1.65E-15 |
| novel-ssc-miR533-3p |  | 25.29 | 1.65E-15 |
| novel-ssc-miR304-5p |  | 25.18 | 1.76E-15 |
| novel-ssc-miR939-5p |  | 25.02 | 2.08E-15 |
| ssc-miR-1249 |  | 24.86 | 1.76E-15 |
| novel-ssc-miR1021-3p |  | 24.71 | 3.94E-15 |
| novel-ssc-miR3-5p |  | 24.60 | 5.09E-15 |
| novel-ssc-miR653-3p |  | 24.03 | 2.14E-14 |
| ssc-miR-1271-5p |  | 23.72 | 4.65E-14 |
| ssc-miR-135 |  | 23.64 | 5.44E-14 |
| novel-ssc-miR349-3p |  | 23.19 | 1.71E-13 |
| novel-ssc-miR28-5p |  | 23.08 | 2.18E-13 |
| novel-ssc-miR1433-5p |  | 23.06 | 2.18E-13 |
| novel-ssc-miR1007-3p |  | 23.05 | 2.18E-13 |
| novel-ssc-miR1272-5p |  | 22.75 | 4.34E-13 |
| novel-ssc-miR1355-5p |  | 22.58 | 6.31E-13 |
| novel-ssc-miR845-5p |  | 22.47 | 8.17E-13 |
| ssc-miR-326 |  | 21.83 | 3.60E-12 |
| novel-ssc-miR168-5p |  | 21.63 | 5.54E-12 |
| novel-ssc-miR769-3p |  | 9.78 | 1.02E-03 |
| ssc-miR-29b |  | 8.89 | 2.01E-02 |
| novel-ssc-miR454-5p |  | 8.79 | 1.45E-02 |
| novel-ssc-miR1087-3p |  | 8.13 | 3.11E-04 |
| ssc-miR-127 |  | 7.62 | 1.18E-03 |
| ssc-miR-98 |  | 6.64 | 2.49E-02 |
| novel-ssc-miR880-5p | miR-93-5p-like | 6.45 | 1.36E-02 |
| ssc-miR-100 |  | 4.60 | 3.67E-02 |
| novel-ssc-miR580-5p |  | 4.24 | 4.66E-02 |
| novel-ssc-miR7-3p |  | -24.99 | 2.08E-15 |
| ssc-miR-141 |  | -24.99 | 2.08E-15 |
| novel-ssc-miR1034-5p |  | -24.88 | 2.60E-15 |
| novel-ssc-miR298-3p |  | -24.52 | 5.91E-15 |
| novel-ssc-miR465-3p |  | -23.26 | 2.08E-15 |
| novel-ssc-miR872-3p |  | -22.84 | 3.62E-13 |
| ssc-miR-423-5p |  | -22.68 | 4.96E-13 |
| novel-ssc-miR251-3p |  | -22.43 | 8.52E-13 |
| ssc-miR-1306-3p |  | -22.25 | 1.32E-12 |
| novel-ssc-miR1228-3p |  | -21.75 | 4.39E-12 |
| novel-ssc-miR24-5p |  | -20.89 | 3.36E-11 |
| ssc-miR-139-3p |  | -12.60 | 2.34E-04 |
| ssc-miR-9 |  | -9.47 | 1.16E-02 |
| novel-ssc-miR1392-5p |  | -9.22 | 1.45E-02 |
| novel-ssc-miR1251-3p |  | -8.86 | 8.24E-03 |
| ssc-miR-320 |  | -8.57 | 3.57E-03 |
| novel-ssc-miR876-5p | miR-200a-3p-like | -8.44 | 3.13E-02 |
| ssc-miR-532-5p |  | -7.12 | 4.18E-02 |
| ssc-miR-574-3p |  | -6.04 | 6.75E-03 |

The table shows the transcript name, the homologous miRNA found in human (*homo sapiens*), the Log2 of the Fold Change (FC) of HF SP vs. RF SP and the Q-Value of the differentially expressed analysis. A total of 52 DEMs were identified, 33 of them upregulated in HF EVs ( $\text{Log2FC (HF/RF)} > 0$ , $\text{Q-Value} < 0.05$ ) and 19 upregulated in RF EVs ( $\text{Log2FC (HF/RF)} < 0$ , $\text{Q-Value} < 0.05$ ).

**Supplementary Table 6.** Total differentially expressed miRNAs (DEMs) identified in spermatozoa (SPZ) from high (HF, n = 6) and reduced fertility (RF, n = 6) animals.

| Gene ID | Log2FC (HF/RF) | Q-value (HF/RF) |
| --- | --- | --- |
| novel-ssc-miR1250-5p | 2.190 | 0.048 |
| novel-ssc-miR1416-5p | 1.980 | 0.048 |
| novel-ssc-miR663-3p | 1.814 | 0.048 |

The table shows the transcript name, the Log2 of the Fold Change (FC) of HF SPZ vs. RF SPZ and the Q-Value of the differentially expressed analysis. A total of 3 novel DEMs were identified, all of them upregulated in HF EVs (Log2FC (HF/RF) > 0, Q-Value < 0.05).

